# MCM10 targets CMG dimers via a conserved mechanism for synchronized helicase activation

**DOI:** 10.64898/2026.05.15.725364

**Authors:** Ella Taylor-Cross, Morgan L. Jones, Johann J. Roske, Emma E. Fletcher, Isaac X.R. Chan, Joseph T. P. Yeeles

## Abstract

Activation of the CMG (CDC45–MCM–GINS) helicase by MCM10 is a central unresolved step in DNA replication. DNA is melted within a dimeric CMG complex and one strand is expelled from each topologically closed MCM ring. How MCM10 drives this process is unclear owing to a lack of structural information. Here, by determining structures of yeast CMG-Mcm10 complexes, and of human CMG-Pol ε dimers assembled by DONSON and bound to MCM10 and the helicase activator RECQL4, we reveal a conserved mechanism of CMG helicase activation in eukaryotes. MCM10 targets CMG dimers through highly conserved and species-specific interactions, that in human also involve RECQL4. Our data indicate this arrangement allows MCM10 to stimulate DNA unwinding by CMG and to utilise the associated conformational changes to drive single-stranded DNA ejection between MCM2 and MCM5. This mechanism provides an explanation for synchronized activation of two CMG helicases at origins of bidirectional replication.

## Introduction

To duplicate large eukaryotic genomes DNA replication is initiated at multiple sites called origins. This process, collectively termed origin firing, involves the chromatin loading of the MCM2-7 complex as a double hexamer (DH) and its subsequent conversion into two active CMG (CDC45-MCM-GINS) helicases that each contain a single MCM2-7 heterohexamer bound by CDC45 and GINS. The MCM DH is loaded in G1 phase by ORC, CDC6 and CDT1 and consists of two topologically closed MCM2-7 rings - bound around double stranded (ds) DNA - that are interlocked by interactions between zinc-finger (ZnF) domains present in the N-terminal domains of each MCM subunit (Remus et al. 2009; Evrin et al. 2009; Yang et al. 2024; Wells et al. 2025; Weissmann et al. 2024). During MCM loading dsDNA is inserted into a central pore in each MCM ring through a gap that is formed between the MCM2 and MCM5 subunits (Samel et al. 2014). During CMG assembly - which occurs in S phase and is orchestrated by several ‘firing factor’ proteins - approximately 0.6-0.7 turns of DNA are untwisted per CMG (Douglas et al. 2018) but the MCM rings remain topologically closed and partially interlocked and both DNA strands remain within the central pore (Lewis et al. 2022). In the final stage of origin firing, CMG activation, dsDNA is further unwound, the lagging-strand template is expulsed from the central pore of each CMG, and the two CMGs separate and translocate past one another to establish replication forks (Douglas, et al. 2018; Georgescu et al. 2017). In *S. cerevisiae*, these structural transitions strictly require the essential and highly conserved protein Mcm10 (Yeeles et al. 2015; Douglas, et al. 2018; van Deursen et al. 2012; Watase, Takisawa, and Kanemaki 2012; Quan et al. 2015). By contrast, Mcm-10 is not essential in *C. elegans* (Xia et al. 2023) and DNA synthesis persists in Xenopus egg extracts depleted of MCM10 (Chadha et al. 2016), indicating that CMGs can be activated independently of MCM10 in some eukaryotes and raising questions about the extent to which MCM10 functions via a conserved mechanism.

The defining structural feature of MCM10 is a conserved OB-fold zinc-finger domain (OBZnF) that displays a preference for binding single-stranded (ss)DNA (Robertson et al. 2008; Fien et al. 2004; Eisenberg et al. 2009; Warren et al. 2009). Its integrity is essential for yeast viability (Homesley et al. 2000; Das-Bradoo, Ricke, and Bielinsky 2006; Cook et al. 2003; Kanke et al. 2012). Notably, a recent study found that, although mutation of the Mcm10 OB-fold abolished ssDNA binding it had no effect on yeast CMG activation in an *in vitro* system (Henrikus et al. 2024). As such, the role of the OBZnF during CMG activation is unresolved. N-terminal of the OBZnF, MCM10 contains a domain that is proposed to mediate homo-oligomerization (Du et al. 2013) and binding to the 9-1-1 clamp (Alver et al. 2014). Neither putative function is essential in *S. cerevisiae* because cells are viable when the first 150 amino acids of Mcm10 are deleted (Alver, et al. 2014). In contrast, truncation of the Mcm10 C-terminus impacts cell growth, likely because it contains binding sites for the Mcm2 and Mcm6 subunits of the CMG complex (Douglas and Diffley 2016). Accordingly, cross-linking mass spectrometry analysis of a CMG:Mcm10 complex revealed that Mcm10 crosslinked with Mcm2 and Mcm6 and the neighbouring subunits Mcm5 and Cdc45 (Mayle et al. 2019).

Despite extensive characterisation of MCM10 since its discovery 40-years ago (Nasmyth and Nurse 1981; Dumas et al. 1982; Maine, Sinha, and Tye 1984), the structural basis for CMG activation remains elusive. It is unclear how the lagging-strand ssDNA exits the MCM ring and if ssDNA binding by MCM10 is involved in this process, how MCM10 engages CMG, and how this engagement drives CMG activation. Efforts to elucidate a mechanism have been limited by a lack of any structures of MCM10 bound to CMG. Although a stoichiometric complex of *S. cerevisiae* CMG:Mcm10 was isolated after anion exchange chromatography, density corresponding to Mcm10 was not identified in cryo-EM maps (Mayle, et al. 2019). A recent cryo-EM analysis of *in vitro* CMG assembly and activation with purified yeast proteins identified density alongside the ZnF domain of Mcm2 (Henrikus, et al. 2024). It was hypothesised that this density was the Mcm10 OB-fold and that Mcm10 activated CMG by disrupting the CMG homodimerization interface to split the two CMG complexes. However, inspection of the published cryo-EM density map (Henrikus, et al. 2024) shows that the region assigned as Mcm10 is only visible at map contour levels where noise becomes prominent, is low resolution, and is devoid of the necessary features required to make a confident domain assignment (Supplementary Figure 1A, B). These studies highlight the considerable challenges in obtaining structures of functionally relevant CMG complexes containing MCM10, which will be essential to reveal how MCM10 activates CMG during eukaryotic DNA replication origin firing.

## Results

### Structure of S. cerevisiae Mcm10 bound to CMG

To examine how Mcm10 engages CMG, we incubated purified *S. cerevisiae* CMG, Mcm10 and forked DNA and isolated complexes by glycerol gradient sedimentation in the presence of the chemical crosslinkers glutaraldehyde and BS3 (Gradient fixation (GraFix)) (Baretić et al. 2020; Kastner et al. 2008). For clarity, from here on *S. cerevisiae* proteins are prefixed _Sc_ and *H. sapiens* proteins _Hs_ (e.g. _Sc_CMG and _Hs_CMG). After buffer exchange to remove glycerol samples were analysed by cryo-EM (Supplementary Fig. 1C-F and 2). Extensive data processing revealed several densities alongside the _Sc_Mcm2 and _Sc_Mcm6 subunits of _Sc_CMG that could be unambiguously modelled as C-terminal regions of _Sc_Mcm10 (amino acids (a.a.) 397-532) with the aid of AlphaFold3 structure predictions (Abramson et al. 2024) (Fig. 1A and Supplementary Fig. 2G-J). We were unable to recover density for the OBZnF of _Sc_Mcm10 (Supplementary Fig. 2A, E), indicating that it might not adopt a stable position on _Sc_CMG. We also considered that a ssDNA region of the forked DNA could sequester the OBZnF preventing it from binding _Sc_CMG. Cryo-EM analysis of an _Sc_CMG:Mcm10 complex assembled without DNA supported this hypothesis (Supplementary Fig. 3 and 4). Figure 1B and C shows that a globular density corresponding to the _Sc_Mcm10 OBZnF was positioned adjacent to the interface between the _Sc_Mcm2 and _Sc_Cdc45 subunits of _Sc_CMG. Comparable density was also observed in a subtly different position in another particle subset (Supplementary Fig. 4B-E, H-J). Moreover, we resolved to side-chain resolution an additional binding site between _Sc_Mcm10 and _Sc_CMG involving an α-helix of _Sc_Mcm10 (a.a. 111-136) immediately N-terminal of the OB-fold bound to a hydrophobic patch on _Sc_Cdc45 (Fig. 1C and Supplementary Fig. 4K, L). A short α-helix in the _Sc_Mcm10 N-terminus (a.a. 90-97) was also observed bound to the _Sc_GINS subunit Psf2 (Supplementary Fig. 4M). The _Sc_Mcm10:Cdc45 interaction anchors the OBZnF alongside _Sc_Mcm2 and _Sc_Cdc45, which explains why the corresponding cryo-EM density was recovered. In neither data set was density recovered for the putative N-terminal homodimerization domain of _Sc_Mcm10 (Fig. 1D).

**Fig. 1.**
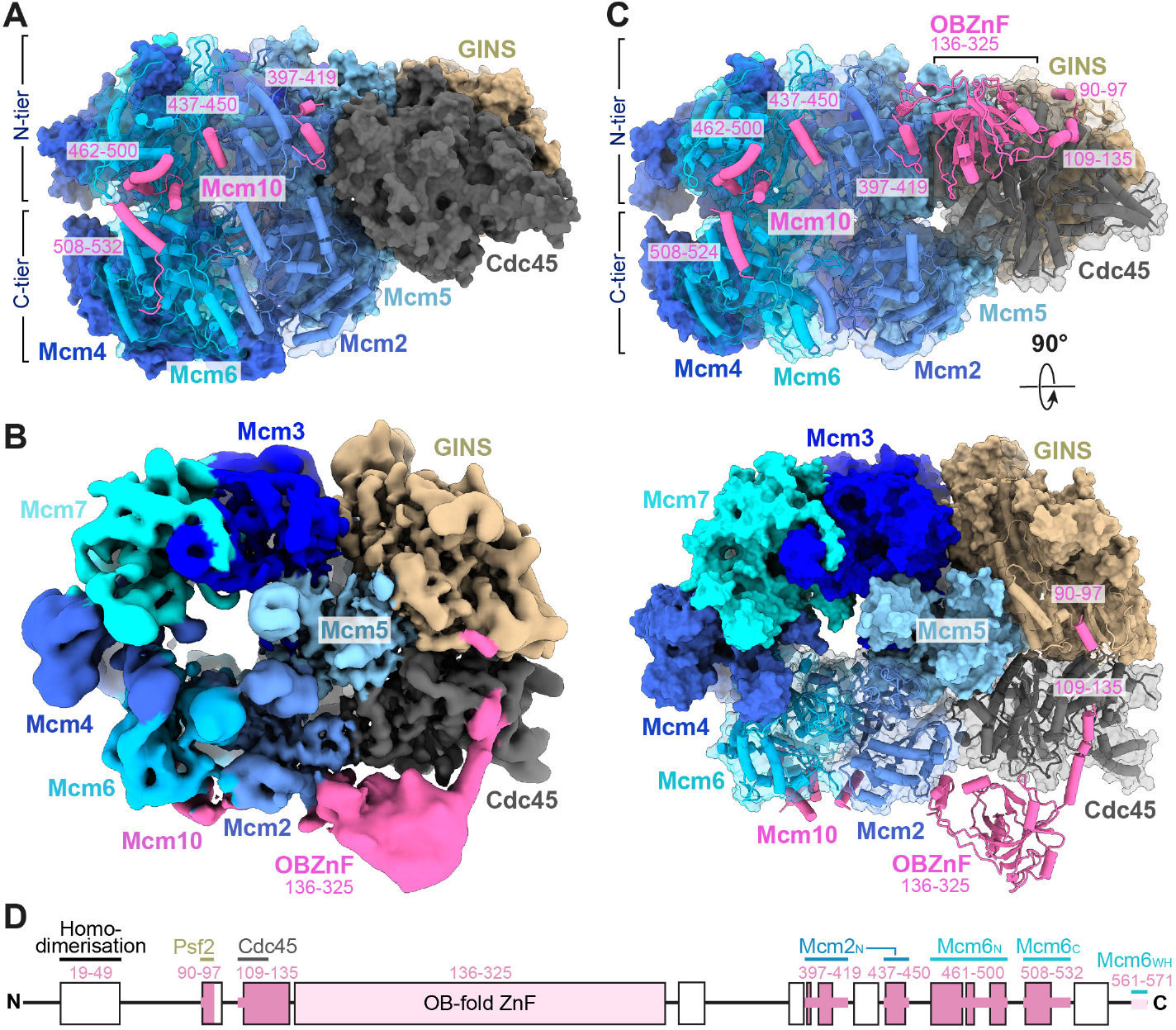
Cryo-EM structures of *S. cerevisiae* Mcm10 bound to CMG. (**A**) Atomic model of _Sc_CMG bound to replication fork DNA and _Sc_Mcm10. Mcm2 and Mcm6, which interact with Mcm10, are shown as cylinders/stubs with a transparent surface. (**B**) Cryo-EM density map of _Sc_Mcm10 bound to _Sc_CMG in the absence of DNA, coloured by subunit occupancy. (**C**) Atomic model of _Sc_CMG bound to Mcm10 in the absence of DNA. Mcm2, Mcm6 and Cdc45, which interact with Mcm10, are shown as cylinders/stubs with a transparent surface. (**D**) Primary structure diagram for _Sc_Mcm10. Boxes represent regions predicted to be structured by AlphaFold3 (Abramson, et al. 2024). Dark pink denotes regions that can be modelled from the cryo-EM structures, light pink denotes regions where AlphaFold3 models can be docked into cryo-EM density and regions not visible in cryo-EM structures are uncoloured. Regions of Mcm10 that interact with CMG subunits are labelled (N = N-tier, C = C-tier, WH = winged-helix).

C-terminal of the OBZnF, _Sc_Mcm10 (a.a. 397-419) binds to a hydrophobic patch on the _Sc_Mcm2 helical domain (HD) at its interface with _Sc_Cdc45 (Fig. 1A and Supplementary Fig. 2G). A short linker in _Sc_Mcm10, bridges to a second small hydrophobic binding site on the _Sc_Mcm2 HD near its interface with _Sc_Mcm6 (Fig. 1A and Supplementary Fig. 2H). This in turn is linked to a more extensive hydrophobic interaction site (_Sc_Mcm10 a.a. 461-491) situated between the _Sc_Mcm6 OB fold and HD (Fig. 1A and Supplementary Fig. 2I). _Sc_Mcm10 then attaches to the C-terminal AAA+ domain of _Sc_Mcm6 via a short α-helix (a.a. 510-522) (Fig. 1A and Supplementary Fig. 2J) while the final ∼40 a.a. of _Sc_Mcm10 are invisible. Binding studies have shown that _Sc_Mcm10 interacts with the winged-helix (WH) domain of _Sc_Mcm6 (Douglas and Diffley 2016). Although this interaction is not visualised in either structure, low resolution density was recovered alongside the _Sc_Mcm6 ATPase and the last modelled residue of _Sc_Mcm10 (P532) and an AlphaFold3 structure prediction indicates a confident interaction between the extreme C-terminus of _Sc_Mcm10 (a.a. 561-571) and the _Sc_Mcm6 WH (Supplementary Fig. 2K). The extensive interactions between _Sc_Mcm10 and _Sc_Mcm6 modelled in our structure (Fig. 1D) are likely important for CMG activation because truncation of _Sc_Mcm10 beyond a.a. 470 abolished DNA replication with purified proteins (Douglas and Diffley 2016).

### ssDNA binding by Mcm10 is dispensable for CMG activation

Figure 2A shows that the interaction between _Sc_Mcm10 and _Sc_Cdc45 orients the ssDNA binding surface of the OB-fold towards the CMG complex, specifically facing the interface between the _Sc_Mcm2 and _Sc_Mcm5 subunits that has been postulated as an exit gate for ssDNA during CMG activation (Weekes et al. 2026). Note that if ssDNA does exit the MCM ring through this interface, the small interface between the Mcm2 HD and Cdc45 will also need to transiently open for complete ssDNA ejection. A simplistic view of the _Sc_Mcm10:Cdc45 interaction is that it functions to position the OBZnF near the exit gate for ssDNA so that DNA binding by Mcm10 facilitates DNA expulsion from the MCM ring. However, although mutants in the OB-fold have been reported to display phenotypes in yeast (Warren et al. 2008), substitution of three amino acids in the OB-fold abrogated DNA binding but did not affect DNA replication with purified proteins, suggesting that DNA binding is not required for CMG activation (Henrikus, et al. 2024). We attempted to repeat these experiments but consistently recovered low yields of the mutant _Sc_Mcm10 protein so that DNA binding could not be assessed. Consistent with this behaviour, mutation of an equivalent residue in the *X. laevis* MCM10 OB-fold was reported to cause protein insolubility (Warren, et al. 2008). This led us to generate an alternative mutant targeting 7 residues in the _Sc_Mcm10 OBZnF that are predicted by AlphaFold3 to participate in ssDNA binding (Supplementary Fig. 5A-C). The mutations abrogated binding of an _Sc_Mcm10 OBZnF construct to ssDNA (Supplementary Fig. 5D). In the context of full-length Mcm10, the mutant protein (_Sc_Mcm10^DNA^) displayed similar behaviour to wild type during purification and bound to MCM DH (Supplementary Fig. 5E, F). _Sc_Mcm10^DNA^ was evaluated *in vitro* in CMG assembly and activation assays on mini-circle DNA substrates (Douglas, et al. 2018), and in origin-specific DNA replication assays on ∼10 kb linear DNA templates (Yeeles, et al. 2015; Yeeles et al. 2017; Taylor and Yeeles 2018). The mini-circle CMG activation assay monitors topological changes in DNA during CMG activation when compensatory negative supercoils are introduced by topoisomerase 1 as the DNA is unwound (Douglas, et al. 2018) (Fig. 2B). With _Sc_Mcm10^DNA^ there was a modest reduction in the fraction of input DNA converted to the -3 and -4 topoisomers - which are hallmarks of CMG activation (Supplementary Fig. 5G, H) - relative to the wild type protein (Fig. 2C). In the DNA replication assay the products produced in the reaction with _Sc_Mcm10^DNA^ were almost indistinguishable from wild type, demonstrating that DNA binding by Mcm10 is not important for replisome progression in this system (Supplementary Fig. 5I). The absence of detectable ssDNA binding with _Sc_Mcm10^DNA^, together with its subtle defect in the mini-circle assay, confirms that ssDNA binding by _Sc_Mcm10 is not essential for CMG activation *in vitro* (Henrikus, et al. 2024), although it may make a minor contribution to the process. Consistent with these results, yeast expressing Mcm10^DNA^ as the sole copy of Mcm10 did not display growth defects, even when the DNA replication checkpoint was disrupted (Supplementary Fig. 5J). Furthermore, the previously evaluated Mcm10 DNA binding mutant (Henrikus, et al. 2024) also supported normal cell growth, confirming that DNA binding is not an essential property of *S. cerevisiae* Mcm10 (Supplementary Fig. 5K).

**Fig. 2.**
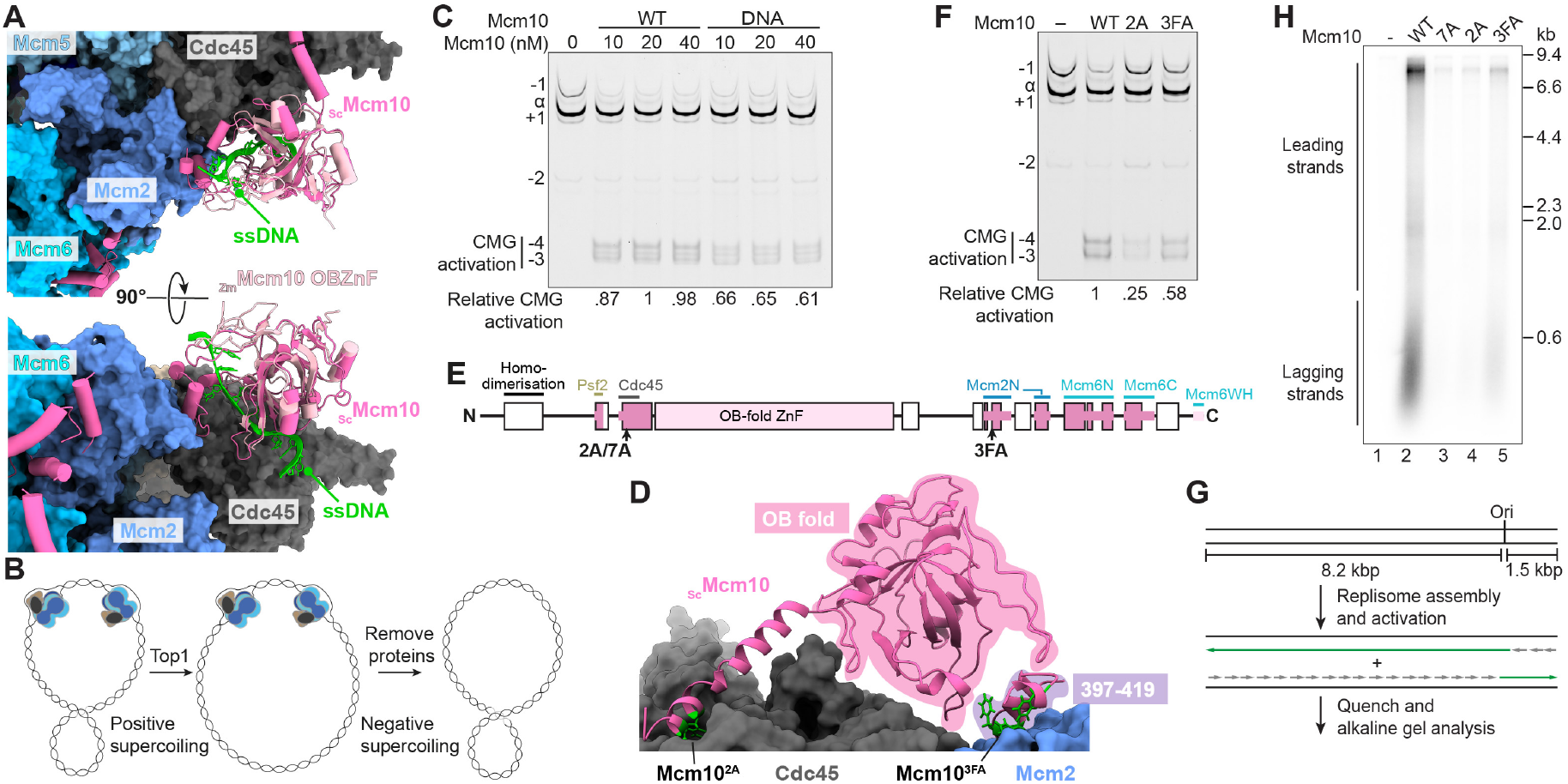
Mcm10 anchoring on Cdc45 is critical for CMG activation. (**A**) Superposition of the *Z. mays* Mcm10 OBZnF bound to a 16 nt ssDNA (PDB: 7Y01) (Zhao, et al. 2023) onto the _Sc_Mcm10 OBZnF. (**B)** Schematic of the mini-circle CMG activation assay. (**C**) CMG activation assay as illustrated in (**B**) on a 657 bp FAM-labelled mini-circle DNA. The relative topological state of each band, as determined in **Supplementary Fig. 5H**, is labelled. The -3 and -4 topoisomers are indicative of CMG activation (see **Supplementary Fig. 5I**). Relative CMG activation is the proportion of mini-circle converted to -3 and -4 topoisomers relative to wild type _Sc_Mcm10 (20 nM). (**D**) Focused view of _Sc_Mcm10 interactions with Mcm2 and Cdc45. Mcm10 residues that mediate interactions and were mutagenized are illustrated as green spheres. **e**, Primary structure diagram of _Sc_Mcm10 (coloured as in **Fig. 1D**) showing the position of mutations. (**F**) Mini-circle CMG activation assay as in (**C**) with the indicated Mcm10 mutants. (**G**) Schematic of the origin-specific DNA replication reaction. (**H**) DNA replication assay performed as illustrated in (**G**) for 12 min with the indicated _Sc_Mcm10 mutants.

### Mcm10:Cdc45 interaction is critical for CMG activation

Because ssDNA binding by Mcm10 is not required for CMG activation (Henrikus, et al. 2024) (Fig. 2C), disrupting the _Sc_Mcm10:Cdc45 interaction should have minimal impact if its main function is to position the OBZnF to bind DNA. This might explain why the Mcm10 α-helix that mediates the interaction with Cdc45 is not broadly conserved and is found only in the Ascomycota phylus of fungi (Supplementary Fig. 6A). To target the interaction, we mutagenized two phenylalanine residues in _Sc_Mcm10 to alanine (_Sc_Mcm10^2A^) (Fig. 2D, E and Supplementary Fig. 5E). Figures 2F-H show that CMG activation and DNA replication were dramatically reduced by the mutations. An additional mutant targeting 7 _Sc_Mcm10 residues at the _Sc_Cdc45 binding interface (_Sc_Mcm10^7A^) (Supplementary Fig. 5E) displayed a comparable defect to _Sc_Mcm10^2A^ in a DNA replication assay (Fig. 2H). By contrast, an _Sc_Mcm10 mutant targeting the interface with the _Sc_Mcm2 HD (_Sc_Mcm10^3FA^) supported more DNA replication and CMG activation (Fig. 2F, H). This attachment is unlikely to significantly constrain the _Sc_Mcm10 OBZnF because it is connected to it via an ∼40-amino-acid linker. However, it does position a C-terminal section of _Sc_Mcm10 against the OB-fold, which might serve to limit its conformational freedom (Fig. 2D and Supplementary Fig. 6B). These data show that the interaction between _Sc_Cdc45 and _Sc_Mcm10 anchors the OBZnF to CMG for efficient helicase activation.

### The Mcm10 C-terminus has a key role in CMG activation, independent of OB fold tethering

A prior study concluded that the C-terminus of _Sc_Mcm10, that includes the Mcm2 and Mcm6 binding sites, functions to tether the OB-fold to CMG to promote helicase activation (Henrikus, et al. 2024). To evaluate this model in the light of our structures, we split _Sc_Mcm10 into two fragments immediately N-terminal of the first modelled binding site on _Sc_Mcm2 (Fig. 3A and Supplementary Fig. 6C). Figure 3B shows that _Sc_Mcm10^C-term^ (a.a. 388-571) did not support CMG activation, whereas CMG activation products were visible with _Sc_Mcm10^N-term^ (1-387), albeit at greatly reduced levels. This is consistent with a previous report that the C-terminal half of _Sc_Mcm10 is not absolutely required for *in vitro* DNA replication (Henrikus, et al. 2024). Strikingly, when both fragments were included, there was a large increase in CMG activation (Fig. 3B). These data demonstrate that the N- and C-terminal halves of _Sc_Mcm10 are both critical for efficient CMG activation and that the primary function of the _Sc_Mcm10 C-terminus is not to tether the OBZnF to CMG via its interactions with Mcm2 and Mcm6. Rather, because the _Sc_Mcm10 C-terminus functions independently of its connection to the OBZnF, we conclude that _Sc_Mcm10 interactions with _Sc_Mcm2, and or _Sc_Mcm6, function to allosterically modulate _Sc_Mcm2-7 during _Sc_CMG activation.

**Fig. 3.**
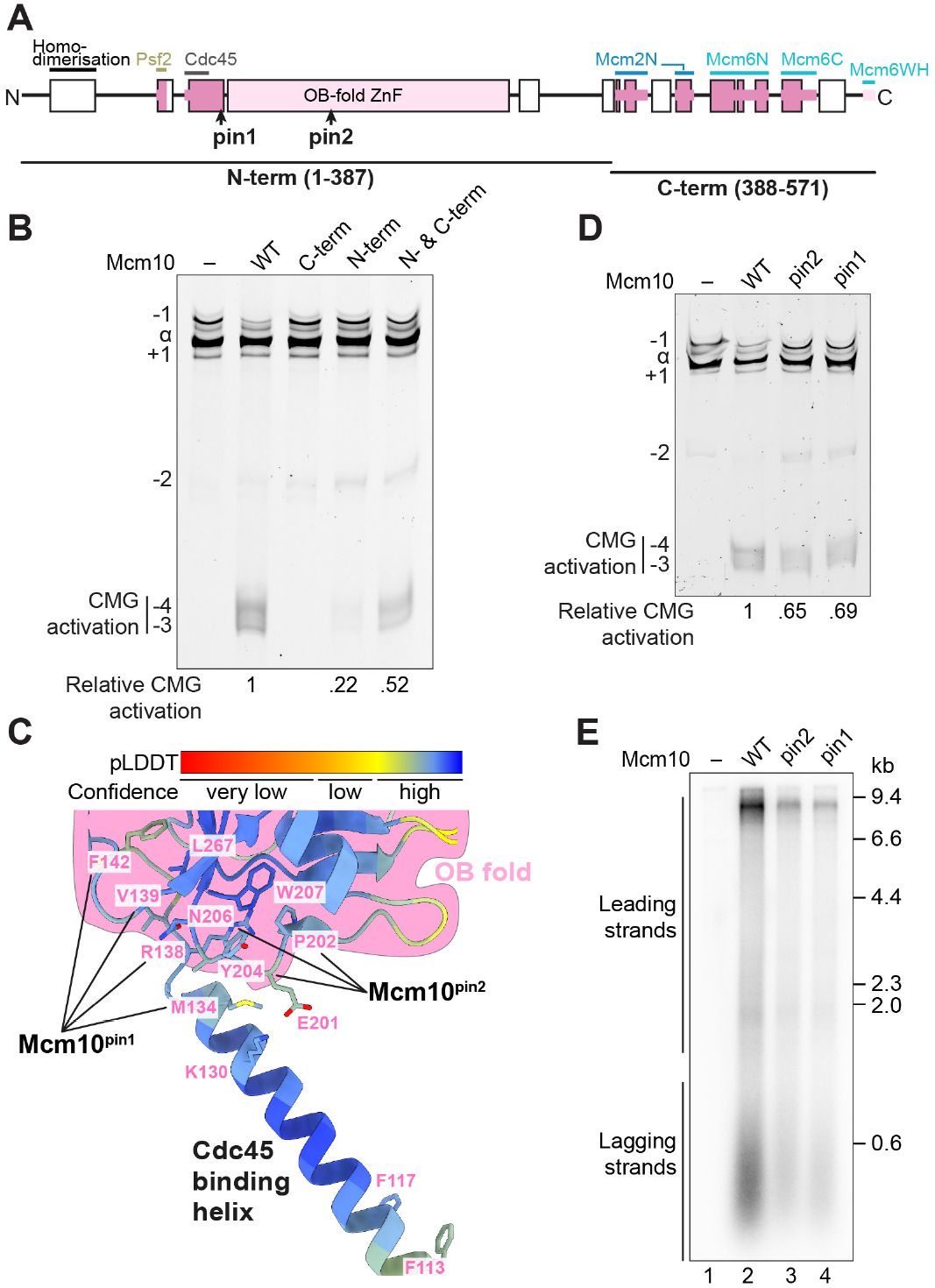
OB fold positioning is important for CMG activation. (**A**) Primary structure diagram of _Sc_Mcm10 (coloured as in **Fig. 1D**) showing the position of mutations and truncations. (**B**) Mini-circle CMG activation assay with _Sc_Mcm10 truncations shown in (**A**). (**C**) Closeup view of the interaction network at the junction between the OB fold and Cdc45 binding helix of _Sc_Mcm10 coloured by pLDDT (AlphaFold3 structure prediction). Amino acids mutated to disrupt this network are labelled. (**D, E**) Mini-circle CMG activation (**D**) and DNA replication (**E**) assays with _Sc_Mcm10 mutants designed to disrupt the connection between the OB fold and Cdc45 binding helix.

### A structural role for the OB-fold in CMG activation

The _Sc_Mcm10 α-helix that binds Cdc45 is predicted to connect to the OB-fold through a short, structured linker that pins the two elements together, restricting flexibility at their junction (Fig. 3C). This limited flexibility is illustrated by predicted aligned error (PAE) values of less than 5Å between the two elements (Supplementary Fig. 6D) and is consistent with the relatively stable positioning of the OB-fold in our structures (Supplementary Fig. 4I, J). Given the importance of the _Sc_Mcm10:Cdc45 interaction (Fig. 2F and H) and the dispensability of ssDNA binding (Henrikus, et al. 2024) (Fig. 2C) for CMG activation, we hypothesised that the OBZnF functions as a structural element, with its precise positioning being important for this role. To test this, we generated two _Sc_Mcm10 mutants targeting the network of interactions at the junction between the Cdc45 binding helix and OB-Fold, _Sc_Mcm10^pin1^ and _Sc_Mcm10^pin2^ (Fig. 3A, C and Supplementary Fig. 6E). Although both mutants retained the ability to bind MCM double hexamers, Cdc45 and ssDNA (Supplementary Fig. 6F-H), they were defective in CMG activation and DNA replication assays (Fig. 3D, E). The interaction of Mcm10 with Cdc45 is therefore not simply a recruitment mechanism; rather, it serves to position the OBZnF in a specific conformation necessary for efficient CMG activation.

### Targeting dimeric CMG

We next sought to address why anchoring the OBZnF on _Sc_Cdc45 is critical for CMG activation. The ZnF is a universal feature of MCM10 proteins (Supplementary Fig. 6A) and its disruption causes lethality in *S. cerevisiae* (Cook, et al. 2003). AlphaFold3 predictions also indicate that MCM10 from diverse eukaryotes contain a short helix of invariant length immediately downstream of the ZnF (ZnF-helix) with no assigned function (Supplementary Fig. 6A). This helix projects away from _Sc_CMG and therefore its putative function was not immediately apparent (Supplementary Fig. 7A). During CMG activation Mcm10 presumably targets dimeric CMG complexes associated via the N-terminal ZnF domains of the Mcm subunits (Lewis, et al. 2022). We therefore docked the model of _Sc_CMG:Mcm10 onto both copies of _Sc_CMG in a CMG dimer structure that was obtained by assembling yeast CMG *in vitro* in the absence of Mcm10 (Lewis, et al. 2022) (Supplementary Fig. 7B). In this arrangement, the ZnF-helix of each Mcm10 protein is oriented toward the Mcm4 subunit of the opposing CMG complex but is not close enough to mediate direct protein–protein interactions. In metazoa, the dimeric firing factor DONSON is essential for CMG assembly (Xia, et al. 2023; Lim et al. 2023; Hashimoto et al. 2023; Kingsley et al. 2023), and cryo-EM analysis of DONSON-bound CMG complexes isolated from *X. laevis* egg extracts revealed a CMG dimer proposed to represent a helicase assembly intermediate (Cvetkovic et al. 2023). Strikingly, docking two copies of _Sc_CMG:Mcm10 onto this structure positioned the ZnF-helices of both _Sc_Mcm10 proteins adjacent to a pocket on the surface of the _Sc_Mcm4 subunit of the opposing MCM hexamer (Fig. 4A). We previously identified this pocket as a conserved binding site for the “anchor” domain of the replisome proteins _Sc_Tof1 (Baretić, et al. 2020) and _Hs_TIMELESS (Morgan L. Jones et al. 2021) and therefore hypothesised it could also be a docking site for the _Sc_Mcm10 ZnF-helix. This idea is strongly supported by AlphaFold3 structure predictions showing a confident interaction between _Sc_Mcm10 and _Sc_Mcm4 that positions the ZnF-helix in the _Sc_Mcm4 pocket (Fig. 4B, C and Supplementary Fig. 7C). Moreover, the predicted interaction shares features of the _Sc_Tof1 anchor interaction including an invariant arginine residue positioned between the OB-fold and HD of _Sc_Mcm4 (Fig. 4D and Supplementary Fig. 7D). The model of _Sc_Mcm10 bound to _Sc_CMG dimers also suggests how Mcm10 homo-dimerisation may contribute to CMG activation. Figure S7E shows that in the context of the CMG dimer, _Sc_Mcm10 binding to Psf2 (GINS) (Fig. 1C, Supplementary Fig. 4M) positions the two _Sc_Mcm10 N-termini so they can interact via an anti-parallel coiled coil that is predicted with high confidence by AlphaFold3 (Supplementary Fig. 7F). Consistently, removal of the _Sc_Mcm10 N-terminus from the N-term fragment impaired CMG activation (Supplementary Fig. 7G). This indicates that Mcm10 homo-oligomerisation contributes to CMG activation, potentially to ensure that two copies of Mcm10 associate with the CMG dimer for synchronous activation of both helicases.

**Fig. 4.**
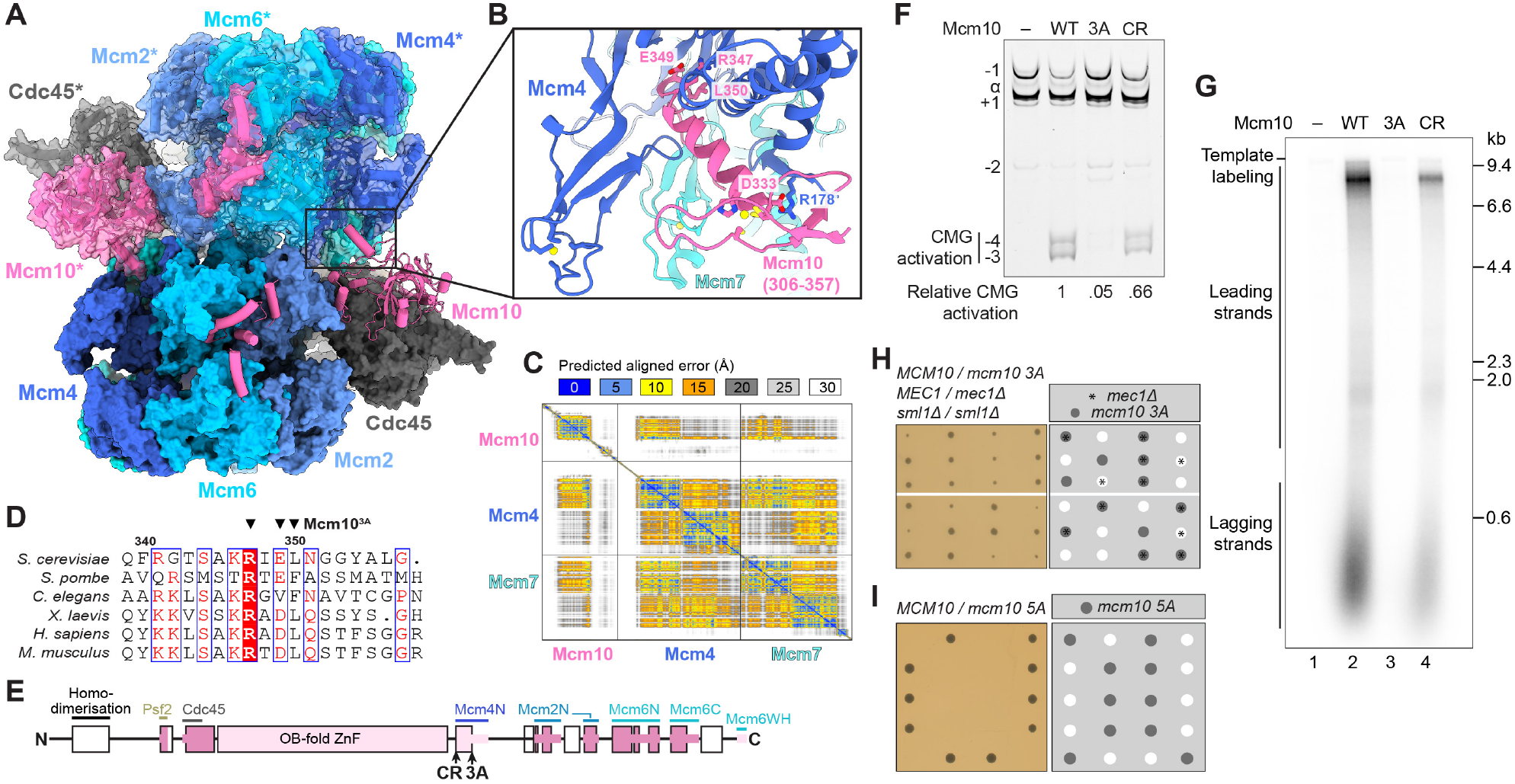
Mcm10 targets dimeric CMG complexes. (**A**) Two copies of the _Sc_CMG-Mcm10 model aligned on the DONSON-mediated *X. laevis* CMG dimer structure (PDB: 8Q6O) (Cvetkovic, et al. 2023). Subunits from the second CMG complex (top) are shown in cartoon representation with a transparent surface and are denoted with a *. (**B**) Closeup view of an AlphaFold3 structure prediction for _Sc_Mcm10 with _Sc_Mcm4 and _Sc_Mcm7. Residue side chains involved in protein:protein interactions that were targeted for mutagenesis are shown as sticks. (**C**) Predicted aligned error plot for the AlphaFold3 structure prediction shown in (**B**). (**D**) Multiple sequence alignment showing the region of Mcm10 predicted to interact with Mcm4. Residues mutated to alanine in _Sc_Mcm10^3A^ are denoted by (▼). (**E**) Primary structure diagram of _Sc_Mcm10 (coloured as in **Fig. 1D**) showing the position of mutations. (**F, G**) CMG activation (**F**) and DNA replication (**G**) assays with _Sc_Mcm10 mutants targeting the predicted interface between Mcm10 and Mcm4. (**H**) Diploid budding yeast cells of the indicated genotype were sporulated and the resulting tetrads dissected and grown on YPD for 2 days at 30°C. Mcm10 5A combines the mutations in Mcm10^3A^ and Mcm10^2A^ targeting the Mcm4 and Cdc45 interactions, respectively.

To examine if the predicted interaction between the _Sc_Mcm10 ZnF-helix and _Sc_Mcm4 is involved in CMG activation, we substituted to alanine three conserved residues in _Sc_Mcm10 (_Sc_Mcm10^3A^) including the arginine residue (R347) that mimics the _Sc_Tof1 anchor interaction (Baretić, et al. 2020) (Fig. 4D, E and Supplementary Fig. 7C, D, H, I). Topological changes associated with CMG activation were severely reduced with _Sc_Mcm10^3A^ and DNA replication was barely detectable (Fig. 4F, G). An _Sc_Mcm10 charge-reversal mutant (_Sc_Mcm10^CR^) targeting two residues at the N-terminus of the ZnF-helix also displayed defects (Fig. 4B, F, G). Despite these defects, yeast cells harbouring the *mcm10-3A* allele grew like wild type (Supplementary Fig. 7J). However, a synthetic growth defect was observed with deletion of the checkpoint kinase Mec1 (Fig. 4H), indicating that *mcm10-3A* cells display a subtle DNA replication defect (Kilkenny et al. 2012). Moreover, combining the Mcm10 3A and 2A (Cdc45 binding site) (Supplementary Fig. 7K) mutations (mcm10-5A) was lethal (Fig. 4I). To conclude, binding of _Sc_Mcm10 to _Sc_Mcm4 via the ZnF-helix is critical for CMG activation, and this mode of engagement is facilitated by _Sc_Mcm10:Cdc45 interactions on the opposing CMG in the context of dimeric helicase complexes. These interactions orient the ssDNA binding surface of the Mcm10 OB-fold towards the Mcm2:Mcm5 interface, strongly suggesting it is poised to sequester ssDNA as it exits the MCM complex through this interface during CMG activation.

### Human MCM10 binding to dimeric CMG

The MCM10 OBZnF and ZnF-helix are universally conserved, however, the helix in _Sc_Mcm10 that mediates the critical interaction with _Sc_Cdc45 is found only in Ascomycota (Supplementary Fig. 6A). These differences raise questions about the extent to which the mechanism of MCM10-mediated CMG activation is conserved. In metazoans, in addition to MCM10, RECQL4 is also involved in CMG activation (Terui et al. 2024; Terui et al. 2025) and interaction between the RECQL4 N-terminus (a.a. 76-145) and _Hs_MCM10 was found to be important for efficient origin firing (Kliszczak et al. 2015). Regulated human CMG assembly has not yet been reconstituted with purified proteins. Therefore, to investigate how _Hs_MCM10 targets _Hs_CMG and how this might be facilitated by _Hs_RECQL4, we developed a strategy to assemble human CMG dimers off DNA for cryo-EM analysis. This approach exploits the capacity of DONSON to dimerise CMG in a configuration that likely represents a CMG assembly intermediate (Cvetkovic, et al. 2023). _Hs_DONSON co-sedimented with _Hs_CMG in a glycerol gradient at higher glycerol concentrations (later fractions) than the peak _Hs_CMG fractions and negative stain EM confirmed that this was due to _Hs_CMG dimerization (Supplementary Fig. 8A-C).

An N-terminal fragment of _Hs_RECQL4 lacking the helicase domain complements RECQL4 deletion in DT40 (Abe et al. 2011), suggesting it contains the regions necessary for CMG activation. Therefore, to simplify our analysis we purified _Hs_RECQL4 (a.a. 1-440) (_Hs_RECQL4^Nterm^) (Supplementary Fig. 8A). Figure S8D shows that _Hs_RECQL4^Nterm^ bound to the _Hs_MCM10-OBZnF, consistent with previous reports (Kliszczak, et al. 2015). Although the functional domains of _Hs_MCM10 involved in CMG activation have not been fully established, we found that the full-length protein was prone to aggregation, an issue we resolved by removing the first 229 a.a. (_Hs_MCM10^∆Nterm^). For complex formation, together with _Hs_CMG, _Hs_DONSON, _Hs_RECQL4^Nterm^ and _Hs_MCM10^∆Nterm^ Pol ε was included because it interacts with _Hs_DONSON (Lim, et al. 2023). Both _Hs_RECQL4^Nterm^ and _Hs_MCM10^∆Nterm^ predominantly sedimented with _Hs_CMG:Pol ε fractions containing _Hs_DONSON, strongly suggesting that they associate preferentially with CMG dimers (Supplementary Fig. 8E). Equivalent fractions from a GraFix gradient were applied directly to graphene oxide grids, buffer exchanged on-grid and vitrified for cryo-EM single particle analysis (Supplementary Fig. 9-12).

The resulting cryo-EM reconstructions reveal dimeric _Hs_CMG bound to _Hs_DONSON, homologous to observations in *X. laevis* (Cvetkovic, et al. 2023) (Fig. 5A). The _Hs_Pol ε non-catalytic module is bound at both copies of CMG in its described position (Jones, et al. 2021), and we recovered and refined to side-chain resolution cryo-EM density for its interaction with the _Hs_DONSON NPF motif (a.a. 76-83) (Keskitalo et al. 2025; Xia, et al. 2023; Lim, et al. 2023) (Fig. 5A and Supplementary Fig. 11C). Moreover, we could model two regions of the _Hs_RECQL4 N-terminus: a short α-helix (a.a. 103-114) bound to a hydrophobic patch on _Hs_CDC45 and a more extensive interaction (a.a. 171-197) with the Pol ε non-catalytic module (Fig. 5B and Supplementary Fig. 11B, C). The structure contains two copies of _Hs_MCM10 with well-defined density for the ZnF-helix, enabling us to visualise the docking site on _Hs_MCM4 (Fig. 5C) that is crucial for CMG activation in *S. cerevisiae* (Fig. 4F, G). The ZnF-helix enters the MCM4 pocket forming side-chain interactions with the N-terminal HD of MCM4, before leading into the invariant R-x-D/E-L motif (Fig. 4D), which docks at the junction of the MCM4 OB, ZnF and HD domains. Clinical data indicates that this interaction is important for MCM10 function in humans because mutation of the invariant arginine to cysteine (R427C) was identified in one allele of a patient with natural killer cell deficiency linked to compound heterozygous mutations in MCM10 (Mace et al. 2020). Following the R-x-D/E-L motif, the _Hs_MCM10 segment traverses the _Hs_MCM4 pocket and exits near the ZnF domain of _Hs_MCM7. Aided by AlphaFold 3 predictions, medium-resolution cryo-EM density at the _Hs_MCM2 HD and _Hs_MCM6 AAA+ domain could be assigned to _Hs_MCM10 residues 515-535 and 557-571, respectively (Supplementary Fig. 12A, B). This illustrates that MCM6 and MCM2 binding by the MCM10 C-terminus is a conserved feature of CMG engagement.

**Fig. 5.**
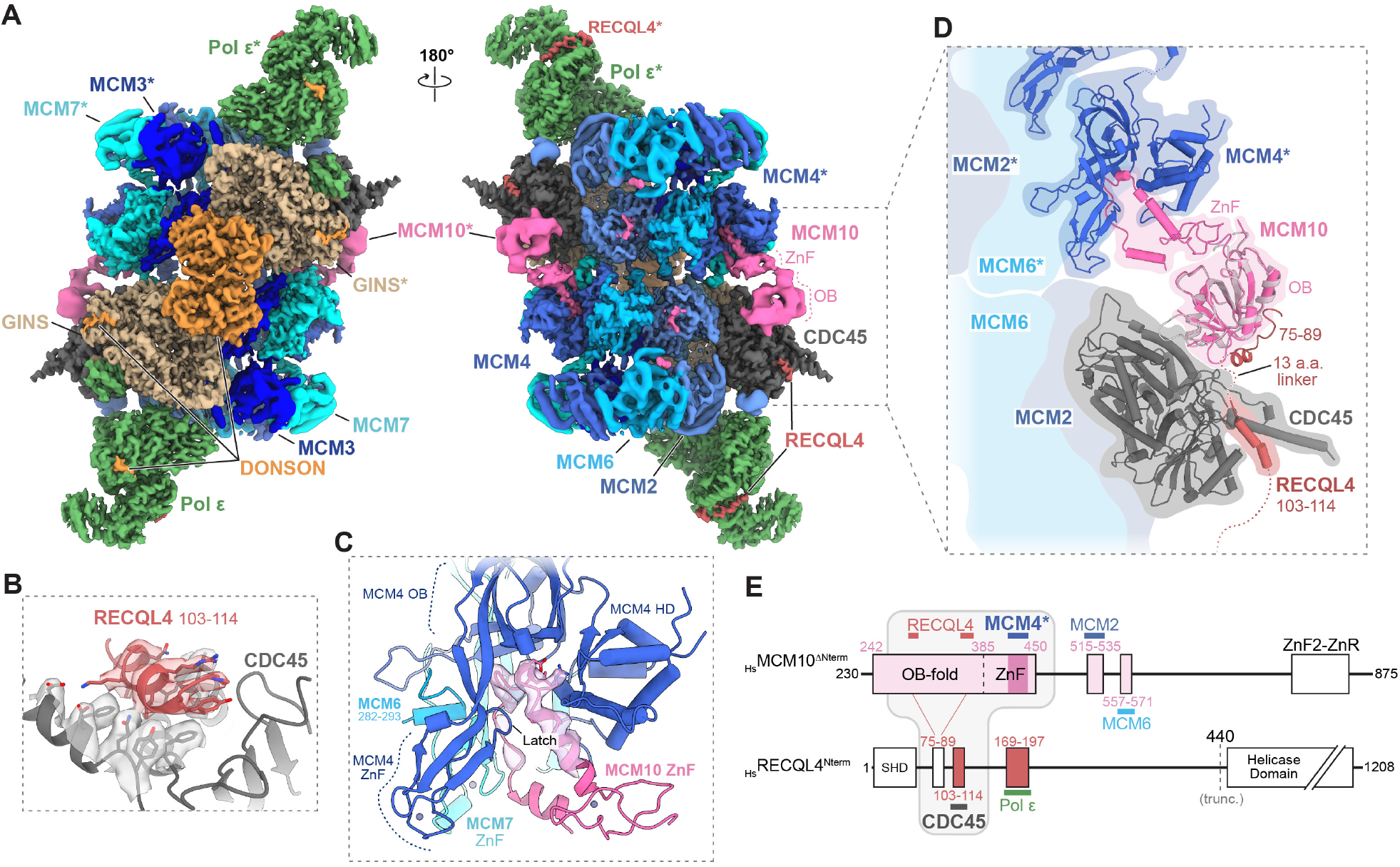
Cryo-EM structure of _Hs_MCM10, _Hs_RECQL4 and _Hs_DONSON bound to dimeric _Hs_CMG:Pol ε. (**A**) Composite of local cryo-EM density maps obtained and constructed as described in Methods, coloured by subunit. Asterisks (*) denote the subunits from the second CMG complex (top). (**B**) Detailed view of the interaction between _Hs_RECQL4 a.a. 103-114 and _Hs_CDC45. Cryo-EM density is shown as semi-transparent surface, with side chain residues involved in the interaction shown as stick models. (**C**) Detailed view of the interaction between _Hs_MCM4* and the ZnF of _Hs_MCM10. Cryo-EM density is shown around the ZnF helix and ensuing MCM4-interacting region of MCM10. Residues equivalent of _Sc_Mcm10^3A^ mutant are shown as sticks. (**D**) View of the interaction network across the two _Hs_CMG copies mediated by _Hs_MCM10 OBZnF and _Hs_RECQL4^Nterm^. Atomic model is shown as cylinders/stubs. An AlphaFold3 structure prediction of the interaction between the _Hs_MCM10 OB-fold and _Hs_RECQL4 a.a. 75-89 is shown as a ribbon model (MCM10 in light grey, RECQL4 in dark red), superimposed on the MCM10 OB-fold. (**E**) Primary structure diagram of _Hs_MCM10^∆Nterm^ and _Hs_RECQL4^Nterm^. The light grey box outlines the network of interactions that bridges _Hs_CDC45 with the _Hs_MCM4 pocket of the opposing CMG (MCM4*). ZnR, zinc finger ribbon; SHD, Sld2 homology domain.

The _Hs_MCM10 OB-fold lies immediately N-terminal of its ZnF domain, but it displays a degree of rotational flexibility that limited the recovery to low-resolution globular density. Nevertheless, the density is the correct shape and volume to accommodate an AlphaFold3 prediction of the _Hs_MCM10 OB-fold, indicating that it displays minimal translational movement relative to CMG (Supplementary Fig. 8H). Notably, superposition of an AlphaFold3 prediction covering the experimentally mapped interaction between the _Hs_MCM10 OB-fold and _Hs_RECQL4 (Kliszczak, et al. 2015) shows that it positions RECQL4 residues 75-89 (shown as ribbon model in Fig. 5D) ∼ 20 Å from the RECQL4 binding site on CDC45 (a.a. 103-114). Because only 13 amino acids separate these sites in RECQL4 (Fig. 5D), the data illustrate that RECQL4 tethers the MCM10 OB-fold to CDC45 (Fig. 5E) and prior studies show that this region of RECQL4 is necessary for efficient origin firing (Abe, et al. 2011; Kliszczak, et al. 2015). Through this tether, and its interactions with the MCM4 pocket of the opposing CMG, the MCM10 OB-fold bridges both copies of CMG. This arrangement resembles our model for _Sc_MCM10 engagement (Fig. 4A), but in *H. sapiens* the RECQL4 tether substitutes for the direct Mcm10:Cdc45 in *S. cerevisiae* (Fig. 5A, D).

### _Hs_MCM10 binding in the _Hs_MCM4 pocket remodels the MCM N-tier of CMG

Within the MCM4 pocket, _Hs_MCM10 is secured behind a β-hairpin formed by a segment of the MCM4 ZnF domain (a.a. 314-324) which acts as a latch (Fig. 5C, “Latch”). This arrangement suggests that a conformational change occurs between the unbound and MCM10-bound states to permit initial engagement of the ZnF-helix while the MCM4 Latch is in an open position, followed by a conformational closure that positions the interacting motif of MCM10 behind the MCM4 Latch. Comparison of _Hs_MCM10-bound CMG with a cryo-EM structure of _Hs_CMG alone (Supplementary Fig. 13), as well as with structures of _Hs_CMG as part of the replisome (Jones, et al. 2021) or bound only to DNA (Batra et al. 2025), indicate that MCM4 ZnF and HD domains and the ZnF domain of MCM7 can shift relative to one another to shape the MCM4 pocket to accommodate different binding partners, including the MCM4 N-terminal region (NTR), TIMELESS and MCM10 (Supplementary Fig. 8F). Strikingly, only in the case of bound MCM10 the MCM4 Latch closes the MCM4 pocket around the interacting motif (Supplementary Fig. 8G). This transition is accompanied by a relative shift between ZnF domains of the MCM ring, with the MCM4 ZnF-Latch moving away from the central pore to close the MCM4 pocket while the MCM7 ZnF shifts toward the center of the MCM ring. Coincident with this rearrangement, we resolve part of the insert within the N-terminal hairpin (NtHp) of the MCM6 OB-Fold (residues 282-292) in this region (Fig. 5C and Supplementary Fig. 12D). The MCM6 NtHp engages with parental DNA in MCM double hexamers (Weissmann, et al. 2024; Yang, et al. 2024) and the elongating CMG complex (Batra, et al. 2025; M. L. Jones et al. 2021), yet it adopts markedly different conformations in these two contexts. The configuration observed in our structure is distinct from either previously described state, indicating that MCM10 binding to the MCM4 pocket has the potential to remodel regions of MCM that are involved in DNA interactions, perhaps to promote DNA strand separation. A second distinction is the conformation of the MCM N-tier more broadly which appears more dilated in the MCM10-bound state than in other CMG structures (Movie S1). Although a direct relationship between the observed local and global changes is not immediately evident, this observation raises the possibility of longer-range allosteric effects of MCM10 binding to CMG with potential implications for gate opening during lagging-strand extrusion.

## Discussion

During CMG activation, dsDNA is melted by ATP-dependent DNA translocation and lagging-strand template ssDNA is expelled from the MCM central pore through the transient opening of an interface between MCM subunits. Our structural and functional analyses of _Sc_Mcm10 (Fig. 1-4) show that precise positioning of the Mcm10 OBZnF, adjacent to the Cdc45-Mcm2 and Mcm2– Mcm5 interfaces, is essential for CMG activation, whereas ssDNA binding is not (Henrikus, et al. 2024). Moreover, the structure of a human CMG dimer assembled by DONSON (Fig. 5) reveals that this positioning is conserved, illustrates how MCM10, aided by RECQL4, targets dimeric CMG by bridging across the dimer, and shows that MCM10 alters the conformation of the MCM N-tier by engaging a pocket on MCM4—an interaction critical for CMG activation in yeast. Based on these discoveries and building on existing literature, we propose a unified model for eukaryotic CMG activation by MCM10 (Fig. 6).

**Fig. 6.**
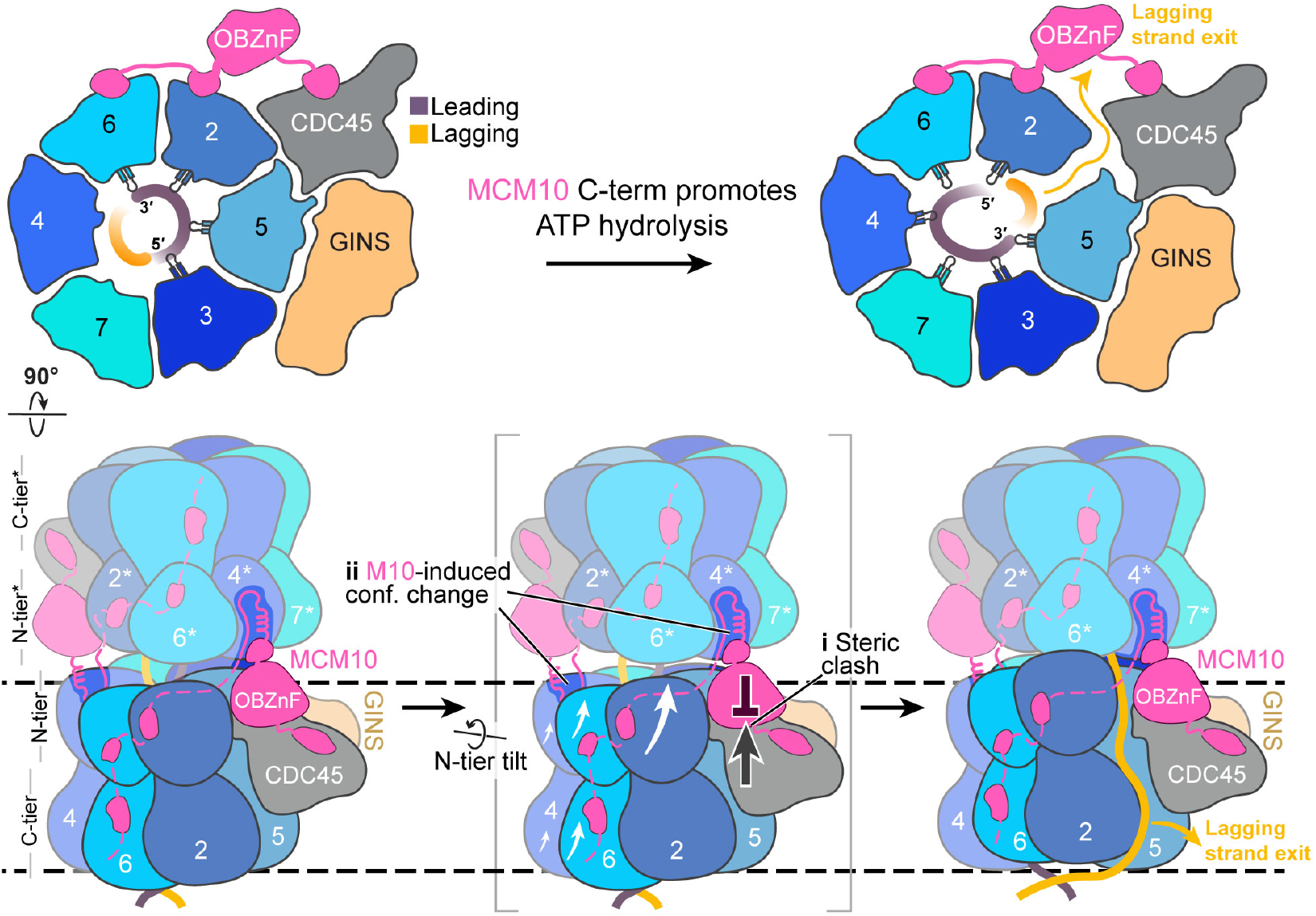
Model for CMG activation by MCM10. After CMG assembly MCM6-2, MCM2-5 and MCM 5-3 engage leading-strand DNA (Lewis, et al. 2022) (top left). The MCM10 C-terminus stimulates ATP hydrolysis at the MCM6-2 and MCM2-5 interfaces to initiate the next steps in the CMG translocation cycle that involve a planar-to-spiral transition of the MCM pore loops (Batra, et al. 2026; Batra, et al. 2025) and concomitant tilt of the MCM N-tier hinged on the MCM3-MCM7 side of the ring (white arrows - middle panel). MCM10 couples these conformational changes to open the MCM2-MCM5 and MCM2-CDC45 interfaces by: (**i**) the OBZnF sterically blocks the movement of CDC45, thereby weakening its interface with MCM2; and / or (**ii**) modulating the conformation of the MCM N-tier by binding within the MCM4 pocket. The lagging strand is now positioned for extrusion through the MCM2-MCM5 interface, after which it may be engaged by the MCM10 OBZnF.

### Model for Mcm10-dependent CMG activation

After CMG assembly DNA translocation is initiated in both CMG complexes (Douglas, et al. 2018), potentially stimulated by C-terminal MCM10 interactions with MCM2 and MCM6 (Fig. 1 and Supplementary Fig. 2G-K). Because neither CMG can advance along DNA or freely rotate due to their head-to-head orientation and inter-hexamer interactions, the DNA within the N-tiers of each MCM ring is sheared(Langston and O’Donnell 2019; Langston and O’Donnell 2017). Simultaneously, conformational changes in the MCM ring that underlie DNA translocation are coupled, via MCM10, to opening of the MCM2–MCM5 and MCM2–CDC45 interfaces, enabling lagging-strand ejection. Although ssDNA binding by the MCM10 OBZnF is not essential for CMG activation (Henrikus, et al. 2024) (Fig. 2C), it is conserved in eukaryotes (Warren, et al. 2009; Henrikus, et al. 2024; Zhao et al. 2023). We propose that ssDNA is sequestered by MCM10 following its ejection before being transferred to the lagging-strand replication machinery, potentially facilitated by Mcm10-Pol α interaction (Warren, et al. 2009; Das-Bradoo, Ricke, and Bielinsky 2006). Once CMG is active, our structures suggest that MCM10 is replaced by the Fork Protection Complex (FPC) (_Sc_Tof1-Csm3-Mrc1 / _Hs_TIM-TIPIN-CLASPIN) and _Sc_Ctf4, whose binding sites overlap almost all MCM10-CMG attachment points (Baretić, et al. 2020; Jones, et al. 2021; Jenkyn-Bedford et al. 2021; Jones et al. 2023) (Supplementary Fig. 14A-C). This indicates that MCM10 is not constitutively associated with elongating replisomes. However, the FPC is displaced from CMG under certain conditions (Somyajit et al. 2017; Westhorpe, Roske, and Yeeles 2024), which might allow for MCM10 reassociation.

During CMG assembly exchange of ADP for ATP drives unwinding of ∼0.6-0.7 turns of the DNA per CMG (Douglas, et al. 2018). _Sc_CMG formed in the absence of _Sc_Mcm10 is ATP bound at the MCM6-2, 2-5 and 5-3 interfaces and these subunits engage the leading-strand template (Lewis, et al. 2022). Although DNA in the MCM C-tier is duplex at this stage, its coordination mirrors a key intermediate in the helical inchworm translocation model (Batra, et al. 2025), with the DNA-bound ATPase pore loops arranged in a planar configuration. In the next step of the helical inchworm model MCM6 and MCM2 dissociate from DNA, likely driven by ATP hydrolysis at the MCM6-2 and 2-5 interfaces, which frees the MCM2 pore loops to advance 12 nucleotides while the neighboring MCM5 subunit maintains its association with DNA (Batra et al. 2026; Batra, et al. 2025). This involves a planar to spiral transition of the MCM pore loops, with MCM6, 4 and 7 also advancing. Consistent with a similar translocation sequence during CMG activation, an Mcm3 arginine finger mutant (targeting the MCM3/7 ATPase) supported CMG assembly but was defective for activation, as measured by RPA recruitment (Kang, Warner, and Bell 2014). The likely requirement for ATP hydrolysis at the MCM6-2 interface is notable because our structures show MCM10 binding to both MCM6 and MCM2 is a conserved feature of the MCM10 C-terminus. We hypothesize that these interactions allosterically modulate the MCM hexamer to initiate and / or sustain ATP hydrolysis and DNA translocation after CMG assembly to facilitate DNA melting.

Relative to the MCM5 ATPase domain, which does not alter its DNA engagement state during the planar-to-spiral transition, the MCM N-tier tilts in a C-tier to N-tier direction (Batra, et al. 2026; Batra, et al. 2025)—i.e., toward the opposing hexamer in the CMG dimer—with MCM7 sitting at the pivot point and CDC45, MCM2 and MCM5 undergoing the largest translational movement. We propose that MCM10 exploits this movement to facilitate MCM2–MCM5 gate opening for ssDNA ejection. Because a minimal _Sc_Mcm10 construct containing only the Cdc45-binding helix, OBZnF and ZnF-helix (Mcm4 binding) is sufficient for CMG activation (Supplementary Fig. 7G), and Drosophila Mcm10 complements Mcm10 deletion in *S. cerevisiae* (Christensen and Tye 2003) despite sharing homology only in the OBZnF and ZnF-helix (Supplementary Fig. 14D, E), this function likely resides within these domains. Our structures suggest two non–mutually exclusive mechanisms. First, because _Hs_MCM10 binding in the MCM4 pocket locally (Supplementary Fig. 8G) – and perhaps globally (Movie S1) – remodels the MCM N-tier, we propose that allosteric changes in the MCM N-tier, driven by MCM10 binding to MCM4, weaken the MCM2-MCM5 interface to trigger its opening. Second, owing to its positioning between CDC45 and MCM4 of the opposing CMG - which is critical for CMG activation in yeast (Fig. 1-4) - the OB fold is a steric block that limits CDC45 movement towards the opposing CMG during the planar-to-spiral transition, destabilising the MCM2–CDC45 interface and consequently the MCM2–MCM5 gate for strand ejection. Because the OBZnF is more rigidly anchored to CDC45 in yeast than human, this mechanism may be more a prominent feature of yeast CMG activation. We speculate this might reflect the presence of DONSON, which is absent in yeast, aiding MCM10 during human CMG activation by restraining movement between the two helicases. This might also explain why MCM10 is not essential in *C. elegans* (Xia, et al. 2023) or for DNA replication in *X. laevis* egg extracts (Chadha, et al. 2016). Nevertheless, tethering of MCM10 to CDC45 is necessary for efficient CMG activation in metazoa and this is achieved through RECQL4. However, this is unlikely to be the only mechanism by which RECQL4 contributes to CMG activation because a minimal construct containing only the MCM10 and CDC45 interaction motifs did not complement RECQL4 loss in DT40 (Abe, et al. 2011). Further work is required to fully understand how RECQL4 promotes CMG activation, including the function of its interaction with Pol ε (Fig. 5A).

In support of our model for CMG activation by MCM10, a two–amino acid substitution in Mcm2, predicted to destabilize the interface between its OB-fold and helical domain (Supplementary Fig. 14F), bypasses the requirement for Mcm10 in *in vitro* yeast DNA replication assays (Looke, Maloney, and Bell 2017). It is plausible these mutations weaken the Mcm2-Cdc45 interface sufficiently so that ATP-dependent conformational changes in MCM drive Mcm2-5 gate opening in the absence of Mcm10. These data and the structures presented herein support the notion that the CMG complex drives its own activation with MCM10 augmenting less favorable mechanistic steps.

## Acknowledgments

We thank J. Shi and K. Turton for operation of the LMB baculovirus facility; F. Fassetta for baculovirus generation and protein expression; M. Hodskinson and Y. Baris for generating protein expression constructs; LMB media and glass wash for cell culture media; S. Chen, G. Sharov, G. Cannone, A. Yeates, B. Ahsan, D. Bellini and H. Safdari for smooth running of the MRC LMB EM facility; S. McLaughlin for assistance with fluorescence polarisation experiments; J. Grimmett, T. Darling, and I. Clayson for maintenance of scientific computing facilities.

## Author contributions

Conceptualization: MLJ, JTPY; Methodology: ETC, JJR, EEF, JTPY; Investigation: ETC, MLJ, JJR, EEF, IXRC, JTPY; Funding acquisition: JTPY; Supervision: JTPY; Writing – original draft: ETC, MLJ, JJR, EEF, IXRC, JTPY; Writing – review & editing: ETC, MLJ, JJR, EEF, IXRC, JTPY

## Data, code, and materials availability

Depositions of cryo-EM maps to EMDB and atomic models to PDB will be made at a later date, prior to publication. The authors declare that all other data supporting the findings of this study are available within the paper and its Supplementary Information files.

## References

Abe, T., A. Yoshimura, Y. Hosono, S. Tada, M. Seki, and T. Enomoto. 2011. “The N-Terminal Region of Recql4 Lacking the Helicase Domain Is Both Essential and Sufficient for the Viability of Vertebrate Cells. Role of the N-Terminal Region of Recql4 in Cells.” Biochim Biophys Acta 1813, no. 3 (Mar): 473–9. 10.1016/j.bbamcr.2011.01.001.

Abramson, J., J. Adler, J. Dunger, R. Evans, T. Green, A. Pritzel, O. Ronneberger, L. Willmore, A. J. Ballard, J. Bambrick, S. W. Bodenstein, D. A. Evans, C. C. Hung, M. O’Neill, D. Reiman, K. Tunyasuvunakool, Z. Wu, A. Zemgulyte, E. Arvaniti, C. Beattie, O. Bertolli, A. Bridgland, A. Cherepanov, M. Congreve, A. I. Cowen-Rivers, A. Cowie, M. Figurnov, F. B. Fuchs, H. Gladman, R. Jain, Y. A. Khan, C. M. R. Low, K. Perlin, A. Potapenko, P. Savy, S. Singh, A. Stecula, A. Thillaisundaram, C. Tong, S. Yakneen, E. D. Zhong, M. Zielinski, A. Zidek, V. Bapst, P. Kohli, M. Jaderberg, D. Hassabis, and J. M. Jumper. 2024. “Accurate Structure Prediction of Biomolecular Interactions with Alphafold 3.” Nature 630, no. 8016 (Jun): 493–500. 10.1038/s41586-024-07487-w.

Afonine, P. V., B. K. Poon, R. J. Read, O. V. Sobolev, T. C. Terwilliger, A. Urzhumtsev, and P. D. Adams. 2018. “Real-Space Refinement in Phenix for Cryo-Em and Crystallography.” Acta Crystallogr D Struct Biol 74, no. Pt 6 (Jun 1): 531–544. 10.1107/S2059798318006551.

Alver, R. C., T. Zhang, A. Josephrajan, B. L. Fultz, C. J. Hendrix, S. Das-Bradoo, and A. K. Bielinsky. 2014. “The N-Terminus of Mcm10 Is Important for Interaction with the 9-1-1 Clamp and in Resistance to DNA Damage.” Nucleic Acids Res 42, no. 13 (Jul): 8389–404. 10.1093/nar/gku479.

Baretić, Domagoj, Michael Jenkyn-Bedford, Valentina Aria, Giuseppe Cannone, Mark Skehel, and Joseph T. P. Yeeles. 2020. “Cryo-Em Structure of the Fork Protection Complex Bound to Cmg at a Replication Fork.” Molecular Cell 78, no. 5 (2020/06/04/): 926–940.e13. 10.1016/j.molcel.2020.04.012.

Baris, Y., M. R. G. Taylor, V. Aria, and J. T. P. Yeeles. 2022. “Fast and Efficient DNA Replication with Purified Human Proteins.” Nature 606, no. 7912 (Jun): 204–210. 10.1038/s41586-022-04759-1.

Batra, S., B. Allwein, C. Kumar, S. Devbhandari, J. G. Bruning, S. Bahng, C. M. Lee, K. J. Marians, R. K. Hite, and D. Remus. 2025. “G-Quadruplex-Stalled Eukaryotic Replisome Structure Reveals Helical Inchworm DNA Translocation.” Science 387, no. 6738 (Mar 7): eadt1978. 10.1126/science.adt1978.

Batra, S., B. Allwein, Y. L. Wang, R. K. Hite, and D. Remus. 2026. “DNA Translocation by the Cmg Helicase: The Helical Inchworm Model.” Biochem Soc Trans 54, no. 2 (Feb 25). 10.1042/BST20250145.

Chadha, G. S., A. Gambus, P. J. Gillespie, and J. J. Blow. 2016. “Xenopus Mcm10 Is a Cdk-Substrate Required for Replication Fork Stability.” Cell Cycle 15, no. 16 (Aug 17): 2183–2195. 10.1080/15384101.2016.1199305.

Chen, V. B., W. B. Arendall, 3rd, J. J. Headd, D. A. Keedy, R. M. Immormino, G. J. Kapral, L. W. Murray, J. S. Richardson, and D. C. Richardson. 2010. “Molprobity: All-Atom Structure Validation for Macromolecular Crystallography.” Acta Crystallogr D Biol Crystallogr 66, no. Pt 1 (Jan): 12–21. 10.1107/S0907444909042073.

Christensen, T. W., and B. K. Tye. 2003. “Drosophila Mcm10 Interacts with Members of the Prereplication Complex and Is Required for Proper Chromosome Condensation.” Mol Biol Cell 14, no. 6 (Jun): 2206–15. 10.1091/mbc.e02-11-0706.

Cook, C. R., G. Kung, F. C. Peterson, B. F. Volkman, and M. Lei. 2003. “A Novel Zinc Finger Is Required for Mcm10 Homocomplex Assembly.” J Biol Chem 278, no. 38 (Sep 19): 36051–8. 10.1074/jbc.M306049200.

Croll, T. I. 2018. “Isolde: A Physically Realistic Environment for Model Building into Low-Resolution Electron-Density Maps.” Acta Crystallogr D Struct Biol 74, no. Pt 6 (Jun 1): 519–530. 10.1107/S2059798318002425.

Cvetkovic, M. A., P. Passaretti, A. Butryn, A. Reynolds-Winczura, G. Kingsley, A. Skagia, C. Fernandez-Cuesta, D. Poovathumkadavil, R. George, A. S. Chauhan, S. S. Jhujh, G. S. Stewart, A. Gambus, and A. Costa. 2023. “The Structural Mechanism of Dimeric Donson in Replicative Helicase Activation.” Mol Cell 83, no. 22 (Nov 16): 4017–4031 e9. 10.1016/j.molcel.2023.09.029.

Das-Bradoo, S., R. M. Ricke, and A. K. Bielinsky. 2006. “Interaction between Pcna and Diubiquitinated Mcm10 Is Essential for Cell Growth in Budding Yeast.” Mol Cell Biol 26, no. 13 (Jul): 4806–17. 10.1128/MCB.02062-05.

Douglas, M. E., F. A. Ali, A. Costa, and J. F. X. Diffley. 2018. “The Mechanism of Eukaryotic Cmg Helicase Activation.” Nature 555, no. 7695 (Mar 8): 265–268. 10.1038/nature25787.

Douglas, M. E., and J. F. Diffley. 2016. “Recruitment of Mcm10 to Sites of Replication Initiation Requires Direct Binding to the Minichromosome Maintenance (Mcm) Complex.” J Biol Chem 291, no. 11 (Mar 11): 5879–88. 10.1074/jbc.M115.707802.

Du, W., A. Josephrajan, S. Adhikary, T. Bowles, A. K. Bielinsky, and B. F. Eichman. 2013. “Mcm10 Self-Association Is Mediated by an N-Terminal Coiled-Coil Domain.” PLoS One 8, no. 7: e70518. 10.1371/journal.pone.0070518.

Dumas, L. B., J. P. Lussky, E. J. McFarland, and J. Shampay. 1982. “New Temperature-Sensitive Mutants of Saccharomyces Cerevisiae Affecting DNA Replication.” Mol Gen Genet 187, no. 1: 42–6. 10.1007/BF00384381.

Eisenberg, S., G. Korza, J. Carson, I. Liachko, and B. K. Tye. 2009. “Novel DNA Binding Properties of the Mcm10 Protein from Saccharomyces Cerevisiae.” J Biol Chem 284, no. 37 (Sep 11): 25412–20. 10.1074/jbc.M109.033175.

Emsley, P., B. Lohkamp, W. G. Scott, and K. Cowtan. 2010. “Features and Development of Coot.” Acta Crystallogr D Biol Crystallogr 66, no. Pt 4 (Apr): 486–501. 10.1107/S0907444910007493.

Evrin, C., P. Clarke, J. Zech, R. Lurz, J. Sun, S. Uhle, H. Li, B. Stillman, and C. Speck. 2009. “A Double-Hexameric Mcm2-7 Complex Is Loaded onto Origin DNA during Licensing of Eukaryotic DNA Replication.” Proc Natl Acad Sci U S A 106, no. 48 (Dec 1): 20240–5. 10.1073/pnas.0911500106.

Fien, K., Y. S. Cho, J. K. Lee, S. Raychaudhuri, I. Tappin, and J. Hurwitz. 2004. “Primer Utilization by DNA Polymerase Alpha-Primase Is Influenced by Its Interaction with Mcm10p.” J Biol Chem 279, no. 16 (Apr 16): 16144–53. 10.1074/jbc.M400142200.

Frigola, J., D. Remus, A. Mehanna, and J. F. Diffley. 2013. “Atpase-Dependent Quality Control of DNA Replication Origin Licensing.” Nature 495, no. 7441 (Mar 21): 339–43. 10.1038/nature11920.

Georgescu, R., Z. N. Yuan, L. Bai, R. D. A. Santos, J. C. Sun, D. Zhang, O. Yurieva, H. L. Li, and M. E. O’Donnell. 2017. “Structure of Eukaryotic Cmg Helicase at a Replication Fork and Implications to Replisome Architecture and Origin Initiation.” Proceedings of the National Academy of Sciences of the United States of America 114, no. 5 (Jan 31): E697–E706. 10.1073/pnas.1620500114.

Hashimoto, Y., K. Sadano, N. Miyata, H. Ito, and H. Tanaka. 2023. “Novel Role of Donson in Cmg Helicase Assembly during Vertebrate DNA Replication Initiation.” EMBO J 42, no. 17 (Sep 4): e114131. 10.15252/embj.2023114131.

Henrikus, S. S., M. H. Gross, O. Willhoft, T. Puhringer, J. S. Lewis, A. W. McClure, J. F. Greiwe, G. Palm, A. Nans, J. F. X. Diffley, and A. Costa. 2024. “Unwinding of a Eukaryotic Origin of Replication Visualized by Cryo-Em.” Nat Struct Mol Biol 31, no. 8 (Aug): 1265–1276. 10.1038/s41594-024-01280-z.

Homesley, L., M. Lei, Y. Kawasaki, S. Sawyer, T. Christensen, and B. K. Tye. 2000. “Mcm10 and the Mcm2-7 Complex Interact to Initiate DNA Synthesis and to Release Replication Factors from Origins.” Genes Dev 14, no. 8 (Apr 15): 913–26. https://www.ncbi.nlm.nih.gov/pubmed/10783164.

Jenkyn-Bedford, M., M. L. Jones, Y. Baris, K. P. M. Labib, G. Cannone, J. T. P. Yeeles, and T. D. Deegan. 2021. “A Conserved Mechanism for Regulating Replisome Disassembly in Eukaryotes.” Nature 600, no. 7890 (Dec): 743–747. 10.1038/s41586-021-04145-3.

Jones, M. L., V. Aria, Y. Baris, and J. T. P. Yeeles. 2023. “How Pol Alpha-Primase Is Targeted to Replisomes to Prime Eukaryotic DNA Replication.” Mol Cell 83, no. 16 (Aug 17): 2911–2924 e16. 10.1016/j.molcel.2023.06.035.

Jones, M. L., Y. Baris, M. R. G. Taylor, and J. T. P. Yeeles. 2021. “Structure of a Human Replisome Shows the Organisation and Interactions of a DNA Replication Machine.” EMBO J 40, no. 23 (Dec 1): e108819. 10.15252/embj.2021108819.

Jones, Morgan L., Yasemin Baris, Martin R. G. Taylor, and Joseph T. P. Yeeles. 2021. “Structure of a Human Replisome Shows the Organisation and Interactions of a DNA Replication Machine.” The EMBO Journal n/a, no. n/a: e108819. 10.15252/embj.2021108819

Kang, S., M. D. Warner, and S. P. Bell. 2014. “Multiple Functions for Mcm2-7 Atpase Motifs during Replication Initiation.” Mol Cell 55, no. 5 (Sep 4): 655–65. 10.1016/j.molcel.2014.06.033.

Kanke, M., Y. Kodama, T. S. Takahashi, T. Nakagawa, and H. Masukata. 2012. “Mcm10 Plays an Essential Role in Origin DNA Unwinding after Loading of the Cmg Components.” EMBO J 31, no. 9 (May 2): 2182–94. 10.1038/emboj.2012.68.

Kastner, B., N. Fischer, M. M. Golas, B. Sander, P. Dube, D. Boehringer, K. Hartmuth, J. Deckert, F. Hauer, E. Wolf, H. Uchtenhagen, H. Urlaub, F. Herzog, J. M. Peters, D. Poerschke, R. Luhrmann, and H. Stark. 2008. “Grafix: Sample Preparation for Single-Particle Electron Cryomicroscopy.” Nat Methods 5, no. 1 (Jan): 53–5. 10.1038/nmeth1139.

Keskitalo, S., B. Zambo, D. Malaymar Pinar, A. Tuhkala, K. Salokas, T. Turunen, N. Deutsch, N. Davey, Z. Dosztanyi, M. Varjosalo, and G. Gogl. 2025. “The Non-Catalytic DNA Polymerase Epsilon Subunit Is an Npf Motif Recognition Protein.” Nat Commun 17, no. 1 (Dec 13): 586. 10.1038/s41467-025-67284-5.

Kilkenny, M. L., G. De Piccoli, R. L. Perera, K. Labib, and L. Pellegrini. 2012. “A Conserved Motif in the C-Terminal Tail of DNA Polymerase Alpha Tethers Primase to the Eukaryotic Replisome.” J Biol Chem 287, no. 28 (Jul 6): 23740–7. 10.1074/jbc.M112.368951.

Kingsley, G., A. Skagia, P. Passaretti, C. Fernandez-Cuesta, A. Reynolds-Winczura, K. Koscielniak, and A. Gambus. 2023. “Donson Facilitates Cdc45 and Gins Chromatin Association and Is Essential for DNA Replication Initiation.” Nucleic Acids Res 51, no. 18 (Oct 13): 9748–9763. 10.1093/nar/gkad694.

Kliszczak, M., H. Sedlackova, G. P. Pitchai, W. W. Streicher, L. Krejci, and I. D. Hickson. 2015. “Interaction of Recq4 and Mcm10 Is Important for Efficient DNA Replication Origin Firing in Human Cells.” Oncotarget 6, no. 38 (Dec 1): 40464–79. 10.18632/oncotarget.6342.

Langston, L. D., and M. E. O’Donnell. 2019. “An Explanation for Origin Unwinding in Eukaryotes.” Elife 8 (Jul 8). 10.7554/eLife.46515.

Langston, L., and M. O’Donnell. 2017. “Action of Cmg with Strand-Specific DNA Blocks Supports an Internal Unwinding Mode for the Eukaryotic Replicative Helicase.” Elife 6 (Mar 27): e23449. 10.7554/eLife.23449.

Lewis, J. S., M. H. Gross, J. Sousa, S. S. Henrikus, J. F. Greiwe, A. Nans, J. F. X. Diffley, and A. Costa. 2022. “Mechanism of Replication Origin Melting Nucleated by Cmg Helicase Assembly.” Nature 606, no. 7916 (Jun): 1007–1014. 10.1038/s41586-022-04829-4.

Lim, Y., L. Tamayo-Orrego, E. Schmid, Z. Tarnauskaite, O. V. Kochenova, R. Gruar, S. Muramatsu, L. Lynch, A. V. Schlie, P. L. Carroll, G. Chistol, M. A. M. Reijns, M. T. Kanemaki, A. P. Jackson, and J. C. Walter. 2023. “In Silico Protein Interaction Screening Uncovers Donson’s Role in Replication Initiation.” Science 381, no. 6664 (Sep 22): eadi3448. 10.1126/science.adi3448.

Looke, M., M. F. Maloney, and S. P. Bell. 2017. “Mcm10 Regulates DNA Replication Elongation by Stimulating the Cmg Replicative Helicase.” Genes Dev 31, no. 3 (Feb 1): 291–305. 10.1101/gad.291336.116.

Mace, E. M., S. Paust, M. I. Conte, R. M. Baxley, M. M. Schmit, S. L. Patil, N. C. Guilz, M. Mukherjee, A. E. Pezzi, J. Chmielowiec, S. Tatineni, I. K. Chinn, Z. C. Akdemir, S. N. Jhangiani, D. M. Muzny, A. Stray-Pedersen, R. E. Bradley, M. Moody, P. P. Connor, A. G. Heaps, C. Steward, P. P. Banerjee, R. A. Gibbs, M. Borowiak, J. R. Lupski, S. Jolles, A. K. Bielinsky, and J. S. Orange. 2020. “Human Nk Cell Deficiency as a Result of Biallelic Mutations in Mcm10.” J Clin Invest 130, no. 10 (Oct 1): 5272–5286. 10.1172/JCI134966.

Maine, G. T., P. Sinha, and B. K. Tye. 1984. “Mutants of S. Cerevisiae Defective in the Maintenance of Minichromosomes.” Genetics 106, no. 3 (Mar): 365–85. https://www.ncbi.nlm.nih.gov/pubmed/6323245.

Mayle, R., L. Langston, K. R. Molloy, D. Zhang, B. T. Chait, and M. E. O’Donnell. 2019. “Mcm10 Has Potent Strand-Annealing Activity and Limits Translocase-Mediated Fork Regression.” Proc Natl Acad Sci U S A 116, no. 3 (Jan 15): 798–803. 10.1073/pnas.1819107116.

Nasmyth, K., and P. Nurse. 1981. “Cell Division Cycle Mutants Altered in DNA Replication and Mitosis in the Fission Yeast Schizosaccharomyces Pombe.” Mol Gen Genet 182, no. 1: 119–24. 10.1007/BF00422777.

Pettersen, E. F., T. D. Goddard, C. C. Huang, G. S. Couch, D. M. Greenblatt, E. C. Meng, and T. E. Ferrin. 2004. “Ucsf Chimera--a Visualization System for Exploratory Research and Analysis.” J Comput Chem 25, no. 13 (Oct): 1605–12. 10.1002/jcc.20084.

Punjani, A., and D. J. Fleet. 2021. “3d Variability Analysis: Resolving Continuous Flexibility and Discrete Heterogeneity from Single Particle Cryo-Em.” J Struct Biol 213, no. 2 (Jun): 107702. 10.1016/j.jsb.2021.107702.

Punjani, A., and D. J. Fleet.. 2023. “3dflex: Determining Structure and Motion of Flexible Proteins from Cryo-Em.” Nat Methods 20, no. 6 (Jun): 860–870. 10.1038/s41592-023-01853-8.

Punjani, A., H. Zhang, and D. J. Fleet. 2020. “Non-Uniform Refinement: Adaptive Regularization Improves Single-Particle Cryo-Em Reconstruction.” Nat Methods 17, no. 12 (Dec): 1214–1221. 10.1038/s41592-020-00990-8.

Punjani, Ali, John L. Rubinstein, David J. Fleet, and Marcus A. Brubaker. 2017. “Cryosparc: Algorithms for Rapid Unsupervised Cryo-Em Structure Determination.” Nature Methods 14, no. 3 (2017/03/01): 290–296. 10.1038/nmeth.4169.

Punjani, Ali, Haowei Zhang, and David J. Fleet. 2020. “Non-Uniform Refinement: Adaptive Regularization Improves Single-Particle Cryo-Em Reconstruction.” Nature Methods 17, no. 12 (2020/12/01): 1214–1221. 10.1038/s41592-020-00990-8.

Quan, Y., Y. Xia, L. Liu, J. Cui, Z. Li, Q. Cao, X. S. Chen, J. L. Campbell, and H. Lou. 2015. “Cell-Cycle-Regulated Interaction between Mcm10 and Double Hexameric Mcm2-7 Is Required for Helicase Splitting and Activation during S Phase.” Cell Rep 13, no. 11 (Dec 22): 2576–2586. 10.1016/j.celrep.2015.11.018.

Remus, D., F. Beuron, G. Tolun, J. D. Griffith, E. P. Morris, and J. F. Diffley. 2009. “Concerted Loading of Mcm2-7 Double Hexamers around DNA during DNA Replication Origin Licensing.” Cell 139, no. 4 (Nov 13): 719–30. 10.1016/j.cell.2009.10.015.

Robertson, P. D., E. M. Warren, H. Zhang, D. B. Friedman, J. W. Lary, J. L. Cole, A. V. Tutter, J. C. Walter, E. Fanning, and B. F. Eichman. 2008. “Domain Architecture and Biochemical Characterization of Vertebrate Mcm10.” J Biol Chem 283, no. 6 (Feb 8): 3338–3348. 10.1074/jbc.M706267200.

Roske, J. J., and J. T. P. Yeeles. 2024. “Structural Basis for Processive Daughter-Strand Synthesis and Proofreading by the Human Leading-Strand DNA Polymerase Pol Epsilon.” Nat Struct Mol Biol 31, no. 12 (Dec): 1921–1931. 10.1038/s41594-024-01370-y.

Rzechorzek, Neil J., Steven W. Hardwick, Vincentius A. Jatikusumo, Dimitri Y Chirgadze, and Luca Pellegrini. 2020. “Cryoem Structures of Human Cmg–Atpγs–DNA and Cmg–and-1 Complexes.” Nucleic Acids Research 48, no. 12: 6980–6995. Accessed 4/12/2021. 10.1093/nar/gkaa429.

Samel, S. A., A. Fernandez-Cid, J. Sun, A. Riera, S. Tognetti, M. C. Herrera, H. Li, and C. Speck. 2014. “A Unique DNA Entry Gate Serves for Regulated Loading of the Eukaryotic Replicative Helicase Mcm2-7 onto DNA.” Genes Dev 28, no. 15 (Aug 1): 1653–66. 10.1101/gad.242404.114.

Scheres, S. H. 2012. “Relion: Implementation of a Bayesian Approach to Cryo-Em Structure Determination.” J Struct Biol 180, no. 3 (Dec): 519–30. 10.1016/j.jsb.2012.09.006.

Schmid, E. W., and J. C. Walter. 2025. “Predictomes, a Classifier-Curated Database of Alphafold-Modeled Protein-Protein Interactions.” Mol Cell 85, no. 6 (Mar 20): 1216–1232 e5. 10.1016/j.molcel.2025.01.034.

Somyajit, K., R. Gupta, H. Sedlackova, K. J. Neelsen, F. Ochs, M. B. Rask, C. Choudhary, and J. Lukas. 2017. “Redox-Sensitive Alteration of Replisome Architecture Safeguards Genome Integrity.” Science 358, no. 6364 (Nov 10): 797–802. 10.1126/science.aao3172.

Taylor, M. R. G., and J. T. P. Yeeles. 2018. “The Initial Response of a Eukaryotic Replisome to DNA Damage.” Mol Cell 70, no. 6 (Jun 21): 1067–1080 e12. 10.1016/j.molcel.2018.04.022.

Terui, R., S. E. Berger, L. A. Sambel, D. Song, and G. Chistol. 2024. “Single-Molecule Imaging Reveals the Mechanism of Bidirectional Replication Initiation in Metazoa.” Cell 187, no. 15 (Jul 25): 3992–4009 e25. 10.1016/j.cell.2024.05.024.

Terui, R., L. Sambel, D. Song, and G. Chistol. 2025. “Mcm10 and Recql4 Synergize to Activate the Eukaryotic Replicative DNA Helicase.” bioRxiv (Aug 6). 10.1101/2025.08.05.668837.

van Deursen, F., S. Sengupta, G. De Piccoli, A. Sanchez-Diaz, and K. Labib. 2012. “Mcm10 Associates with the Loaded DNA Helicase at Replication Origins and Defines a Novel Step in Its Activation.” EMBO J 31, no. 9 (May 2): 2195–206. 10.1038/emboj.2012.69.

Warren, E. M., H. Huang, E. Fanning, W. J. Chazin, and B. F. Eichman. 2009. “Physical Interactions between Mcm10, DNA, and DNA Polymerase Alpha.” J Biol Chem 284, no. 36 (Sep 4): 24662–72. 10.1074/jbc.M109.020438.

Warren, E. M., S. Vaithiyalingam, J. Haworth, B. Greer, A. K. Bielinsky, W. J. Chazin, and B. F. Eichman. 2008. “Structural Basis for DNA Binding by Replication Initiator Mcm10.” Structure 16, no. 12 (Dec 10): 1892–901. 10.1016/j.str.2008.10.005.

Watase, G., H. Takisawa, and M. T. Kanemaki. 2012. “Mcm10 Plays a Role in Functioning of the Eukaryotic Replicative DNA Helicase, Cdc45-Mcm-Gins.” Curr Biol 22, no. 4 (Feb 21): 343–9. 10.1016/j.cub.2012.01.023.

Weekes, Christopher, Lia Willerding, Sanjay P. Khadayate, Korbinian Liebl, Audrey Mossler, Alex Montoya, Vanessa Rauthe, Mohammad M. Karimi, Martin Zacharias, Helle D. Ulrich, Christian Speck, and L. Maximilian Reuter. 2026. “Mechanisms of Mcm2-7 Helicase Activation and Initial DNA Melting at near Base-Pair Resolution.” bioRxiv: 2026.01.27.699880. 10.64898/2026.01.27.699880.

Weissmann, F., J. F. Greiwe, T. Puhringer, E. L. Eastwood, E. C. Couves, T. C. R. Miller, J. F. X. Diffley, and A. Costa. 2024. “Mcm Double Hexamer Loading Visualized with Human Proteins.” Nature 636, no. 8042 (Dec): 499–508. 10.1038/s41586-024-08263-6.

Wells, J. N., L. V. Edwardes, V. Leber, S. Allyjaun, M. Peach, J. Tomkins, A. Kefala-Stavridi, S. V. Faull, R. Aramayo, C. M. Pestana, L. Ranjha, and C. Speck. 2025. “Reconstitution of Human DNA Licensing and the Structural and Functional Analysis of Key Intermediates.” Nat Commun 16, no. 1 (Jan 8): 478. 10.1038/s41467-024-55772-z.

Westhorpe, R., J. J. Roske, and J. T. P. Yeeles. 2024. “Mechanisms Controlling Replication Fork Stalling and Collapse at Topoisomerase 1 Cleavage Complexes.” Mol Cell 84, no. 18 (Sep 19): 3469–3481 e7. 10.1016/j.molcel.2024.08.004.

Xia, Y., R. Sonneville, M. Jenkyn-Bedford, L. Ji, C. Alabert, Y. Hong, J. T. P. Yeeles, and K. P. M. Labib. 2023. “Dnsn-1 Recruits Gins for Cmg Helicase Assembly during DNA Replication Initiation in Caenorhabditis Elegans.” Science 381, no. 6664 (Sep 22): eadi4932. 10.1126/science.adi4932.

Yang, R., O. Hunker, M. Wise, and F. Bleichert. 2024. “Multiple Mechanisms for Licensing Human Replication Origins.” Nature 636, no. 8042 (Dec): 488–498. 10.1038/s41586-024-08237-8.

Yeeles, J. T., T. D. Deegan, A. Janska, A. Early, and J. F. Diffley. 2015. “Regulated Eukaryotic DNA Replication Origin Firing with Purified Proteins.” Nature 519, no. 7544 (Mar 26): 431–5. 10.1038/nature14285.

Yeeles, J. T. P., A. Janska, A. Early, and J. F. X. Diffley. 2017. “How the Eukaryotic Replisome Achieves Rapid and Efficient DNA Replication.” Mol Cell 65, no. 1 (Jan 5): 105–116. 10.1016/j.molcel.2016.11.017.

Zhao, X., J. Wang, D. Jin, J. Cheng, H. Chen, Z. Li, Y. Wang, H. Lou, J. K. Zhu, X. Du, and Z. Gong. 2023. “Atmcm10 Promotes DNA Replication-Coupled Nucleosome Assembly in Arabidopsis.” J Integr Plant Biol 65, no. 1 (Jan): 203–222. 10.1111/jipb.13438.

Zheng, S. Q., E. Palovcak, J. P. Armache, K. A. Verba, Y. Cheng, and D. A. Agard. 2017. “Motioncor2: Anisotropic Correction of Beam-Induced Motion for Improved Cryo-Electron Microscopy.” Nat Methods 14, no. 4 (Apr): 331–332. 10.1038/nmeth.4193.

