## Supplementary material for "MCM10 targets CMG dimers via a conserved mechanism for synchronized helicase activation": Methods and Supplementary Data

### Supplementary Figures

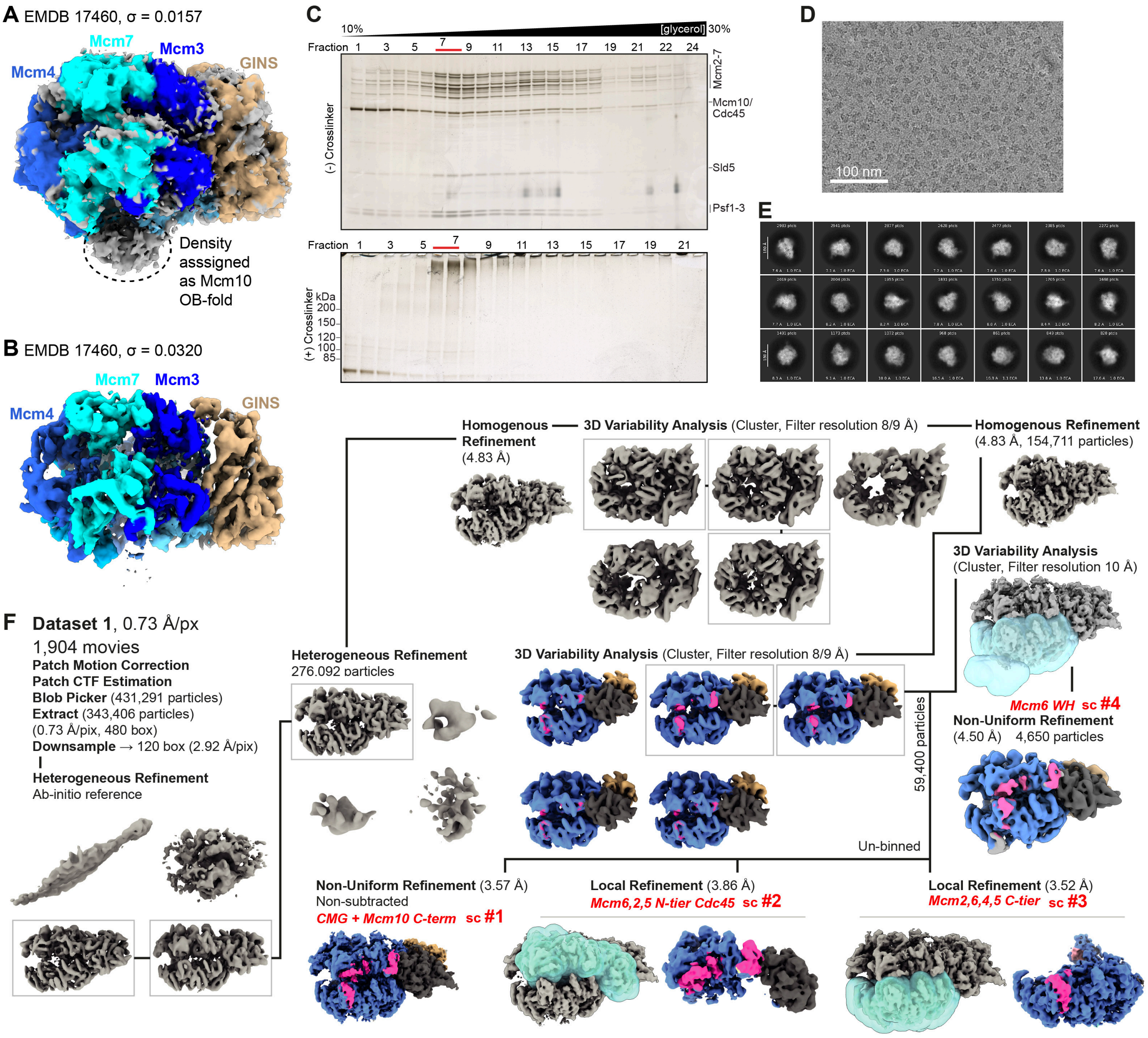


**Supplementary Fig. 1 | Analysis of published cryo-EM density assigned to *S. cerevisiae* MCM10 and cryo-EM processing for the $_{Sc}$CMG-fork DNA-Mcm10 complex**. (**A**) Cryo-EM density for *S. cerevisiae* CMG (EMDB-17460) displayed at a map threshold of σ = 0.0157 and coloured according to chain occupancy from docked PDB model 8P63(Henrikus, et al. 2024). Density assigned to the Mcm10 OB-fold is indicated by a dashed circle(Henrikus, et al. 2024). (**B**) As in (**A**) but displayed at a higher threshold (σ = 0.0320) at which the density attributed to the Mcm10 OB-fold is no longer visible. (**C**) Silver-stained SDS-PAGE analysis of fractions from 10-30% glycerol gradients prepared without crosslinker (top; 90 µl fractions) or with crosslinker (bottom; 100 µl fractions). Fractions used for cryo-EM analysis are indicated by red bars. Fractions 7-9 (top) and 6-7 (bottom) correspond to approximately the same species owing to differences in pipetting volume. (**D**) Representative cryo-EM micrograph with scale bar inset. (**E**) Representative 2D class averages with 180 Å mask diameter. (**F**) Cryo-EM image processing workflow for the CMG-fork DNA-Mcm10 complex (see Materials and Methods for details). Reported resolutions were determined using the FSC = 0.143 criterion. Final reconstructions used in modelling and deposited in the EMDB are indicated by red text labels and coloured according to protein occupancy.

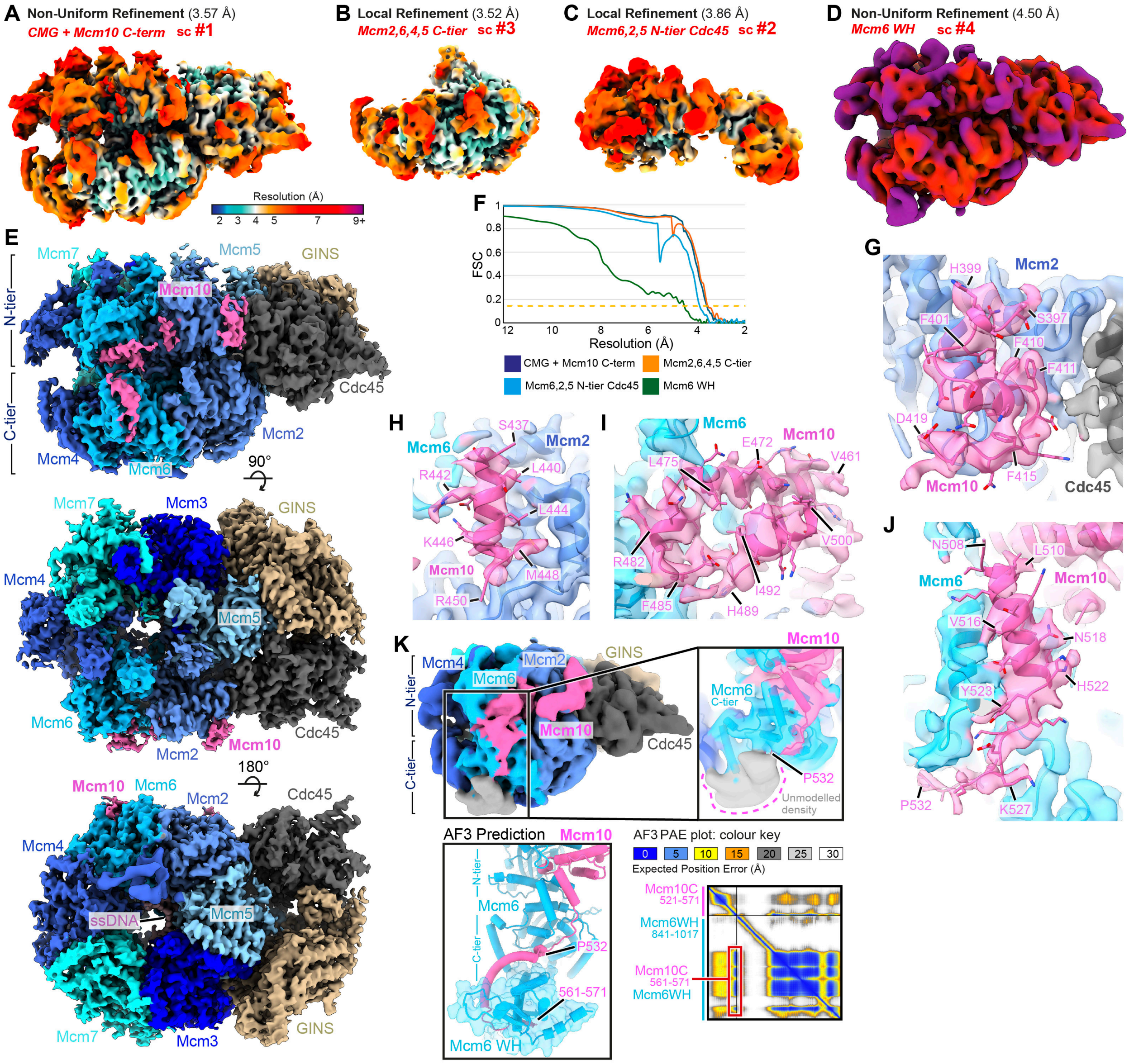


**Supplementary Fig. 2 | Supporting data for the $_{sc}$CMG-fork DNA-Mcm10 complex** (**A-D**) Cryo-EM maps coloured according to local resolution using the key in (**A**). (**A**) Consensus refinement cryo-EM density map (sc #1). (**B**) Local refinement of the Mcm2, Mcm6, Mcm4 and Mcm5 C-tier region (sc #3). (**C**) Local refinement of the Mcm6, Mcm2 and Mcm5 N-tier region together with Cdc45 (sc #2). (**D**) Consensus refinement cryo-EM density map for a particle subset displaying additional density at the base of the Mcm6 C-tier of a size consistent with a winged-helix domain (sc #4). (**E**) Three views of a composite cryo-EM density reconstruction assembled from maps in (**A-C**) and coloured according to protein occupancy. (**F**) Fourier shell correlation (FSC) curves for maps in (**A-D**). Dashed line indicates the FSC = 0.143 criterion. (**G-K**) Focused views of Mcm10-CMG interaction interfaces with cryo-EM density transparent and coloured according to protein occupancy, with docked atomic model and selected side chains annotated. (**G**) Mcm10 residues 397-419 interface with the Mcm2 N-terminal helical domain. (**H**) Mcm10 residues 437-450 interface with the Mcm2 N-terminal helical domain. **i**, Mcm10 residues 461-500 interface between the Mcm6 OB-fold and helical domain. (**J**) Mcm10 residues 508-532 interface with the AAA+ ATPase domain of the Mcm6 C-tier. (**K**) Top left, consensus refinement showing additional Mcm10-dependent density at the base of Mcm6 consistent with a winged-helix domain. Top right, enlarged view with transparent density and docked atomic model indicating the last modelled Mcm10 residue (P532). Bottom, AlphaFold3 structure prediction and predicted aligned error (PAE) plot showing the Mcm6 winged-helix engaging the Mcm10 C-terminus (residues 561-571) in a position consistent with the unmodelled density.

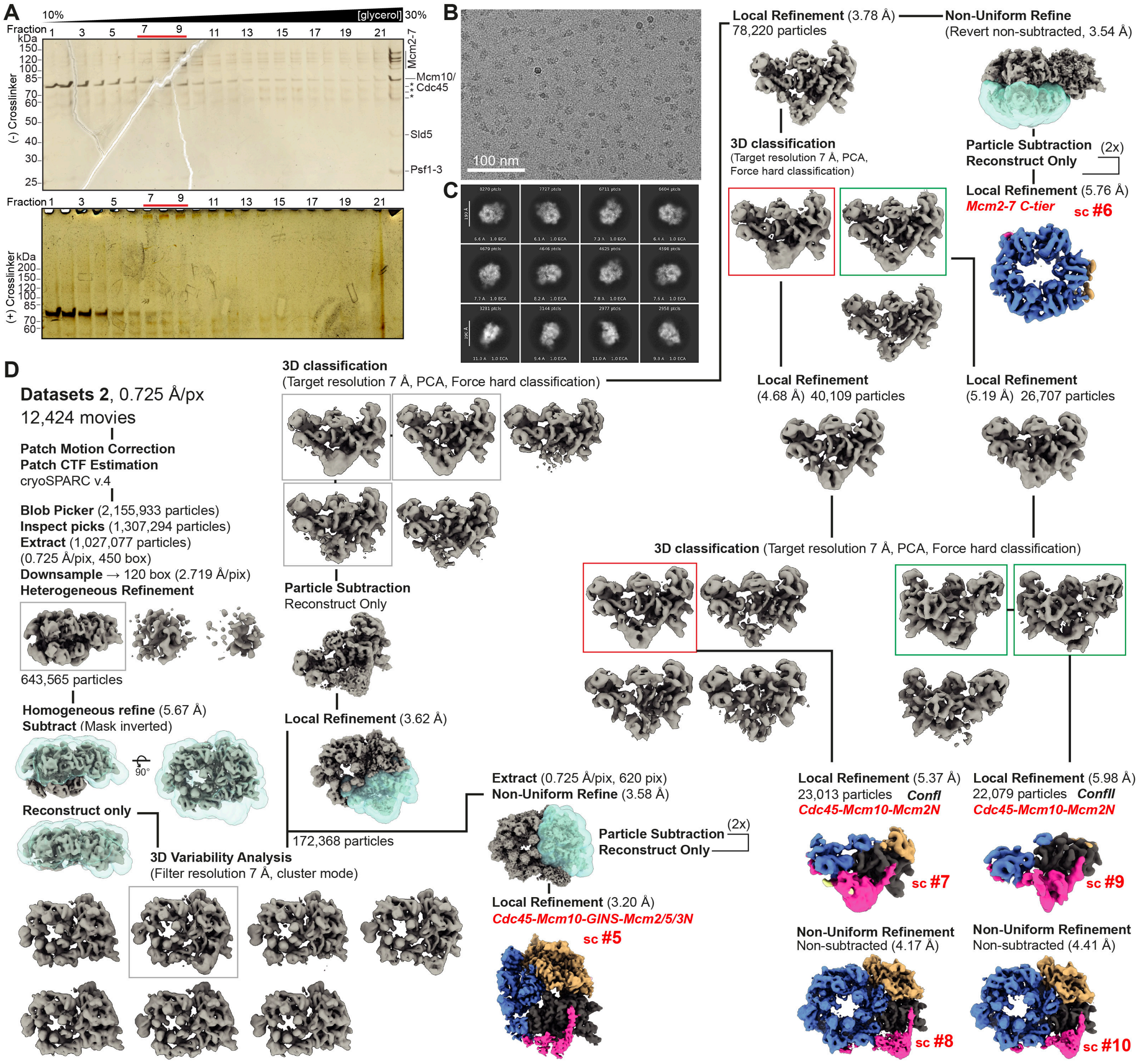


**Supplementary Fig. 3 | Cryo-EM sample preparation and processing data for the $_{sc}$CMG-Mcm10 complex.** (**A**) Silver-stained SDS-PAGE analysis of 100 µl fractions from 10-30% glycerol gradients prepared without crosslinker (top) or with crosslinker (bottom). Fractions 7-9 were used for cryo-EM analysis and indicated by red bars. (**B**) Representative cryo-EM micrograph with scale bar inset. (**C**) Representative 2D class averages with 190 Å mask diameter. (**D**) Cryo-EM image processing workflow for the CMG-Mcm10 complex (see Materials and Methods for details). Reported resolutions were determined using the FSC = 0.143 criterion. Final reconstructions used in modelling are indicated by red text labels and coloured according to protein occupancy.

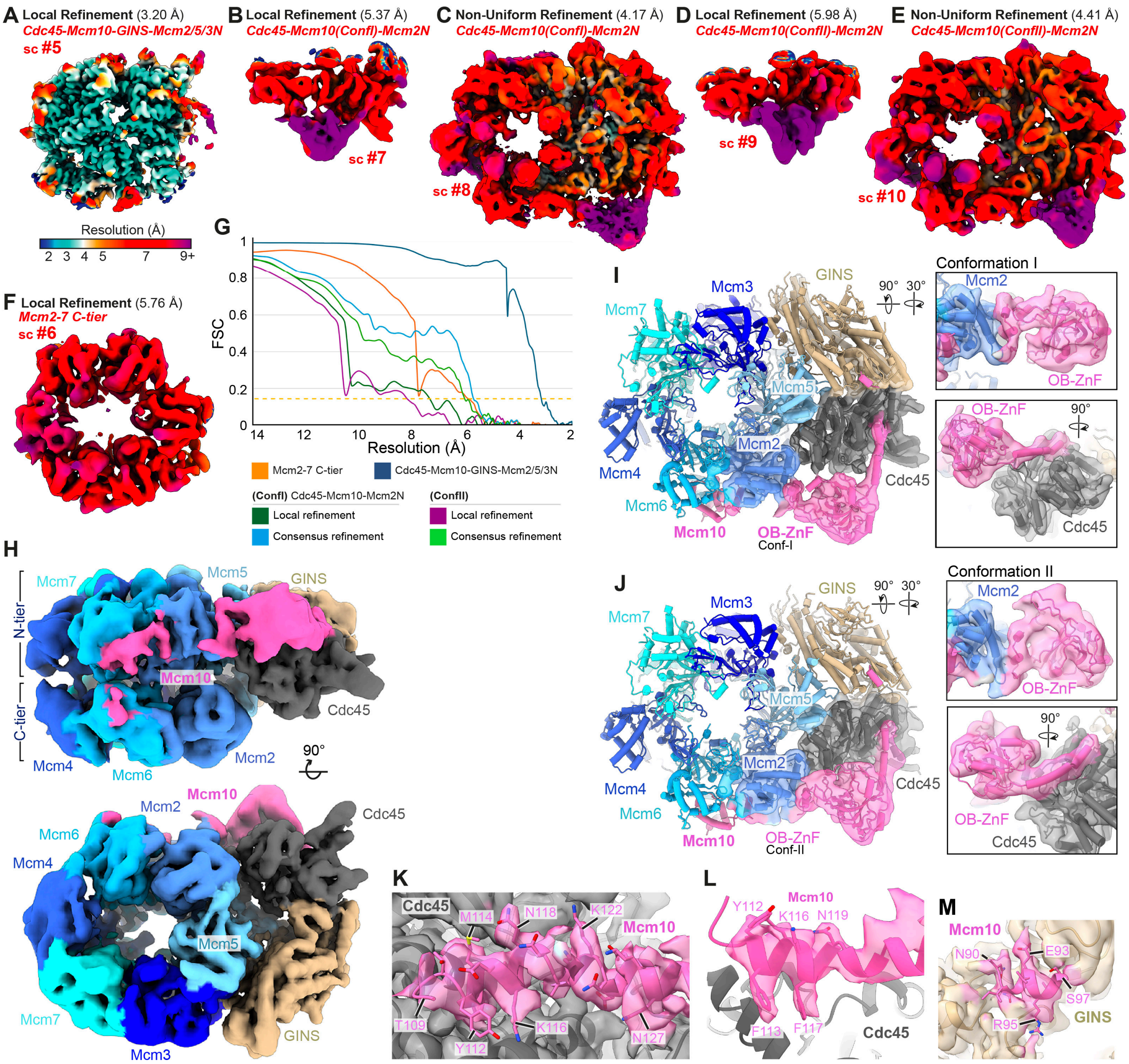


**Supplementary Fig. 4 | Supporting data for the $_{sc}$CMG-Mcm10 complex.** (**A-F**) Cryo-EM maps coloured according to local resolution using the key in (**A**). (**A**) Local refinement of the Cdc45-Mcm10-GINS-Mcm2/5/3 N-tier region (sc #5). (**B**) Local refinement of the Cdc45-Mcm10-Mcm2 N-tier region with the Mcm10 OB-fold ZnF in conformation I (sc #7). (**C**) Consensus cryo-EM density map for particles from (**B**) (sc #8). (**D**) Local refinement of the Cdc45-Mcm10-Mcm2 N-tier region with the Mcm10 OB-fold ZnF in conformation II (sc #9). (**E**) Consensus cryo-EM density map for particles from (**D**) (sc #10). (**F**) Local refinement of the Mcm2-7 C-tier ring (sc #6). (**G**) Fourier shell correlation (FSC) curves for consensus and local refinements. The dashed line indicates the FSC = 0.143 criterion. (**H**) Cryo-EM density for the CMG-Mcm10 complex with the Mcm10 OB-fold ZnF in conformation I, coloured according to protein occupancy. (**I**) Atomic model of CMG-Mcm10 with transparent cryo-EM density from the local refinement in (**B**) showing the Mcm10 OB-fold ZnF in conformation I, coloured according to protein occupancy. Insets show two rotated enlarged views of the OB-fold ZnF density. (**J**) As in (**I**), shown for the Mcm10 OB-fold ZnF in conformation II. (**K**) Focused view of the Mcm10 interface with Cdc45 with selected residues annotated and transparent cryo-EM density coloured according to protein occupancy. (**L**) As in (**K**), with cryo-EM density displayed within a 2 Å radius of Mcm10 residues 109-130 to highlight side-chain density for F113 and F117 mutated in the Mcm10$^{2A}$ mutant. (**M**) Focused view of Mcm10 residues 90-97 interacting with the Psf2 subunit of GINS.

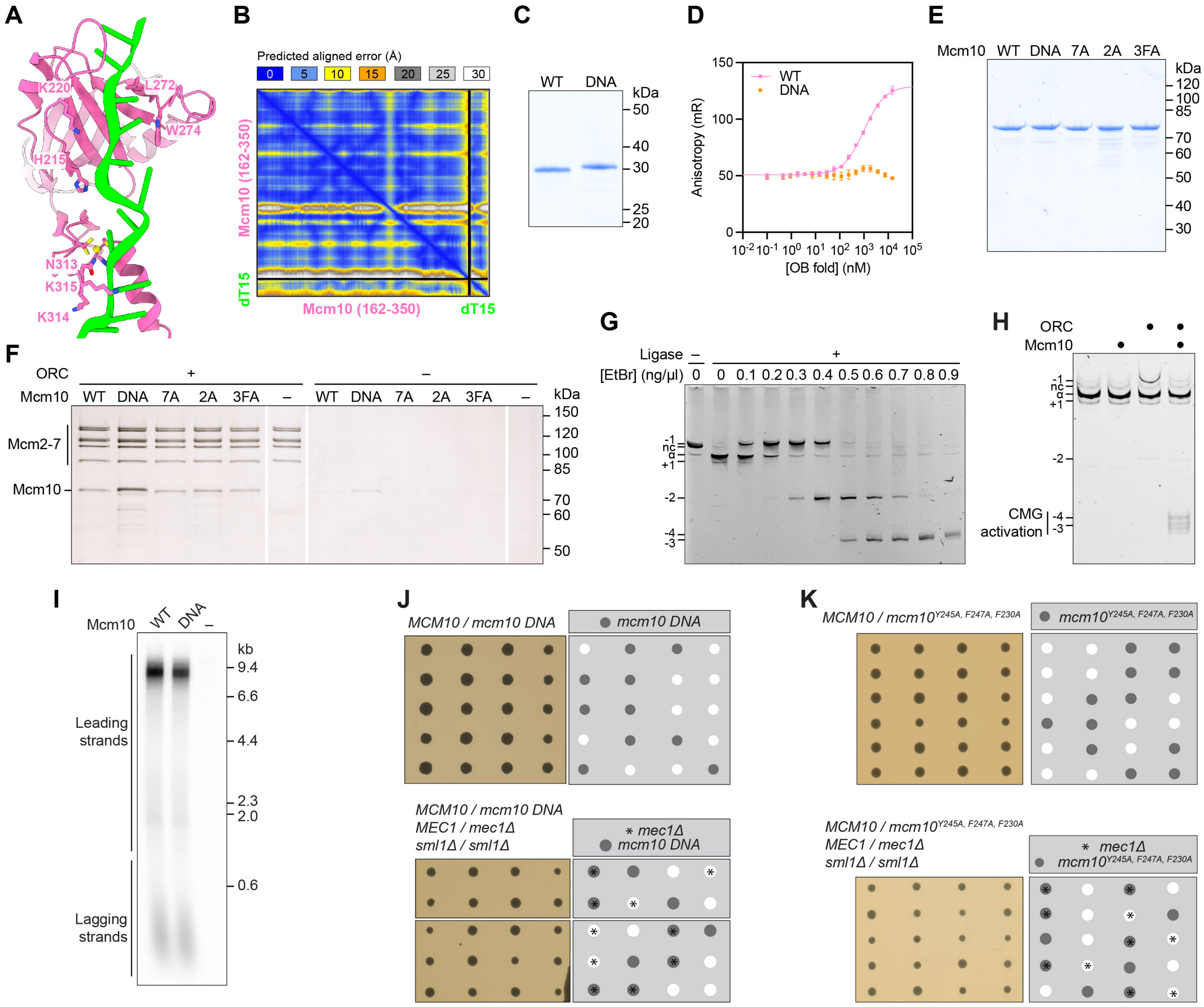


**Supplementary Fig. 5 | Supporting data for the $_{Sc}$Mcm10-Cdc45 interaction** (**A**) AlphaFold3 structure prediction of the $_{Sc}$Mcm10 OBZnF bound to ssDNA (15 nt dT). Residues mutated in $_{Sc}$Mcm10$^{DNA}$ are highlighted. (**B**) PAE plot for the AlphaFold3 structure prediction shown in (**A**). (**C**) Coomassie-stained SDS-PAGE analysis of $_{Sc}$Mcm10 OBZnF constructs used in florescence anisotropy experiments. (**D**) Florescence anisotropy measurements with 10 nM 5′-ATTO488N labelled 23nt oligonucleotide and titrating $_{Sc}$Mcm10 OBZnF (WT and DNA mutant) from 0.7 nM to 20 µM. The data is presented as mean values ± SD derived from three independent measurements. (**E**) Coomassie-stained SDS-PAGE analysis of $_{Sc}$Mcm10 mutants. (**F**) Silver-stained SDS-PAGE analysis of a double hexamer recruitment assay using purified $_{Sc}$Mcm10 mutants. ORC was omitted where indicated. (**G**) Ethidium bromide titration to define the position of mini-circle DNA topoisomers relative to ground state (α), analysed on native 3.5% polyacrylamide TBE gel. Nicked circles (nc). (**H**) Mini-circle CMG activation assay with ORC and Mcm10 omitted where indicated. (**I**) Denaturing agarose gel analysis of 12 min DNA replication reactions performed as in Fig. 2h. (**J, K**) Diploid budding yeast cells of the indicated genotype were sporulated, and the resulting tetrads dissected on YPD medium for 2 days at 30°C. Dissections that displayed abnormal segregation patterns were cropped from plate images.

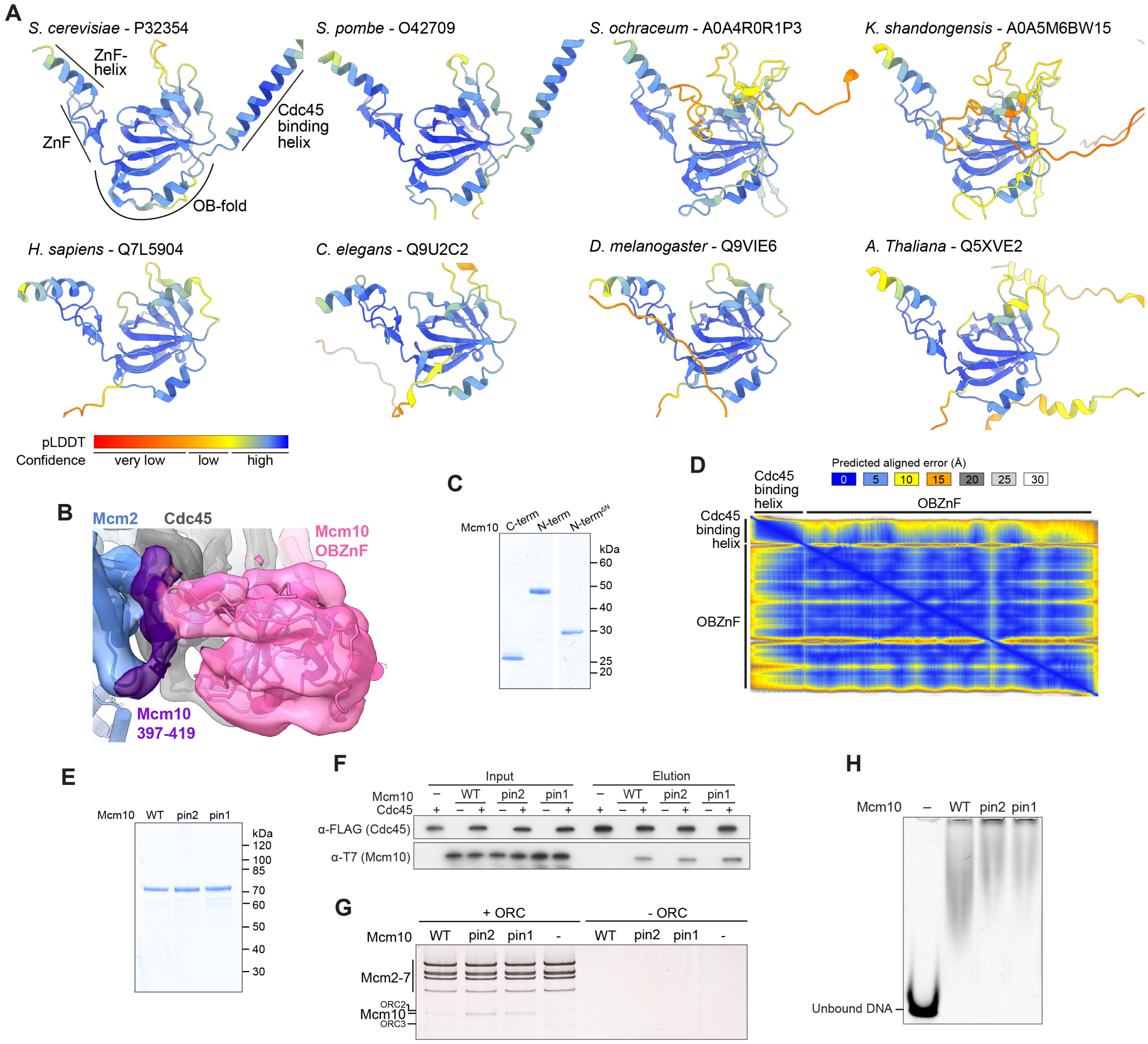


**Supplementary Fig. 6 | Supporting data for $_{Sc}$Mcm10 OB-fold positioning.** (**A**) AlphaFold3 structure predictions of the OBZnF region of MCM10 across a range of species. (**B**) Close-up view of the $_{Sc}$Mcm10 OBZnF (pink) in conformation I, highlighting its proximity to Mcm10 residues 397-419 (purple). Model fit into transparent cryo-EM density. (**C**) Coomassie-stained SDS-PAGE analysis of $_{Sc}$Mcm10 fragments. (**D**) PAE plot for the AlphaFold3 structure prediction shown in **Fig. 3C**. (**E**) Coomassie-stained SDS-PAGE analysis of $_{Sc}$Mcm10 mutants. (**F**) Immunoblot analysis of a $_{Sc}$Cdc45:Mcm10 pull-down. (**G**) Silver-stained SDS-PAGE analysis of a double hexamer recruitment assay using purified $_{Sc}$Mcm10 mutants. ORC was omitted where indicated. (**H**) Electrophoretic mobility shift assay with a 25 nt ssDNA and the indicated $_{Sc}$Mcm10 proteins.

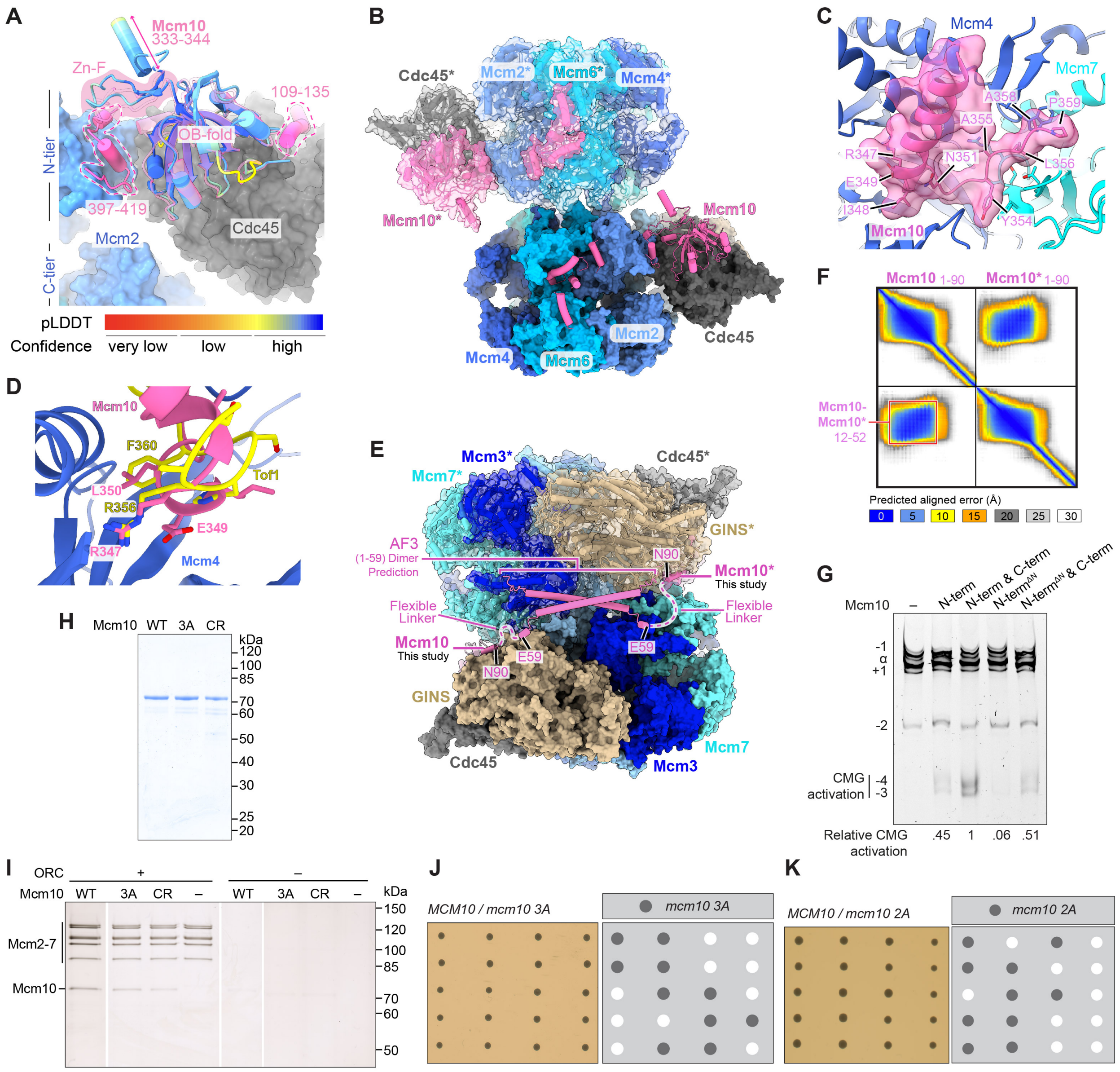


**Supplementary Fig. 7 | $_{c}$Mcm10 targets dimeric $_{Sc}$CMG complexes.** (**A**) Model of $_{Sc}$CMG-Mcm10 with the $_{Sc}$Mcm10 OBZnF domain in conformation I. Cdc45 and Mcm2 are displayed as a surface, with Mcm10 shown as a model with the OBZnF domain transparent. The AlphaFold3 structure prediction of the $_{Sc}$Mcm10 OBZnF domain (residues 138–346) is aligned to the OBZnF and coloured by pLDDT. (**B**) Two copies of the $_{Sc}$CMG:Mcm10 model aligned on a $_{Sc}$CMG-dimer structure (PDB: 7Z13). Subunits from the second $_{Sc}$CMG complex (top) are shown in cartoon representation with a transparent surface and are denoted with a *. (**C**) AlphaFold3 structure prediction showing an interaction between $_{Sc}$Mcm10 and $_{Sc}$Mcm4. Residue side chains involved in protein-protein interactions are highlighted. (**D**) Overlay of the AlphaFold3 structure prediction of the $_{Sc}$Mcm10-$_{Sc}$Mcm4 interface with the structure of $_{Sc}$Tof1 bound to $_{Sc}$Mcm4 (not shown) (PDB 6SKL) aligned on the Mcm4 N-tier. Selected residues involved in protein-protein interactions are highlighted. (**E**) Two copies of $_{Sc}$CMG-Mcm10 aligned to the DONSON-mediated *X. laevis* CMG dimer (PDB 8Q6O) coloured as in (**B**). An AlphaFold3 structure prediction of an N-terminal $_{Sc}$Mcm10 coiled-coil dimer (residues 1–59) was manually positioned adjacent to the next modelled residue (N90) to illustrate how the $_{Sc}$Mcm10 N-terminus is oriented to mediate dimerisation. (**F**) PAE plot coloured according to inset key, for the AlphaFold3 structure prediction shown in **(E)** of two copies of the Mcm10 N-terminus (residues 1-90). Boxed residues indicate the approximate boundary of residues involved in dimerisation. (**G**) Mini-circle CMG activation assay with N- and C-terminal fragments of $_{Sc}$Mcm10 (500 nM) as indicated. (**H**) Coomassie-stained SDS-PAGE analysis of ${}_{Sc}$Mcm10 mutants. (**I**) Silver-stained SDS-PAGE analysis of a double hexamer recruitment assay using purified ${}_{Sc}$Mcm10 mutants. Reactions with WT and no Mcm10 are the same experiment as in **(Supplementary Fig. 5F)**. (**J, K**) Diploid budding yeast cells of the indicated genotype were sporulated, and the resulting tetrads dissected on YPD medium for 2 days at 30°C.

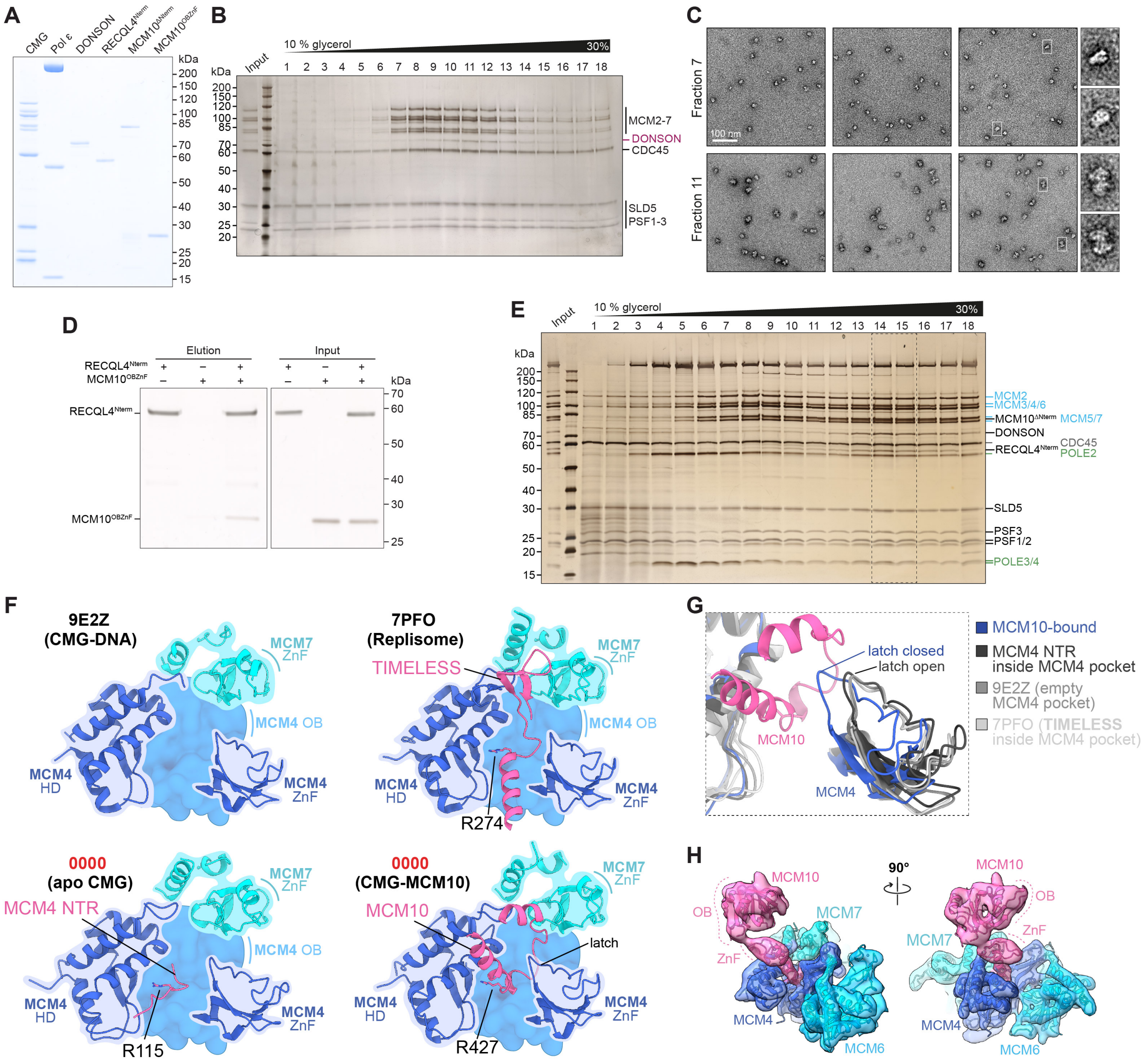


**Supplementary Fig. 8 | Supporting data for reconstitution of human complexes containing CMG, Pol ε, DONSON, MCM10 and RECQL4.** (**A**) Coomassie-stained SDS-PAGE analysis of purified human proteins. (**B**) Silver-stained SDS-PAGE analysis of fractions from a 10-30% glycerol gradient prepared without crosslinker. (**C**) Negative stain EM micrographs prepared with fractions equivalent to fractions 7 and 11 shown in (**B**), but from a 10-30% glycerol gradient containing crosslinker. White boxes outline example particles shown enlarged in the inset on the right. (**D**) Silver-stained SDS-PAGE analysis of a $_{Hs}$RECQL4$^{Nterm}$:MCM10$^{OBZnF}$ pull-down. (**E**) Silver-stained SDS-PAGE analysis of fractions from a 10-30% glycerol gradient prepared without crosslinker. Fractions identified for use in cryo-EM analysis from equivalent gradients containing crosslinker are indicated by dashed lines. (**F**) Comparison of the MCM4 pocket between different available structures of CMG: DNA-bound CMG with an empty MCM4 pocket (PDB: 9E2Z), CMG as part of the Replisome with TIMELESS bound in the MCM4 pocket (PDN: 7PFO), CMG on its own (apo CMG) with the MCM4 N-terminal region (NTR) bound in the MCM4 pocket, and MCM10-bound CMG. The different binding partners share an arginine side chain, indicated and shown as sticks, in a similar position within the MCM4 pocket. HD: helical domain. (**G**) Superposition of the atomic models shown in (**F**) aligned on the MCM4 OB fold domain. The MCM10-bound model is shown in colour, MCM4 of the remaining three models is shown in different tones of grey as indicated. (**H**) Recovered cryo-EM density map for the MCM10 OB fold (map Hs10) coloured by subunit and shown as semi-transparent surface together with the ribbon model for MCM10 residues 244-450, as well as N-tier domains of MCM6, MCM4 and MCM7.

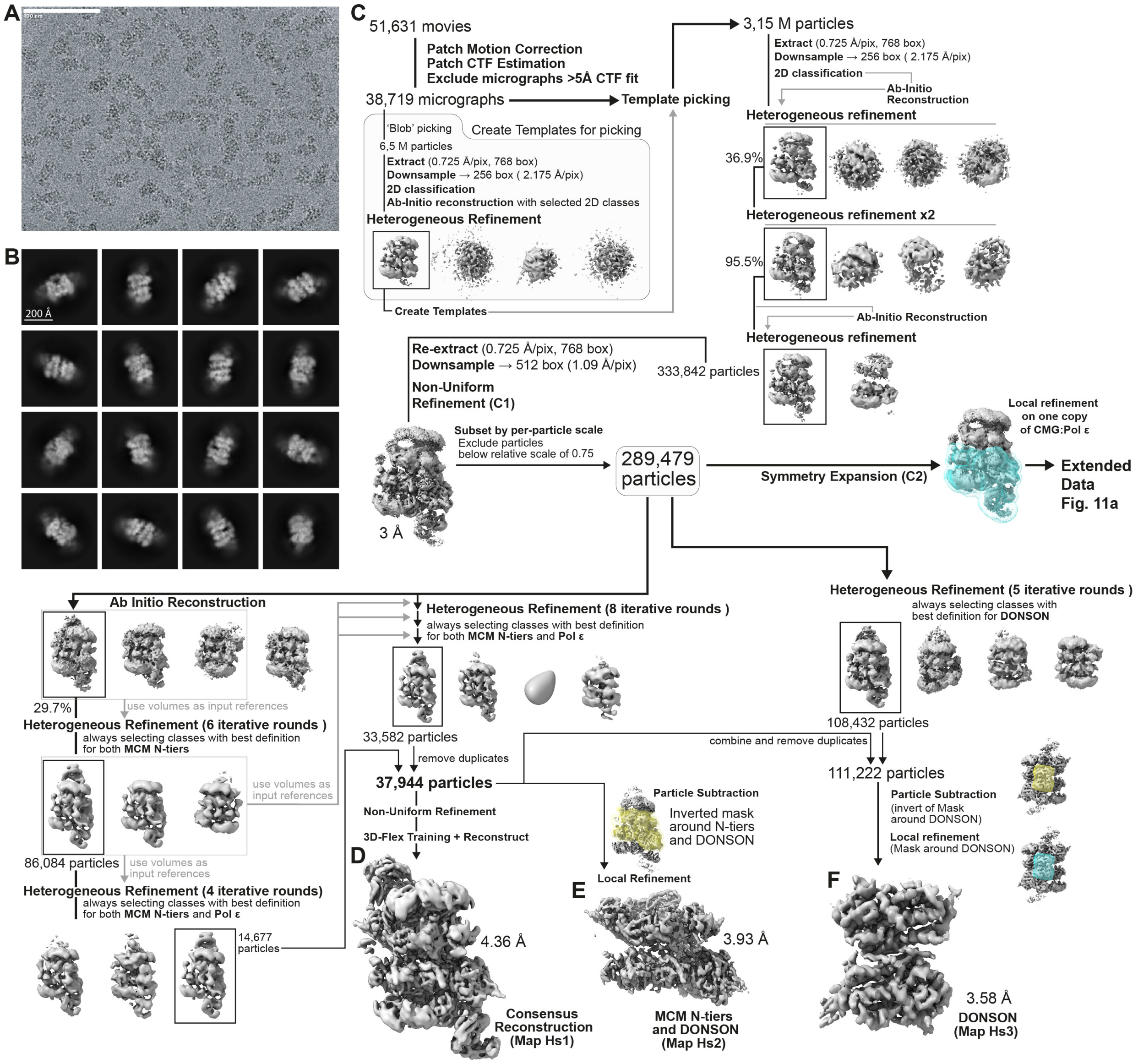


**Supplementary Fig. 9 | Cryo-EM processing pipeline for the $_{Hs}$MCM10- and $_{Hs}$RECQL4-bound dimeric DONSON:CMG:Pol ε complex.** (**A**) Sample cryo-EM micrograph after Patch Motion Correction. Scale bar, 100 nm. (**B**) Representative selected 2D class averages. Scale bar, 20 nm. (**C-F**) Cryo-EM image processing workflow for the consensus reconstruction of the $_{Hs}$MCM10- and $_{Hs}$RECQL4-bound dimeric CMG:Pol ε DONSON complex (Map Hs1, **D**), the focus map of MCM N-tiers of dimeric CMG with DONSON (Map Hs2, **E**), and the focus map of the DONSON (Map Hs3, **F**). Reported resolutions were calculated based on the FSC = 0.143 criterion. Symmetry Expansion was applied only to particles used for further processing of local regions as outlined in **Supplementary Fig. 11-12**.

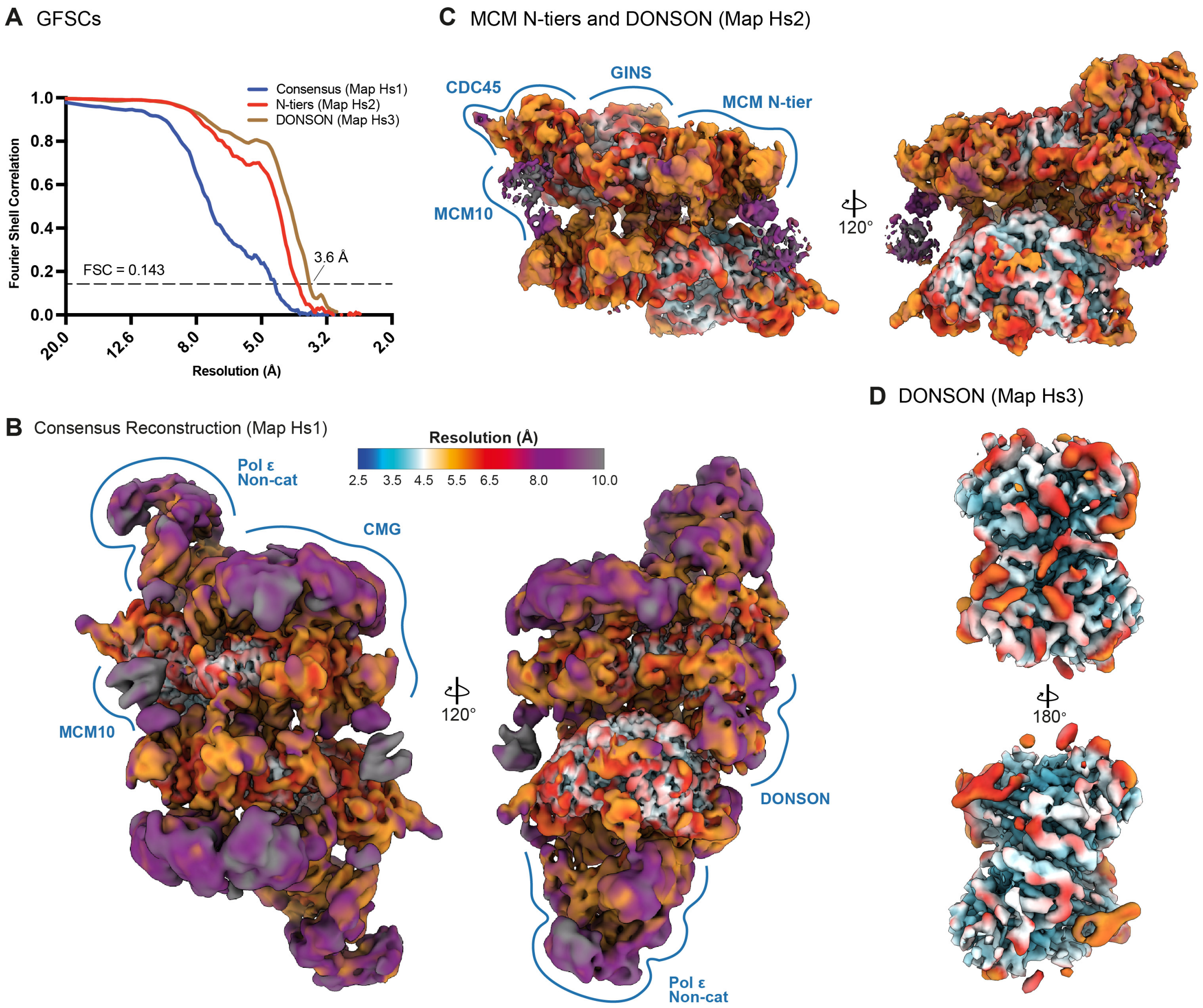


**Supplementary Fig. 10 | Local resolution of the consensus reconstruction, and local maps for MCM N-tiers and DONSON.** (**A**) Global FSC plots. (**B-D**) Maps coloured by local resolution as estimated in CryoSPARC and according to the colour scale in (**B**): Consensus reconstruction (map Hs1, **B**), MCM N-tiers of dimeric CMG together with DONSON (map Hs2, **C**) and only DONSON (map Hs3, **D**).

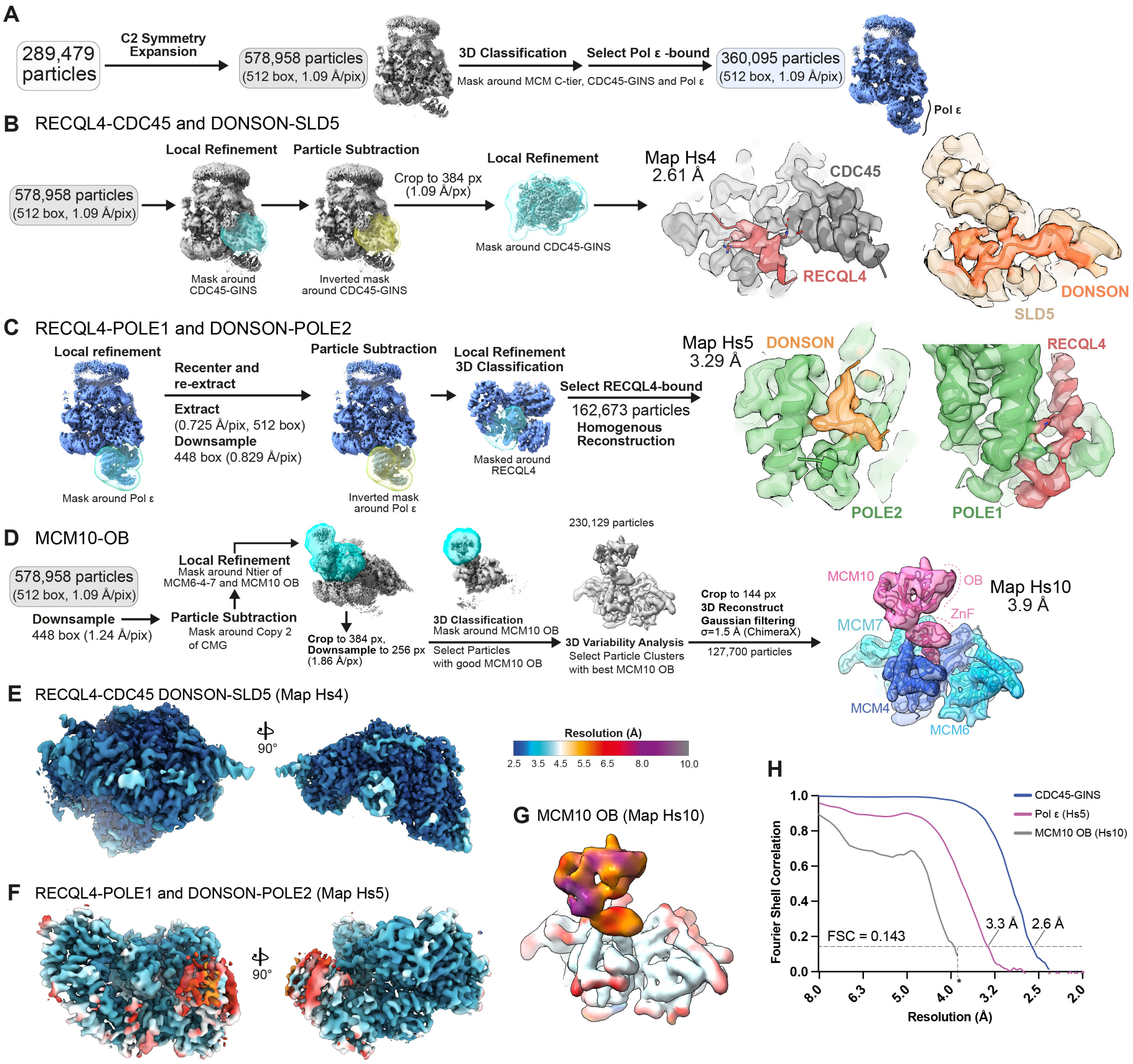


**Supplementary Fig. 11 | Local refinements for $_{Hs}$CMG:Pol ε interactions with MCM10, RECQL4 and DONSON.** (**A**) CMG particles were subclassified into Pol ε-bound particles for downstream refinements. (**B-D**) Local refinement processing pipelines for RECQL4-CDC45 and DONSON-SLD5 interactions (Map Hs4), RECQL4-POLE1 and DONSON-POLE2 interactions (Map Hs5), and MCM10 OB fold (Map Hs10). Cryo-EM density of the final maps is shown on the right as transparent surfaces together with corresponding atomic models displayed as ribbons. (**E-G**) Local resolution of map Hs4 (**E**), Hs5 (**F**) and Hs10 (**G**). Maps are coloured according to the colour scale shown. (**H**) Global FSC plots. Reported resolutions were calculated based on the FSC = 0.143 criterion. Asterisk denotes the Nyquist resolution limit imposed by the binned pixel size of map Hs10.

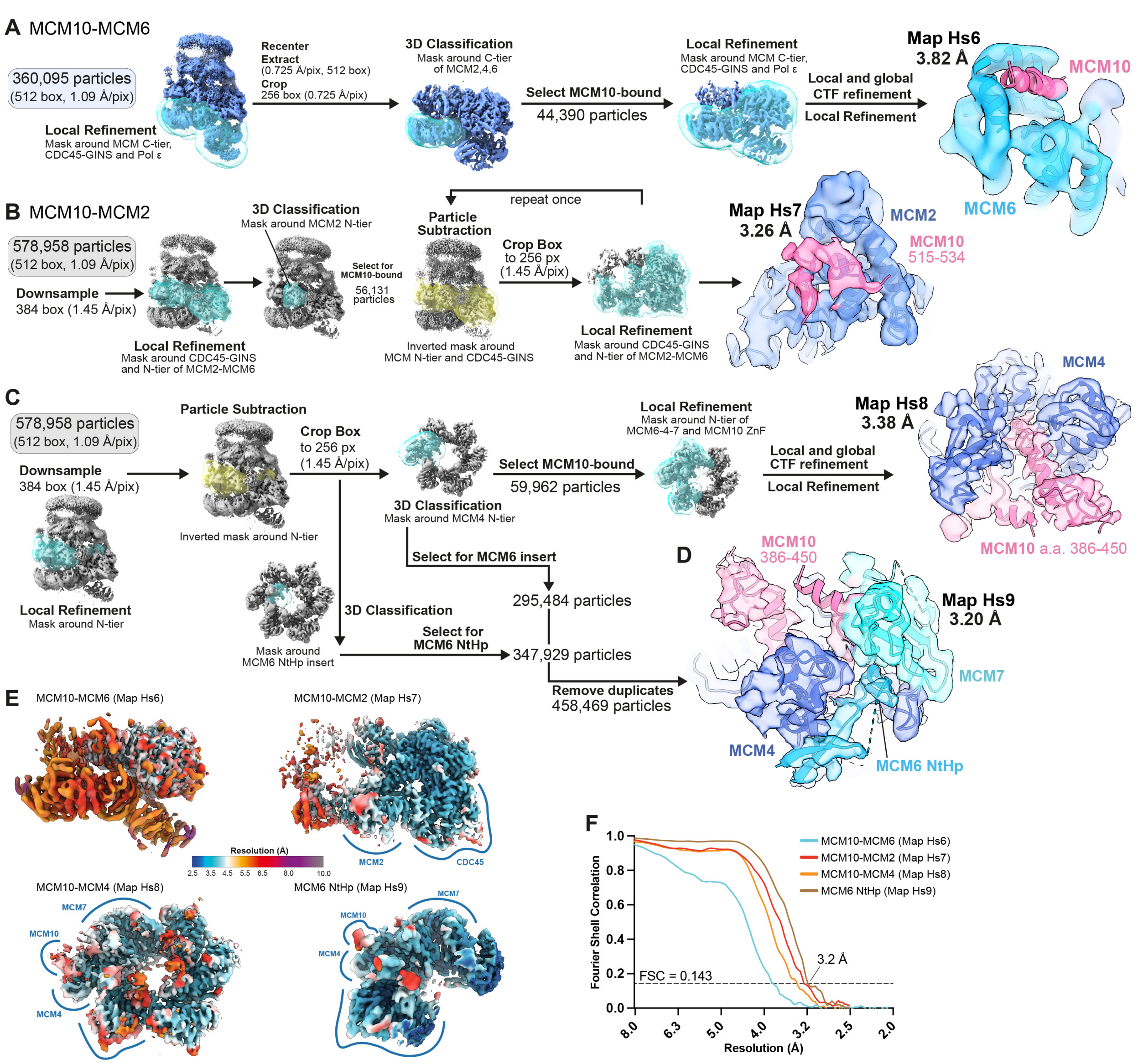


**Supplementary Fig. 12 | Local refinements for $_{Hs}$MCM10, MCM2, MCM4 and MCM6 N-tiers.** (**A-D**) Local refinement processing pipelines for the MCM10-MCM6 interaction (Map Hs6), the MCM10-MCM2 interaction (Map Hs7), MCM10-MCM4 (Map Hs8), and the MCM6 NtHp-insert interactions with MCM4 and MCM7 ZnF domains (Map Hs9). Cryo-EM density of the final maps is shown on the right as transparent surfaces together with corresponding atomic models displayed as ribbons. (**E**) Final maps Hs6-9 coloured by local resolution according to the colour scale shown. (**F**) Global FSC plots. Reported resolutions were calculated based on the FSC = 0.143 criterion.

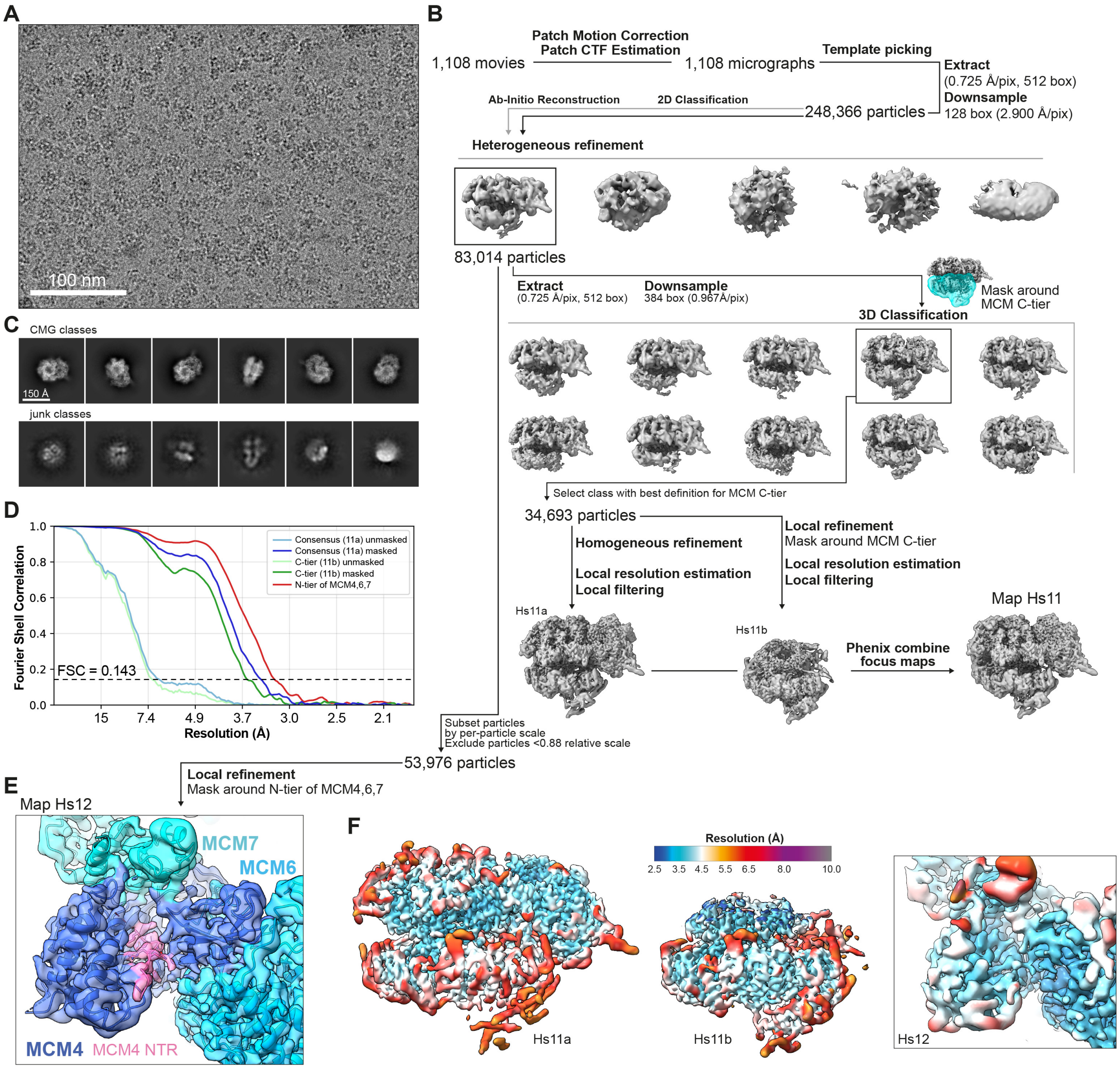


**Supplementary Fig. 13 | Cryo-EM processing pipeline for the $_{Hs}$CMG apo structure.** (**A**) Sample cryo-EM micrograph after Patch Motion Correction. Scale bar, 100 nm. (**B**) Cryo-EM image processing workflow for the $_{Hs}$CMG apo structure. (**C**) Representative selected 2D class averages for CMG or junk classes, respectively. Scale bar, 15 nm. (**D**) Global FSC plots. **(E)** Cryo-EM density of map Hs12 shown as semi-transparent surface together with the atomic models displayed as ribbons. **(F)** Maps coloured by local resolution as estimated in CryoSPARC and according to the colour scale.

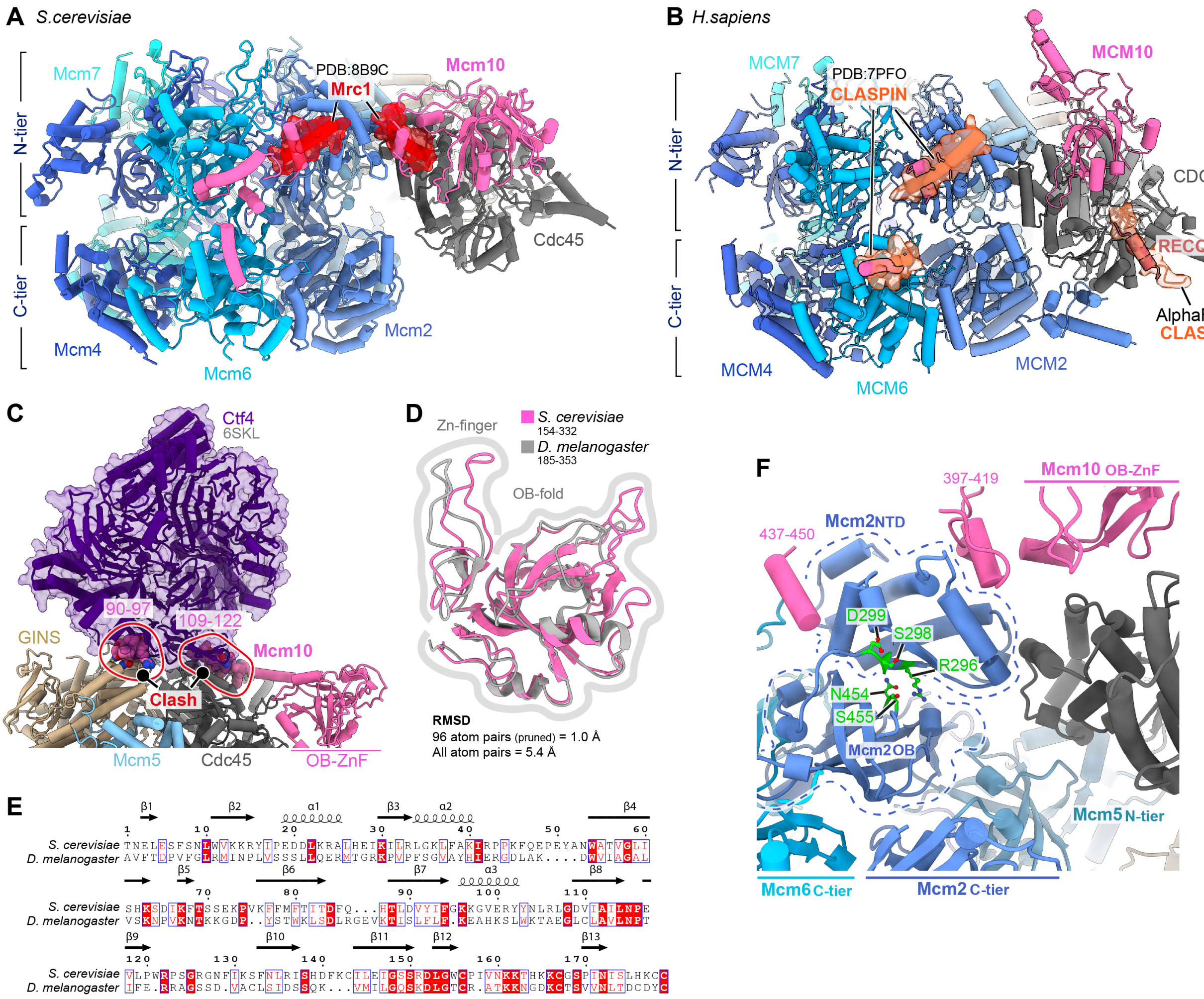


**Supplementary Fig. 14 | Structural context of Mcm10 bypass mutants and Mcm10 compatibility with CMG assembly and replisome elongation states.** (**A**) Structural overlay of the $_{Sc}$CMG-Mcm10 complex from this study with Mrc1 derived from a replisome structure bound to Pol α (PDB 8B9C). The structure was aligned via the Mcm2/6 N-tier and Cdc45. Mrc1 regions that clash with Mcm10 are rendered as transparent red surfaces. (**B**) Structural overlay of one $_{Hs}$CMG-MCM10 complex from this study with CLASPIN derived from a human replisome structure (PDB 7PFO), together with an AlphaFold3 structure prediction of CLASPIN bound to CDC45. Models were aligned via the MCM2/6 N-tier and CDC45, respectively. CLASPIN regions that clash with MCM10 are rendered as transparent red surfaces. (**C**) Structural overlay of the $_{Sc}$CMG-Mcm10 complex from this study with Ctf4 derived from a replisome structure (PDB 6SKL). The structure was aligned to Cdc45 and GINS, with only Ctf4 shown (indigo, transparent surface). Regions clashing with Mcm10 are indicated by red circles. (**D**) Structural alignment of the *S. cerevisiae* (pink) and *D. melanogaster* (grey) Mcm10 OB-fold ZnF domains with residue ranges and RMSD values indicated. (**E**) Sequence alignment of the *S. cerevisiae* and *D. melanogaster* OB-fold ZnF domains shown in (**D**) with secondary-structure assignments indicated. (**F**) Model of the $_{Sc}$CMG-Mcm10 complex highlighting Mcm10 bypass mutants in Mcm2 (green). bom-1 mutations (S455E, N454E) map to the Mcm2 OB-fold interface, whereas bom-2 mutations (R296E, S298A, D299K) map to the Mcm2 N-terminal helical domain interface.

### Materials and Methods

#### *Expression plasmid construction*

An overview of expression constructs produced for this study can be found in Supplementary Table 1. All $_{Sc}$Mcm10 mutants and fragments were cloned from pET28a-Mcm10 (Yeeles, et al. 2015) for expression in *E. coli*. For expression of $_{Sc}$Mcm10 point mutants with an N-terminal 6xHis tag, site-directed mutagenesis was performed to obtain vETC13, vETC15, vETC27, vETC48, vETC86, vETC90, vJY236, and vJY239. For expression of $_{Sc}$Mcm10$^{N\text{-term}}$, Mcm10 residues 388-571 were removed and the N-terminal 6xHis tag replaced with an ALFA tag to produce vETC92. For expression of $_{Sc}$Mcm10$^{N\text{-term}\Delta N}$, Mcm10 residues 2-106 were removed from vETC92 to produce vETC99. $_{Sc}$Mcm10$^{C\text{-term}}$ was cloned by removal of Mcm10 residues 2-387 as well as the N-terminal 6xHis tag from plasmid 1883 and a C-terminal ALFA tag was introduced to produce vJY233. $_{Sc}$Mcm10$^{OB\ fold}$ (Mcm10 residues 110-377) was cloned from plasmid 1883 and a C-terminal StrepII-tag introduced to produce vETC87. $_{Sc}$Mcm10$^{OB\ fold\ DNA}$ was cloned via Gibson Assembly using vETC86 and vETC87 to produce vETC89.

For expression of $_{Hs}$DONSON, the sequence was codon optimised for expression in *S. cerevisiae* with a TEV-cleavable N-terminal 2xFLAG tag, synthesised by GeneArt synthesis, and subcloned into pRS303 expression vector to produce vEF32.

For expression of $_{Hs}$MCM10$^{\Delta Nterm}$, the sequence for full length $_{Hs}$MCM10 was codon optimized for expression in *E. coli*, synthesized by GeneArt synthesis, and subcloned into pET28a with a TEV-cleavable N-terminal 2xStrep tag and a TEV-cleavable C-terminal 2xFLAG tag. Residues 1-229 of MCM10 were removed to produce pJR01. For expression of $_{Hs}$MCM10$^{OBZnF}$, a stop codon was introduced after residue 450 by site-directed mutagenesis of pJR01 to produce pJR02.

For expression of $_{Hs}$RECQL4$^{Nterm}$, the sequence for full-length RECQL4 was codon optimised for expression in *S. frugiperda* and synthesised by GeneArt synthesis. Residues 1-440 were subsequently cloned into pET28a along with a C-terminal 2xFLAG tag for expression in *E. coli.*

#### *S. cerevisiae strain construction*

An overview of *S. cerevisiae* strains (W303) produced for this study can be found in Supplementary Table 2. For expression of $_{Hs}$DONSON, plasmid vEF32 was transformed into *S. cerevisiae* yJF1 (Frigola et al. 2013) to produce yEF32.

For meiotic progeny analysis, Mcm10 mutants were subcloned from pET28a plasmids into pFA6a-URA3 plasmids (see Supplementary Table 1) such that the *URA3* cassette was inserted downstream of the *MCM10* gene. *MCM10-URA3* was then amplified using primers ETC_oligo_154 and ETC_oligo_155 (see Supplementary Table 3) with 57 bp homology overhangs for integration into the *MCM10* locus and transformed into diploid yeast using standard yeast genetics techniques. Correct integration and absence of additional mutations were confirmed by sequencing (Source Bioscience) of PCR products generated from yeast genomic DNA using primers ETC_oligo_173 and ETC_oligo_174 (see Supplementary Table 3).

#### *Protein expression and purification*

*S. cerevisiae* DNA replication proteins were purified as described previously (Yeeles, et al. 2015; Yeeles, et al. 2017; Taylor and Yeeles 2018; Baretić, et al. 2020). An overview of protein purification strategies can be found in Supplementary Table 4. $_{Hs}$CMG and $_{Hs}$Pol ε were purified as described previously (Jones, et al. 2021; Roske and Yeeles 2024).

*$_{Sc}$Mcm10 purification*
All full-length $_{Sc}$Mcm10 mutants were expressed and purified as previously described for wildtype $_{Sc}$Mcm10 (Yeeles, et al. 2015).

Truncated $_{Sc}$Mcm10 constructs including OB fold constructs, N-terminal fragments, and C-terminal fragment were expressed as for $_{Sc}$Mcm10 (Yeeles, et al. 2015). Rosetta2 (DE3), transformed with the relevant pET28a plasmids (Supplementary Table 1), were grown in a 2 L culture at 37°C to an $OD_{600}$ of 0.6. Protein expression was induced by addition of IPTG to 1 mM and growth continued for 3 hr at 25°C. Cells were harvested by centrifugation, washed in PBS and resuspended in 25 mM Tris-HCl pH 7.6, 10% glycerol, 0.01% NP-40-S, 500 mM NaCl (Buffer M + 500 mM NaCl) + protease inhibitors (cOmplete EDTA-free). Cells were lysed by sonication (40%, 2s on/2s off, total 2 min) and the debris removed by centrifugation (257,000 g, 4°C, 30 min).

For StrepII-tagged $_{Sc}$Mcm10$^{OB\ fold}$ constructs, cleared lysate was incubated with 2 ml Strep-Tactin XT 4Flow resin for 1 hr at 4°C. The resin was collected and washed with 20 column volumes (CV) Buffer M + 500 mM NaCl. The resin was resuspended in 10 CV Buffer M + 500 mM NaCl + 10 mM $Mg(OAc)_2$ + 1 mM ATP and incubated for 10 min at 4°C. The flow-through was discarded and the column washed with 20-40 CV Buffer M + 200 mM NaCl. Protein was eluted in 20 CV Buffer M + 200 mM NaCl + 50 mM biotin. The eluate was concentrated and separated on a Superdex 200 Increase 10/300 GL column equilibrated in 25 mM HEPES-KOH pH 7.6, 10% glycerol, 0.01% v/v NP-40-S, 1 mM EDTA, 1 mM DTT, 200 mM potassium glutamate (Buffer H + 200 mM potassium glutamate).

For ALFA-tagged $_{Sc}$Mcm10$^{N\text{-}term}$, $_{Sc}$Mcm10$^{N\text{-}term\Delta N}$, $_{Sc}$Mcm10$^{C\text{-}term}$, cleared lysate was incubated with 0.5 ml ALFA Selector CE resin for 1 hr at 4°C. Resin was collected and washed in 50 CV Buffer M + 500 mM NaCl followed by 20 CV Buffer M + 200 mM NaCl. Protein was eluted in 4 CV Buffer M + 200 mM NaCl + 0.4 mg/ml ALFA elution peptide (NanoTag Biotechnologies). The eluate was applied to a MonoS 5/50 GL column (GE Healthcare) equilibrated in Buffer M + 200 mM NaCl. The column was washed with 5 CV Buffer M + 200 mM NaCl and protein eluted with a 20 CV gradient from 200 mM NaCl to 1 M NaCl in Buffer M. Peak fractions were dialysed against Buffer H + 200 mM potassium glutamate for $_{Sc}$Mcm10$^{N\text{-}term}$ and $_{Sc}$Mcm10$^{N\text{-}term\Delta N}$ or Buffer H + 300 mM KOAc for $_{Sc}$Mcm10$^{C\text{-}term}$.

*$_{Hs}$DONSON purification*
$_{Hs}$DONSON was purified from *S. cerevisiae* yEF32 as follows. A 10 L culture of cells was grown at 30°C to $3x10^7$ cells/ml in YEP supplemented with 2% raffinose. Expression was induced by addition of galactose to 2% and cell growth continued for 3 hr at 30°C. Cells were harvested, resuspended in Buffer D (25 mM HEPES-KOH pH7.6, 400 mM NaCl, 10% glycerol, 0.005% Tween20, 0.5 mM TCEP) + protease inhibitors (cOmplete EDTA-free, one tablet per 50 ml buffer), frozen dropwise in liquid nitrogen and crushed manually using pestle and mortar cooled with liquid nitrogen. Cell powder was resuspended 1:1 in Buffer D + protease inhibitors and cell lysate cleared by centrifugation (235,000 g, 4°C, 45 min). Cleared lysate was filtered and incubated with Anti-FLAG M2 affinity gel (1 ml bed volume per 2.5 L original culture volume) at 4°C for 90 min. The resin was collected and washed with 50 CV Buffer D before washing with 5 CV Buffer D + 5 mM $Mg(OAc)_2$ + 0.5 mM ATP during which flow was stopped for 10 min. The resin was then washed with 5 CV Buffer D before elution of the protein with 1 CV Buffer D + 0.2 mg/ml 3xFLAG peptide followed by 2 CV Buffer D + 0.1 mg/ml 3x FLAG peptide.

The eluate was concentrated to ~ 500 µl and applied to a Superdex 200 Increase 10/300 GL column equilibrated in Buffer D but 300 mM NaCl. Peak fractions were pooled.

*$_{Hs}$MCM10$^{\Delta Nterm}$ purification*

$_{Hs}$MCM10$^{\Delta Nterm}$ was overexpressed in and purified from *E. coli* as follows. A 1 L culture of Rosetta 2(DE3) cells was grown at 37°C in LB to $OD_{600}$ = 0.5, expression induced by addition of 0.5 mM IPTG and growth continued at 20°C overnight. Cells from 1 L of culture were resuspended in $_{Hs}$MCM10 lysis buffer (50 mM HEPES-KOH pH7.6, 300 mM NaCl, 10% glycerol, 0.02% NP-40-S, 0.5 mM TCEP, 50 µM ZnCl) + 1 mM EDTA + protease inhibitors (cOmplete EDTA-free) and lysed via sonication (60%, 2 sec on/ 3 sec off, 2x 1 min). The lysate was cleared by centrifugation (235,000 g, 4°C, 25 min), filtered, and incubated with Strep-Tactin XT 4Flow resin (0.5 ml bed volume) at 4°C for 15 min. The resin was collected and washed with 60CV $_{Hs}$MCM10 lysis buffer before elution of the protein with $_{Hs}$MCM10 lysis buffer + 30 mM biotin. Peak fractions were kept, side fractions were pooled and concentrated.

*$_{Hs}$MCM10$^{OBZnF}$ purification*

$_{Hs}$MCM10$^{OBZnF}$ was purified from *E. coli* as follows. A 1 L culture of Rosetta 2(DE3) cells was grown at 37°C in 2XTY to $OD_{600}$ = 0.25, expression induced by addition of 0.5 mM IPTG and growth continued at 20°C overnight. Cells from 1 L of culture were resuspended in $_{Hs}$MCM10 lysis buffer + 1 mM EDTA + protease inhibitors (cOmplete EDTA-free) and lysed via sonication (40%, 2 sec on/ 2 sec off, total 2 min). The lysate was cleared by centrifugation (235,000 g, 4°C, 25 min), filtered, and incubated with Strep-Tactin XT 4Flow resin (2 ml bed volume) at 4°C for 40 min. The resin was collected and washed with 60 CV $_{Hs}$MCM10 lysis buffer + 1 mM EDTA before washing with 5 CV $_{Hs}$MCM10 lysis buffer + 5 mM $Mg(OAc)_2$ + 0.5 mM ATP during which flow was stopped for 10 min. The resin was then washed with 5 CV $_{Hs}$MCM10 lysis buffer before elution of the protein with 5 CV $_{Hs}$MCM10 lysis buffer + 30 mM biotin.

*$_{Hs}$RECQL4$^{Nterm}$ purification*

$_{Hs}$RECQL4$^{Nterm}$ was overexpressed in Rosetta 2(DE3), induced by 0.5 mM IPTG for 3 hr at 25°C. Cells from 2 L culture were resuspended in 25 ml 50 mM Tris pH 8, 150 mM NaCl, 1 mM EDTA, 0.5 mM DTT + protease inhibitors (cOmplete EDTA-free) and lysed by addition of 0.2 mg/mL lysozyme and 0.2 % Brij-35 followed by gentle stirring for 30 min at room temperature. The lysate was cleared by centrifugation (50,000 rpm, 4°C, 30 min, Beckman Ti 70 rotor) and incubated with Anti-FLAG M2 affinity gel (800 µl suspension) at 4°C for 90 min. The resin was collected and washed with 4 CV 25 mM HEPES pH 7.6, 400 mM NaCl, 1 mM EDTA, 10 % glycerol, 0.005 % Tween 20, 0.5 mM DTT (Buffer R + 400 mM NaCl), before elution of the protein with 1 CV Buffer R + 400 mM NaCl + 0.2 mg/ml 3xFLAG peptide followed by 2 CV Buffer R + 400 mM NaCl + 0.1 mg/ml 3x FLAG peptide. The eluate was diluted with an equal volume of Buffer R and applied to a HiTrap SP FF equilibrated in Buffer R + 200 mM NaCl. The column was washed with 5 CV Buffer R + 200 mM NaCl and the protein was eluted with a 20 CV gradient from 200 mM NaCl to 1 M NaCl in Buffer R. Peak elution fractions were combined and dialysed against Buffer R + 220 mM NaCl.

***Regulated system replication reactions***

Regulated system reactions were conducted as previously described using Ahd1-linearised CsCl gradient purified ZN3 plasmid (Taylor and Yeeles 2018). MCM loading and phosphorylation were performed at 24°C for 10 min in a reaction containing 25 mM HEPES-

KOH pH 7.6, 100 mM potassium glutamate, 0.01% NP-40-S, 1 mM DTT, 10 mM $Mg(OAc)_2$, 0.1 mg/ml BSA, 40 mM KCl, 3 mM ATP, 3 nM Ahd1-linearised ZN3 DNA, 75 nM Cdt1.Mcm2-7, 40 nM Cdc6, 20 nM ORC, and 50 nM DDK. S-CDK was added to 80 nM, and incubation extended for 5 min. The loading reaction was then diluted fourfold into replication buffer containing the following components (reported in their final concentrations): 25 mM HEPES-KOH pH 7.6, 250 mM potassium glutamate, 0.01% NP-40-S, 1 mM DTT, 10 mM $Mg(OAc)_2$, 0.1 mg/ml BSA, 3 mM ATP, 200 µM C/G/UTP, 30 µM dA/dG/dC/dTTP, and 33 nM [α-$^{32}$-P]-dCTP and incubation temperature increased to 30°C. Reactions were initiated by addition of replication proteins to the following concentrations: 30 nM Dpb11, 100 nM GINS, 30 nM Cdc45, 10 nM Mcm10, 20 nM Ctf4, 20 nM Tof1-Csm3, 20 nM PCNA, 20 nM Pol ε, 10 nM Pol δ, 100 nM RPA, 20 nM Pol α-primase, 15 nM Mrc1, 12.5 nM Sld3/7, 20 nM Sld2, 20 nM RFC. Additional salt contributions from protein storage buffers ranged from approximately 35-45 mM. Aliquots were quenched with a final concentration of 50 mM EDTA after 12 min.

Unincorporated nucleotides were removed using Illustra MicroSpin G-50 columns. Samples were analysed on 0.7% alkaline agarose gels in 30 mM NaOH and 2 mM EDTA for 16 hr at 24 V. Gels were fixed with 2x washes in 5% trichloroacetic acid at 4°C and subsequently neutralised with 1 M Tris-HCl pH 8 before drying onto 3 mm chromatography paper. Dried gels were exposed on BAS-IP MS Storage Phosphor Screens and imaged using an Amersham Typhoon phosphorimager. All regulated system replication reactions were performed in triplicate.

***Preparation of mini-circle templates for CMG activation assays***

Mini-circle preparation was based on a published method (Douglas, et al. 2018) with modifications. To assemble FAM-labelled ARS306 mini-circles, a 657 bp fragment around ARS306 was PCR amplified from ZN3 (Baris et al. 2022) using oligonucleotides JY698 and JY699 (JY698 contains an internal FAM modification, Supplementary Table 3) which introduce recognition sites for SpeI and NheI respectively. Between 80-100 µg DNA was digested with 400 U SpeI and NheI at 37°C overnight and purified using QIAquick PCR Purification Kit. Between 60-90 µg DNA was then ligated overnight at 4°C at concentrations <200 ng/ml with 40 U/ml T4 DNA ligase (NEB). Ligation reactions were stopped with 25 mM EDTA and concentrated ~1000 fold through a 10 kDa cutoff spin concentrator before CsCl gradient purification to isolate covalently closed circles.

***Assigning relative supercoiling states***

The relative supercoiling state of different 657 bp mini-circle topoisomers were assigned approximately as described previously (Douglas, et al. 2018). 50 nM 657 bp DNA mini-circles were incubated at 37°C for 3 hours with 1,000 U/ml Nt.BsmAI (NEB). Nt.BsmAI was heat inactivated at 65°C for 20 min. 2.5 nM of digested DNA was ligated in the presence of the ethidium bromide concentrations indicated at room temperature overnight with 10,000 U/ml T4 DNA ligase (NEB). Ligated DNA was phenol:chloroform:isoamylalcohol extracted, ethanol precipitated, and the DNA pellet was resuspended in 1x TE prior to analysis on a 3.5% polyacrylamide TBE gel for 5 hr at 60 V. Final DNA circles are increasingly negatively supercoiled as ethidium bromide concentration is increased in the ligation step. Topoisomers were assigned relative to the ground state (α, the most prevalent topoisomer when ethidium bromide was omitted) by tracking the order in which bands peaked as the ethidium bromide concentration increased.

#### *Mini-circle CMG activation assays*

CMG activation assays were performed approximately as described previously (Douglas, et al. 2018). Reactions containing 25 mM HEPES-KOH pH 7.6, 100 mM potassium glutamate, 10 mM $Mg(OAc)_2$, 0.01% NP-40-S, 1 mM DTT, 0.1 mg/ml BSA, 3 mM ATP, 12 nM 657 bp DNA, 20 nM TopoI, 100 nM Cdt1.Mcm2-7, 20 nM ORC, 40 nM Cdc6, 200 nM DDK, were incubated at 30°C for 30 min. S-CDK was added to 100 nM and incubation continued for a further 10 min. Reactions were diluted twofold into replication buffer containing the following components (reported in their final concentrations): 25 mM HEPES-KOH pH 7.6, 160 mM potassium glutamate, 0.01% NP-40-S, 1 mM DTT, 10 mM $Mg(OAc)_2$, 0.1 mg/ml BSA, 5.5 mM ATP. Reactions were initiated by addition of replication proteins to the following concentrations: 30 nM Dpb11, 200 nM GINS, 30 nM Cdc45, 20 nM Pol ε, 10 nM TopoI, 25 nM Sld3/7, 50 nM Sld2, and 15 nM Mcm10 (or otherwise stated). Reactions were incubated at 30°C for 25 min then quenched with 45 mM EDTA before addition of 0.1 mg/ml Proteinase K and 0.2% SDS and incubation at 37°C for 20 min. Samples were analysed on native 3.5% polyacrylamide TBE gels at 34 V for 16.5 hr. Fluorescence was detected using Amersham Typhoon imager (1.1.0.7) (Cytiva). To quantify CMG activation, lane profiles were taken in ImageJ and the proportion of -3 and -4 topoisomer signal relative to total lane signal was calculated. Proportions were normalized to reactions containing WT Mcm10 or, where a WT Mcm10 control was absent, to the reaction with the most CMG activation. All mini-circle CMG activation assays were performed in triplicate.

#### *Double hexamer recruitment assays*

$_{Sc}$MCMs were loaded onto 1kb linear ARS306 DNA coupled to Dynabead M280 Strepatavidin magnetic beads by incubation of 5 µl DNA-coupled beads in a reaction containing loading buffer (25 mM HEPES-KOH pH 7.6, 10 mM $Mg(OAc)_2$, 0.02% NP-40-S, 1 mM DTT), 100 mM potassium glutamate, 40 mM KCl, 3 mM ATP, 75 nM Cdt1.Mcm2-7, 20 nM ORC, and 40 nM Cdc6 for 30 min at 30°C, 1250 rpm. Beads were washed with 2x 100 µl resuspension in recruitment buffer (loading buffer + 10% glycerol) + 500 mM NaCl followed by 2x 100 µl recruitment buffer + 300 mM potassium glutamate. Beads were then incubated at 30°C, 1250 rpm for 10 min in reactions containing recruitment buffer, 300 mM potassium glutamate, 1 mM ATP, and 20 nM various Mcm10. Beads were washed with 3x 100 µl resuspension in recruitment buffer + 300 mM potassium glutamate before incubation in recruitment buffer + 300 mM NaOAc + 2 mM $CaCl_2$ + 2000 U MNase for 5 min at 30°C, 1250 rpm. Elutions were analysed on 4-12% Bis-Tris SDS-PAGE gels and silver-stained. All double hexamer recruitment assays were performed in triplicate.

#### *Pull-downs*

To detect interactions between $_{Sc}$Cdc45 and $_{Sc}$Mcm10, 20 µl reactions containing interaction buffer (25 mM HEPES-KOH pH 7.6, 100 mM potassium glutamate, 10 mM $Mg(OAc)_2$, 0.05% NP-40-S, 1 mM DTT), 5 pmol $_{Sc}$Cdc45, and 10 pmol $_{Sc}$Mcm10 were incubated on ice for 20 min. Reactions were added to 5 µl Anti-FLAG M2 Magnetic Beads and incubated at 4°C, 1250 rpm, 30 min. Beads were washed by 3x 100 µl resuspension in interaction buffer. Proteins were eluted by incubation of the beads at 4°C, 1250 rpm, 20 min in a 20 µl suspension containing interaction buffer and 0.2 mg/ml 3xFLAG peptide. Samples were analysed on 4-12% Bis-Tris SDS-PAGE gels and immunoblotted with αFLAG-HRP (A8592, 1/10,000) or αT7-HRP (ab19291, 1/1000).

To detect an interaction between $_{Hs}$RECQL4$^{N\text{-}term}$ and $_{Hs}$MCM10$^{230\text{-}450}$, reactions were performed as above, but 5 pmol $_{Hs}$RECQL4$^{N\text{-}term}$ was incubated with 5 µl Anti-FLAG M2 Magnetic Beads at 4°C, 1250 rpm for 15 min, before addition of 10 pmol $_{Hs}$MCM10$^{230\text{-}450}$ and incubation continued for 30 min. Samples were analysed on 4-12% Bis-Tris SDS-PAGE gels and silver-stained. All pull-downs were performed in triplicate.

***Electrophoretic mobility shift assays***

To detect binding of DNA by $_{Sc}$Mcm10, 10 µl reactions containing 15 mM Tris-HCl pH 7.4, 100 mM sodium acetate, 10 mM ammonium sulfate, 10% glycerol, 50 nM oligonucleotide oJR021 (25 nt, 5'-Cy3) (Supplementary Table 3), and 150 nM $_{Sc}$Mcm10 were incubated at 23°C for 20 min. An additional 4 µl of 50% glycerol was added and samples analysed on native 10% polyacrylamide TBE gels at 150 V for 40 min. Fluorescence was detected using Amersham Typhoon imager (1.1.0.7) (Cytiva). All gel shift assays were performed in triplicate.

***Fluorescence anisotropy***

To analyse $_{Sc}$Mcm10 binding to ssDNA via fluorescence anisotropy, we used $_{Sc}$Mcm10$^{OB\ fold}$ constructs (a.a. 110-377) rather than full-length proteins enabling us to test higher protein concentrations. A twofold dilution series of each construct from 20 µM to 0.7 nM was prepared and mixed 3:1 with 40 nM 5'-ATTO488N labelled 23nt oligonucleotide ETC_oligo_206 (Supplementary Table 3). All reactions were performed in reaction buffer (25 mM HEPES-KOH pH 7.6, 100 mM potassium glutamate, 10 mM $Mg(OAc)_2$, 0.02% NP-40-S, 0.5 mM TCEP). Reactions were carried out in a total volume of 40 µl and incubated for 30 min at 25°C in black, flat-bottom, non-binding surface 384-well plates (Corning) before measurements. Measurements were performed with a PheraStar FSX plate reader (BMG Labtech) using an optic module for $\lambda_{ex}$ = 485 nm, $\lambda_{em}$ = 520 nm. All experiments were performed as technical triplicates and data analysed using PRISM 10 (GraphPad Software). The data is presented as mean values ± SD derived from three independent measurements. The binding model used to fit the DNA binding data was as follows: nonlinear regression (curve fit) using a one-site binding model with ligand depletion. The observed anisotropy was fit as:

$$F = F_0 + (F_1 - F_0)\frac{([P_T] + [L_T] + K_d) - \sqrt{([P_T] + [L_T] + K_d)^2 - 4[P_T][L_T]}}{2[L_T]}$$

where $F$ is the measured anisotropy, $F_0$ and $F_1$ are the anisotropy values of free and fully bound DNA, $P_T$ is total protein concentration, $L_T$ is total fluorescent DNA concentration, and $K_d$ is the equilibrium dissociation constant.

***S. cerevisiae tetrad dissection***

Details of yeast strains can be found in Supplementary Table 2. Diploid yeast cells were patched onto sporulation plates (0.25% yeast extract; 0.1% glucose; 1.5% potassium acetate; 2% agar; 5 µg/ml Arginine, 10 µg/ml Adenine , 10 µg/ml Uracil; 5 µg/ml Histidine; 5 µg/ml Leucine; 5 µg/ml Lysine; 5 µg/ml Tryptophan, 2 µg/ml Tyrosine; 25 µg/ml Phenylalanine; 5 µg/ml Methionine; 1 µg/ml Proline) and incubated for 3-5 days at 30°C. Sporulated cells were picked using sterile toothpicks and resuspended in 50 µl sterile Milli-Q water and 2 µl

Lyticase (Sigma #L2524). Following incubation at room temperature for 5 min, 1 ml sterile Milli-Q water was added and 80 µl of this mix was streaked onto YPD plates (1.1% yeast extract; 2.2% peptone; 2% glucose; 55 µg/ml Adenine; 2.5% agar). Tetrads were dissected using a micromanipulator (Singer Instruments) and the resulting cells were grown for 2 days at 30°C. Plates were imaged on an Epson Perfection V850 Pro scanner. Cells were genotyped by analysing growth on appropriate selective plates.

***Glycerol gradients to determine dimeric $_{Hs}$CMG formation***

To assess formation of $_{Hs}$CMG dimers in the presence of $_{Hs}$DONSON, a 31 µl reaction containing 25 mM HEPES-NaOH pH7.6, 20 mM NaOAc, 7.5 mM $Mg(OAc)_2$, 0.5 mM TCEP, 0.1 mM AMP-PNP, 200 nM $_{Hs}$CMG, 100 nM $_{Hs}$DONSON (dimeric concentration) was incubated on ice for 20 min. The reaction was split and applied to 200 µl 10-30% glycerol gradients (40 mM HEPES-NaOH pH7.6, 150 mM NaOAc), either in the absence or presence of glutaraldehyde crosslinker as described below. The samples were separated by centrifugation at 200,000 g, 4°C, 60 min and 12 µl fractions manually collected. The fractions from the gradient in the absence of crosslinker were analysed on 4-12% Bis-Tris SDS-PAGE gels and silver-stained. Corresponding desired crosslinked fractions were applied to 3 nm carbon film on 400-mesh copper (Agar Scientific), which had been freshly glow-discharged at 0.39 mbar and 30 mA for 40-50 s using a PELCO easiGlow. After incubation for 1-2 min, the grid surface was washed and stained by picking up three droplets (each 10 µl) of water and two drops of 2% uranyl acetate, blotting the grid on the side between drops. The second drop of uranyl acetate was incubated for 1 min and blotted and grids were dried overnight. Negative-stain samples were imaged on a 120-kV FEI Tecnai Spirit equipped with a Gatan Ultrascan 1000XP detector.

***Cryo-EM sample and grid preparation: $_{Sc}$CMG-Mcm10***

Fork DNA substrates were generated by mixing equal volumes of oligonucleotides DBo1 and DBo2 (Supplementary Table 3) and annealing by gradual cooling from 75°C to room temperature. Oligonucleotides were prepared in (25 mM HEPES-NaOH pH 7.5, 150 mM NaOAc, 2 mM $Mg(OAc)_2$ and 0.5 mM TCEP).

For replisome reconstitution, a reaction containing 25 mM HEPES-NaOH pH 7.6, 100 mM NaOAc, 15 mM $Mg(OAc)_2$, 0.5 mM TCEP, 0.5 mM AMP-PNP, 225 nM CMG (Ctf4-depleted) was incubated with a 1.5-fold molar excess over CMG of fork DNA on ice for 30 minutes. Next, a 4-fold molar excess over CMG of Mcm10 was added and incubated for a further 30 mins on ice. Then the reaction was diluted in 1x buffer to 200 ul total volume. 80 ul was then applied to 2 glycerol gradients in the presence of crosslinker, with the remaining 40 µl diluted 2-fold in 1x buffer and applied to a gradient without crosslinker. The CMG-Mcm10 sample in the absence of DNA was prepared identically except Mcm10 was added at 2-fold molar excess.

Density gradient ultracentrifugation and cryo-EM grid preparation was carried out as previously described. (Baretić, et al. 2020) Buffer A (40 mM HEPES-NaOH pH 7.5, 150 mM NaOAc, 0.5 mM TCEP, 10% v/v glycerol, 0.5 mM AMP-PNP and 3 mM $Mg(OAc)_2$) was layered over an equal volume of Buffer B (Buffer A containing 30% v/v glycerol; for crosslinked gradients Buffer B additionally contained 0.16% glutaraldehyde and 2 mM bis(sulfosuccinimidyl)suberate (BS3)) in 2.2 ml TLS-55 tubes (Beranek Laborgeräte), formed using a gradient-making station (Biocomp Instruments), and cooled on ice. Samples were centrifuged at 200,000 × g for 2 h at 4 °C, and 100 µl fractions were collected manually. Subsamples were analysed by 4-12% Bis-Tris SDS-PAGE and silver staining using the

SilverQuest staining kit. Desired crosslinked fractions were pooled and buffer-exchanged against 25 mM HEPES-NaOH pH 7.5, 150 mM NaOAc, 3 mM $Mg(OAc)_2$, 0.5 mM TCEP, 0.1 mM AMP-PNP and 0.005% TWEEN-20 six times in a 200ul concentrator, each time at 21,000 × g for 1 min per round at 4 °C. Samples were finally concentrated via a 7 minute centrifugation step to the minimum volume and 3 µl applied QUANTIFOIL R2/2 grids coated with ~2 nm continuous carbon.

***Cryo-EM data collection: $_{Sc}$CMG-Mcm10***

Dataset 1 and Dataset 2 were collected on a 300 keV Titan Krios microscope (FEI) equipped with a K3 direct electron detector (Gatan) operated in electron-counting mode, using a BioQuantum energy filter (Gatan) with a 20 eV slit width (Supplementary Table 5). Data were acquired in super-resolution mode and binned twofold, yielding an effective pixel size of 0.73 Å $px^{-1}$ at a nominal magnification of 105,000×. Images were recorded over defocus ranges of -1.2 to -3.4 µm (Dataset 1) and -1.0 to -3.5 µm (Dataset 2). Movies were dose-fractionated into 39 frames over a 4 s exposure, corresponding to total doses of 61.74 $e^-$ $Å^{-2}$ (Dataset 1) and 40.14 $e^-$ $Å^{-2}$ (Dataset 2).

***Cryo-EM data processing: $_{Sc}$CMG-Mcm10***

*Mcm10-CMG-Fork DNA*

Motion correction and CTF estimation were performed on 1,904 movies in cryoSPARC v4 (Punjani et al. 2017). Particles were initially selected using reference-free blob picking (431,291 particles) and extracted (343,406 particles; box size 480 pixels at 0.73 Å/pix). Extracted particles were down-sampled to a 120-pixel box (2.92 Å/pix) for initial processing. Ab-initio reconstruction was used to generate reference volumes for two sequential rounds of heterogeneous refinement, enabling removal of poorly aligned particles and non-CMG classes. The resulting particle subset was subjected to homogeneous refinement, yielding a reconstruction to the Nyquist 4.83 Å resolution from 276,092 particles.

To resolve conformational heterogeneity, global three-dimensional (3D) variability analysis was performed using clustering with filter resolutions of 8-9 Å. 154,711 particles belonging to clusters displaying density for Mcm10 and ssDNA were selected and subjected to a second homogeneous refinement, yielding a reconstruction at 4.83 Å resolution. A further round of global 3D variability analysis (Punjani and Fleet 2021) was then performed using clustering with a filter resolution of 8-9 Å to enrich for particles displaying density corresponding to Mcm10 bound at the Mcm2 and Mcm6 interfaces.

59,400 particles displaying full Mcm10 occupancy were reverted to their non-binned counterparts and subjected to three parallel refinement strategies. Non-uniform refinement (Ali Punjani, Haowei Zhang, and David J. Fleet 2020) of the consensus particle set yielded a reconstruction at 3.57 Å resolution (map: sc #1). Local refinements using focused masks were performed for the Mcm6, Mcm2 and Mcm5 N-tier region together with Cdc45 (3.86 Å resolution, map: sc #2) and for the Mcm2, Mcm6, Mcm4 and Mcm5 C-tier region (3.52 Å resolution, map: sc #3).

To identify particles containing Mcm6 winged-helix domain density, an additional round of focused 3D variability analysis was performed using a heavily dilated soft mask generated from a molmap reference. The reference map was constructed from an AlphaFold-3 prediction of the Mcm6 winged-helix domain bound to Mcm10 and merged with the consensus reconstruction. This strategy identified a subset of 4,650 particles, which were subjected to non-uniform refinement, yielding a reconstruction at 4.50 Å resolution (map: sc #4).

*Mcm10-CMG*

Motion correction and CTF estimation were performed in cryoSPARC v4 on 12,424 movies. Particles were initially selected using reference-free blob picking (2,155,933 particles), and incorrect picks were removed using the Inspect Picks job, yielding 1,307,294 particles. Particles were extracted (1,027,077 particles; box size 450 pixels at 0.725 Å/pix) and down-sampled to a 120-pixel box (2.719 Å/pix) for initial processing. Ab-initio reconstruction was used to generate reference volumes for heterogeneous refinement, which yielded a subset of 643,565 particles corresponding to CMG assemblies.

This subset was subjected to homogeneous refinement, yielding a reconstruction at the Nyquist limit of 5.67 Å resolution. To interrogate weak peripheral density extending from Cdc45, a soft mask encompassing the N-tier, Cdc45 and GINS, together with low-threshold density tentatively assigned to the Mcm10 OB-fold, was generated. The mask was inverted and used for signal subtraction. Subtracted particles were reconstructed without alignment and subjected to global 3D variability analysis using clustering with a filter resolution of 7 Å to identify particle classes displaying density consistent with the Mcm10 OB-fold.

Particles belonging to selected clusters were re-extracted un-binned (172,368 particles; box size 620 pixels at 0.725 Å/pix) and subjected to non-uniform refinement, yielding a consensus reconstruction at 3.58 Å resolution. Two sequential rounds of particle subtraction followed by focused local refinement enabled reconstruction of the Cdc45-Mcm10-GINS-Mcm2/5/3 N-tier region at 3.20 Å resolution (map: sc #5).

The same non-subtracted particle subset was locally refined within a mask encompassing the Mcm2 N-tier, Cdc45 and candidate Mcm10 OB-fold density, yielding a reconstruction at 3.62 Å resolution. This mask was then used for signal subtraction, and the resulting particles were subjected to 3D classification without alignment (target resolution 7 Å; PCA clustering with force-hard classification) to improve density assigned as the Mcm10 OB-fold. A subset of 78,220 particles was selected and locally refined to 3.78 Å resolution. Reversion to non-subtracted particles followed by non-uniform refinement yielded a consensus reconstruction at 3.54 Å resolution. A soft mask encompassing the Mcm2-7 C-tier was generated and used for two additional rounds of particle subtraction, followed by local refinement to generate an Mcm2-7 C-tier map at 5.76 Å resolution (map: sc #6).

Further 3D classification of the 78,220-particle subset using the same parameters resolved two conformational states of the Mcm10 OB-fold (conformations I and II). A subsequent round of 3D classification yielded final subsets of 23,013 particles for conformation I and 22,079 particles for conformation II, which were locally refined to 5.37 Å (map: sc #7) and 5.98 Å (map: sc #9) resolution respectively. These particles were then reverted to non-subtracted particles and subjected to non-uniform refinement, producing consensus reconstructions at 4.17 Å for conformation I (map: sc #8) and 4.41 Å for conformation II (map: sc #10). A summary of relevant statistics for maps sc #1-10 can be found in Supplementary Table 6.

***Molecular modeling: $_{sc}$Mcm10-$_{sc}$CMG complexes***

Mcm10-CMG-DNA complexes were modelled by first docking in CMG from a previously solved *S. cerevisiae* replisome structure (PDB 6SKL) (Baretić, et al. 2020). Ctf4 and Csm3-Tof1 were removed prior to docking. The fit of the CMG-DNA model was optimised using real-space refinement in ISOLDE (Croll 2018), with secondary-structure restraints applied, using maps covering the highest-resolution regions of each model region. Regions of the model that remained poorly supported by density were then either removed or manually rebuilt in Coot (Emsley et al. 2010). Unassigned density contiguous with CMG protein chains was built with the aid of AlphaFold3 (Abramson, et al. 2024) predictions of individual CMG subunits. Remaining density that could not be accounted for by CMG was attributed to Mcm10. AlphaFold3-predicted Mcm10-CMG interaction models were then rigid-body docked into these unmodelled regions, and residues lacking density support were pruned. Local real-space refinement was performed in ISOLDE, followed by global real-space refinement in Phenix against the consensus locally refined map (*sc* map #1).

An analogous strategy was used to model Mcm10-CMG complexes in the absence of DNA. The Mcm2-7 C-tier was initially modelled using PDB 6SKO as a starting template. Mcm10-Cdc45 and -GINS contacts were incorporated based on AlphaFold3 predictions prior to local refinement in ISOLDE. AlphaFold3 models of the Mcm10 OB-ZnF domain (residues 137-325) were rigid-body docked into cryo-EM density corresponding to conformations I and II (*sc* maps #7-10) before heavily constrained refinement in ISOLDE. Independent models of the entire complex were generated for each OB-ZnF conformation and refined by iterative local real-space refinement in ISOLDE, followed by global refinement in Phenix against consensus maps.

***Cryo-EM sample and grid preparation: $_{Hs}$MCM10/RECQL4 bound DONSON:CMG:Pol ε***

Reconstitution reactions were set up to yield a final concentration of 300 nM $_{Hs}$CMG and a 1:1 molar ratio with Pol ε, DONSON, $_{Hs}$RECQL4$^{Nterm}$ and $_{Hs}$MCM10$^{\Delta Nterm}$ in reconstitution buffer (25 mM HEPES-NaOH pH 7.5, 80 mM NaOAc, 5 mM MgOAc, 0.5 mM TCEP, 200 mM NaCl). $_{Hs}$CMG was incubated on ice with Pol ε and DONSON for 10 min. $_{Hs}$RECQL4$^{Nterm}$ and $_{Hs}$MCM10$^{\Delta Nterm}$ were added and the reaction adjusted to 20 µl using reconstitution buffer. The reaction was incubated on ice for 90 min before loading onto a 10-30% glycerol gradient.

Glycerol gradients were prepared by layering, from bottom to top, 40 µl volumes of buffers A-E containing successively decreasing concentrations of glycerol, glutaraldehyde and bis(sulfosuccinimidyl)suberate (BS3, Thermo Fisher Scientific)). Successive layers increase incrementally by 5% glycerol, 0.04% glutaraldehyde, and 0.5 mM BS3, such that buffer A (top layer) contains 10% glycerol, 0% glutaraldehyde, and 0 mM BS3, and buffer E (bottom layer) contains 30% glycerol, 0.16% glutaraldehyde, and 2.0 mM BS3. All buffers contain 40 mM HEPES-NaOH (pH 7.6) and 80 mM NaOAc. Gradients were sedimented by ultracentrifugation (259,000g, 4°C, 1 h) and then manually separated into fractions of 12 µL. Peak fractions containing dimeric complex were identified using silver-stained SDS-PAGE of samples +/− crosslinker and used for cryo-EM grid preparation.

Grids were prepared for cryo-EM using a manual plunger. 4 uL of the sample from peak fractions of the glycerol gradient were applied to freshly prepared graphene oxide grids, incubated for 1 minute at 4°C, quickly blotted away (<0.5s blotting time), followed by three more rounds of sample application and blotting. After the fourth sample application and incubation, 5 uL of reconstitution buffer (25 mM HEPES-NaOH, 80 mM NaOAc, 5 mM MgOAc, 0.5 mM TCEP, 200 mM NaCl) was applied to the grid and immediately blotted away (<0.5s blotting time), followed by 2 more rounds of buffer application and blotting. After the third buffer application, excess buffer was blotted away for 7 s and grids were plunged into liquid ethane.

Graphene oxide supports were prepared by applying 3 µl of 0.2 mg/ml graphene oxide suspension (Sigma Aldrich 763705) to freshly glow-discharged (5 min at 30-40 mA) UltrAuFoil 300 R1.2/1.3 grids. Excess liquid was blotted away after 1 min and grids were washed three times with 20 µl water, twice on the top face and once on the bottom face of the grid.

***Cryo-EM data collection: $_{Hs}$MCM10/RECQL4 bound DONSON:CMG:Pol ε***

For the $_{Hs}$MCM10- and $_{Hs}$RECQL4-bound dimeric DONSON:CMG:Pol ε complex, 51,631 movies were collected in a single session on a Titan Krios microscope (FEI) operating at 300 kV equipped with a K3 direct electron detector (Gatan) and BioQuantum energy filter (Gatan). EPU software (Thermo Fisher Scientific) with "faster acquisition" (AFIS) enabled was used for automatic acquisition of movies. A slit width of 20 eV was used for the BioQuantum energy filter and data was collected in super-resolution mode bin 2 at an effective pixel size of 0.725 Å/pixel over a defocus range of -2.6 to -0.6 µm. Movies were dose-fractionated into 40 fractions over a 1.05 s exposure, resulting in a total dose of 40.5 e-/Å$^2$ (Supplementary Table 5).

***Cryo-EM data processing: $_{Hs}$MCM10/RECQL4 bound DONSON:CMG:Pol ε***

The data processing pipeline is schematised in Supplementary Fig. 9, 11 and 12. An initial subset of 1000 movies was used to generate a gain reference using RELION 4.0's gain estimation tool (Scheres 2012; Zheng et al. 2017). Apart from gain estimation, all other processing steps were performed with CryoSPARC v4.7.1 (henceforth referred to as "CryoSPARC") (Punjani, et al. 2017).

*Pre-processing*

The pre-processing pipeline is schematised in Supplementary Fig. 9C. Gain-corrected movies were dose-weighted and motion-corrected using CryoSPARC's patch motion job. Corrected micrographs were subjected to patch CTF estimation. Micrographs of poor quality were excluded from further processing. 6.5 million particles were blob-picked from the remaining 38,719 micrographs and after 2D classification were subjected to one round of heterogenous refinement. Templates were generated from the best class, and 3.5 million particles were re-picked with the template-picking. Particles were subjected to iterative rounds of heterogeneous refinement until a subset of 333,842 particles were obtained. After one round of Non-Uniform Refinement (A. Punjani, H. Zhang, and D. J. Fleet 2020), particles with a per-particle scale lower than 0.75 were discarded and the final subset of 289,479 particles (hereafter referred to as "best" particles) that was used for all further processing steps.

*Consensus reconstruction (Map Hs1)*

The processing pipeline leading to the consensus reconstruction is schematised in Supplementary Fig. 9C and 9D. Four ab-initio reconstructions were generated from the 289,479 best particles, and the particles comprising the reconstruction with the best definition for both copies of the dimer were subjected to iterative rounds of heterogeneous refinement until three distinct classes were achieved: Pol ε bound at both $_{Hs}$CMG copies, Pol ε bound at only one copy, or classes without signal for Pol ε non-catalytic domain. These were used as input volumes for renewed heterogenous refinement of all 289,479 best particles. Particles from the best class were subjected to further heterogeneous refinement. In all rounds of iterative heterogeneous refinement, classes with best definition for both $_{Hs}$CMG copies were selected. After combining the two subsets of particles and removing duplicates,

37,944 particles were obtained. These were subjected to non-uniform refinement, followed by 3D-Flex training and reconstruction to generate the final consensus reconstruction (Map Hs1, Supplementary Fig. 9D) (Punjani and Fleet 2023). The reported resolution was calculated based on the FSC = 0.143 criterion.

*MCM N-tiers and DONSON (Map Hs2)*
Particle subtraction was performed on the subset of 37,944 particles used for the consensus reconstruction using a mask inverted around the MCM N-tiers and DONSON. The subtracted particles were then locally refined to obtain the final map of the MCM N-tiers and DONSON (Supplementary Fig. 9E). The reported resolution was calculated based on the FSC = 0.143 criterion.

*DONSON (Map Hs3)*
The 289,479 best particles were subjected to five rounds of heterogenous refinement. In each round, the class with the best definition for DONSON was selected for further refinement. This resulted in a subset of 108,432 particles, which was combined with the 37,944 particles comprising the consensus reconstruction. After duplicates were removed, a final subset of 111,222 particles was obtained. Particle subtraction was performed on this subset using a mask inverted around DONSON. The subtracted particles were locally refined around DONSON to obtain the final DONSON map. (Supplementary Fig. 9F). The reported resolution was calculated based on the FSC = 0.143 criterion.

*Local refinements of $_{Hs}$MCM10, RECQL4 and DONSON interactions with the CMG:Pol ε dimer (maps Hs4-10)*
The 289,479 best particles were symmetry-expanded (C2) and locally refined around one $_{Hs}$CMG:Pol ε copy (Supplementary Fig. 11A). The resulting 578,958 particles were subjected to 3d-classification with a mask around the MCM C-tiers and Pol ε, and a total of 360,095 particles with Pol ε bound to $_{Hs}$CMG were subclassified. The total symmetry-expanded CMG particles (578,958) and Pol ε-bound $_{Hs}$CMG particles (360,095) were used for further processing as outlined below.

The processing pipeline for the RECQL4-CDC45 and DONSON-SLD5 interactions is shown as a schematic in Supplementary Fig. 11B. Symmetry-expanded $_{Hs}$CMG particles were locally refined around CDC45-GINS. The particles were then subtracted with a mask inverted around CDC45-GINS and then subjected to another round of local refinement to obtain the final map showing RECQL4-CDC45 and DONSON-SLD5 interactions (Map Hs4). The reported resolution was calculated based on the FSC = 0.143 criterion and the map colored by local resolution is shown in Supplementary Fig. 11E.

The processing pipeline for the RECQL4-POLE1 and DONSON-POLE2 interactions is shown as a schematic in Supplementary Fig. 11C. Symmetry-expanded Pol ε-bound $_{Hs}$CMG particles were re-centred on Pol ε, re-extracted, and subjected to particle subtraction with a mask inverted around Pol ε. Subtracted particles were subclassified with a mask around RECQL4 and particles with clear density for RECQL4 were pooled and reconstructed to obtain the final map showing RECQL4-POLE1 and DONSON POLE2 (Map Hs5). The reported resolution was calculated based on the FSC = 0.143 criterion and the map colored by local resolution is shown in Supplementary Fig. 11F.

The processing pipeline for the MCM10 OB-fold is shown as a schematic in Supplementary Fig. 11D. Symmetry-expanded $_{Hs}$CMG particles were downsampled to a 448 px box (1.24 Å/pix) and the signal for the second copy of the dimeric complex was removed by particle subtraction after local refinement around the second copy. Subtracted particles were locally refined with a mask around the N-tier domains of MCM6-4-7 and OBZnF of MCM10 and subsequently recentered to the center of the same mask. Particles were cropped to a 384 px box and downsampled to a 256 px box (1.86 Å/px). After 3d classification with a mask around the OBZnF domain of MCM10, particles with the best signal for MCM10 OB were selected, refined as above and subjected to 3D variability analysis in cluster mode. Particle clusters with the best density for MCM10 OB were selected, cropped to a 144 px box and used for homogenous 3D reconstruction. The resulting unsharpened map was filtered with a gaussian filter (1.5 Å standard deviation) inside ChimeraX. The reported resolution was calculated based on the FSC = 0.143 criterion and the map colored by local resolution is shown in Supplementary Fig. 11G.

The processing pipeline for the MCM10-MCM6 interaction is shown as a schematic in Supplementary Fig. 12A. Symmetry-expanded Pol ε-bound $_{Hs}$CMG particles were locally refined with a mask around the MCM C-tier and Pol ε. Particles were recentred around the refinement mask and re-extracted, then subjected to 3d classification using a mask around the MCM2,6,4 C-tiers. MCM10-bound particles were pooled and locally refined with a mask around the MCM C-tier, CDC45-GINS, and Pol ε. Particles were locally and globally CTF-refined, then subject to a final round of local refinement to obtain the final map showing the MCM10-MCM6 interaction (Map Hs6). The reported resolution was calculated based on the FSC = 0.143 criterion and the map colored by local resolution is shown in Supplementary Fig. 12E.

The processing pipeline for the MCM10-MCM2 interaction is shown as a schematic in Supplementary Fig. 12B. Symmetry-expanded $_{Hs}$CMG particles were locally refined around the MCM2 and MCM6 N-tier and CDC45-GINS, then subjected to 3d classification using a mask around the MCM2 N-tier. MCM10-bound particles were pooled and underwent particle subtraction with a mask inverted around the MCM N-tier and CDC45-GINS. Particles were cropped and locally refined using a mask around CDC45-GINS and the N-tier of MCM2 and MCM6. Particle subtraction and local refinement were repeated once more to obtain the final map showing the MCM10-MCM2 interaction (Map Hs7). The reported resolution was calculated based on the FSC = 0.143 criterion and the map colored by local resolution is shown in Supplementary Fig. 12E.

The processing pipeline for the MCM10-MCM4 interaction and the MCM6 NtHp are shown as a schematic in Supplementary Fig. 12C. Symmetry-expanded $_{Hs}$CMG particles were locally refined around the MCM N-tier and then subjected to particle subtraction using a mask inverted around the MCM N-tier. The subtracted particles were cropped and then underwent 3d classification with a mask around the MCM4 N-tier. MCM10-bound classes were pooled and locally refined around the MCM2-6-4 N-tiers and MCM10 ZnF, and after one round of local and global CTF refinement and local refinement, the map showing the interaction between MCM10-MCM4 was obtained (Map Hs8). The cropped particles were also subjected to 3d classification using a mask around the MCM6 NtHp. Classes showing clear density for the insert were pooled and combined with classes from the 3d classification with the MCM4 mask which also showed density for the same insert. After removing duplicates, the remaining particles were locally refined to obtain the final map showing the MCM6 NtHp interacting with MCM4 and MCM7 (Map Hs9). The reported resolution was calculated based on the FSC = 0.143 criterion and the map colored by local resolution is shown in Supplementary Fig. 12E.

The composite cryo-EM map shown in Figure 5A was obtained by docking map Hs3 and two respective copies of maps Hs4-10 into the composite map Hs1. Maps are displayed around their corresponding region of the model at contour levels (σ-thresholds) listed in Supplementary Table 7.

#### *Molecular modeling: $_{Hs}$MCM10/RECQL4-bound DONSON:CMG:Pol ε*

An initial atomic model was obtained by docking two copies of $_{Hs}$CMG:Pol ε, obtained from 7PFO (Jones EMBO), into the consensus map Hs1 using UCSF Chimera (Pettersen et al. 2004). A model for dimeric HsDONSON was generated by Alphafold3 (Abramson, et al. 2024) and docked into the consensus map. Models for small interacting motifs between DONSON-SLD5, DONSON-POLE2, MCM10 and MCMs, RECQL4-POLE1 and RECQL4-CDC45 were obtained by Alphafold3 and from predictomes (Schmid and Walter 2025). The fit-to-density and model geometry were subsequently refined using Coot (Emsley, et al. 2010) ISOLDE inside ChimeraX (Croll 2018) using individual map with the best local resolution for the refined region. The resulting model was refined using phenix.real_space_refine (Afonine et al. 2018) against the consensus map with global minimization enabled. Model validation was carried out using MolProbity (Chen et al. 2010) inside Phenix after a single round of atomic displacement parameter refinement and is summarised in Tables S5 and S7.

#### *Human CMG apo structure*

$_{Hs}$CMG was dialysed against 40 mM HEPES-KOH pH 7.6, 80 mM KOAc, 0.25 mM EDTA, 1 mM DTT, 0.005 % NP-40 and applied to freshly glow discharged (4.5 min, 30 mA) UltrAuFoil 300 1.2/1.3. Excess sample was blotted away for 2 s and grids were manually plunged into liquid ethane. 1,108 movies were collected on a Titan Krios microscope (FEI) operating at 300 kV equipped with a K3 direct electron detector (Gatan) and BioQuantum energy filter (Gatan). EPU software (Thermo Fisher Scientific) with “faster acquisition” (AFIS) enabled was used for automatic acquisition of movies. A slit width of 20 eV was used for the BioQuantum energy filter and data was collected in super-resolution mode bin 2 at an effective pixel size of 0.725 Å/pixel over a defocus range of -3.0 to -1.0 µm. Movies were dose-fractionated into 40 fractions over a 1.35 s exposure (13.6 e-/px/s flux), resulting in a total dose of 35 e-/Å2 (Supplementary Table 5). The data processing pipeline is schematised in Supplementary Fig. 13. Movies were dose-weighted and motion-corrected using RELION v4, imported into CryoSPARC v4 and subjected to patch CTF estimation. Particles were picked using template-based picker and extracted in a 512 px box, Fourier-cropped to 128 px (2.900 Å/pix). After 2D classification, initial 3D references were generated by ab-initio reconstruction from particles subsets belonging to either 2D classes with CMG features or ‘junk’-classes (Supplementary Fig. 13C) and then provided for Heterogenous Refinement. CMG particles were re-extracted (512 px box, Fourier-cropped to 384 px, 0.967 Å/pix). Two separate strategies were applied to yield a map of apo-CMG (Hs11) and a separate, locally refined map with a focus around the N-tier of MCM6-MCM4-MCM7 (Hs12). For Hs11, particles were subjected to 3D classification with a focus mask around the MCM C-tier, and the class with the best definition for the C-tier was selected. Particles were used for Homogenous Refinement to generate a consensus map (Hs11a) and for local refinement to generate a focus map of the MCM C-tier (Hs11b). The two maps were filtered based on local resolution estimates and combined using combine_focused_maps inside Phenix. For Hs12, particles with a relative per-particle scale below 0.88 were removed and the remaining particles were locally refined with a focus mask around the region of MCM4, MCM6 and MCM7 of the MCM N-tier. The resulting map was filtered based on local resolution estimates. Reported models for CMG (PDB: 6XTX (Rzechorzek et al. 2020), PDB: 7PFO (Jones, et al. 2021),) and MCM4 N-terminal region in the context of MCM double hexamer (8W0F (Yang, et al. 2024)) were used to generate an initial atomic model for CMG. The fit-to-density and model geometry were subsequently refined using Coot and ISOLDE inside ChimeraX. The resulting model was refined using phenix.real_space_refine against the composite map Hs11 with global minimization enabled. Model validation was carried out using MolProbity inside Phenix after a single round of atomic displacement parameter refinement and is summarised in Tables S5 and S7.

***Statistics and Reproducibility***

Experiments in Fig. 2H, 4G and Supplementary Fig. 5D, F, I, 6F-H, 7I, 8B, D were performed a minimum of three times. Experiments in Fig. 3E and Supplementary Fig. 5G, H were performed twice. $_{Sc}$Mcm10 mutants in Fig. 2F, 3D were tested in mini-circle assays twice. $_{Sc}$Mcm10 mutants in Fig. 2C, 3B, 4F and Supplementary Fig. 7G were tested in mini-circle assays a minimum of three times, with the exception of $_{Sc}$Mcm10$^{CR}$ in Fig. 4F which was tested once. Yeast tetrad dissections in Fig. 4H, I and Supplementary Fig. 5J, K, S7J, K were each performed three times for two independent clones. Negative stain-EM and cryo-EM sample preparation was performed once for each complex studied, but *in vitro* formation of all described protein complexes was performed a minimum of two times.

| Name | Characteristics | Usage | Reference |
|---|---|---|---|
| 1883 | pet28a/6xHis-Mcm10 | Construction of $_{Sc}$Mcm10 mutants. | Yeeles et al. 2015 |
| pJR01 | pet28a/2xStrep-TEV-MCM10$^{Δ1-229}$-TEV-2xFLAG | Purification of $_{Hs}$MCM10$^{ΔNterm}$ | This study |
| pJR02 | pet28a/2xStrep-TEV-MCM10$^{230-450}$ | Purification of $_{Hs}$MCM10$^{OBZnF}$ | This study |
| vEF32 | pRS303/2xFLAG-TEV-hDONSON | Purification of $_{Hs}$DONSON from *S. cerevisiae* | This study |
| vETC100 | pFA6a/*mcm10-5A-URA3* | Replacement of W303 *S. cerevisiae MCM10* with *mcm10-5A-URA3* allele | This study |
| vETC13 | pet28a/6xHis-Mcm10$^{F113A, K116A, F117A, K121A, K122A, E124A}$ | Purification of $_{Sc}$Mcm10$^{7A}$ | This study |
| vETC15 | pet28a/6xHis-Mcm10$^{F113A, F117A}$ | Purification of $_{Sc}$Mcm10$^{2A}$ | This study |
| vETC27 | pet28a/6xHis -Mcm10$^{D333A, E337A}$ | Purification of $_{Sc}$Mcm10$^{CR}$ | This study |
| vETC48 | pet28a/6xHis-Mcm10$^{R347A, E49A, L350A}$ | Purification of $_{Sc}$Mcm10$^{3A}$ | This study |
| vETC68 | pFA6a/*mcm10-3A-URA3* | Replacement of W303 *S. cerevisiae MCM10* with *mcm10-3A-URA3* allele | This study |
| vETC69 | pFA6a/*mcm10-2A-URA3* | Replacement of W303 *S. cerevisiae MCM10* with *mcm10-2A-URA3* allele | This study |
| vETC70 | pFA6a/*mcm10*$^{Y245A, F247A, F230A}$-*URA3* | Replacement of W303 *S. cerevisiae MCM10* with *mcm10*$^{Y245A, F247A, F230A}$-*URA3* allele | This study |
| vETC86 | pet28a/6xHis-Mcm10$^{N313E, K314E, K315A, W274A, L272A, H215E, K220E}$ | Purification of $_{Sc}$Mcm10$^{DNA}$ | This study |
| vETC87 | pet28a/Mcm10$^{110-377}$-StrepII | Purification of $_{Sc}$Mcm10 OBZnF | This study |
| vETC88 | pFA6a/*mcm10-DNA-URA3* | Replacement of W303 *S. cerevisiae MCM10* with *mcm10-DNA-URA3* allele | This study |
| vETC89 | pet28a/Mcm10$^{110-377\ (N313E, K314E, K315A, W274A, L272A, H215E, K220E)}$-StrepII | Purification of $_{Sc}$Mcm10 OBZnF$^{DNA}$ | This study |
| vETC90 | pet28a/6xHis-Mcm10$^{F401A, F410A, F411A}$ | Purification of $_{Sc}$Mcm10$^{3FA}$ | This study |
| vETC92 | pet28a/ALFA-Mcm10$^{Δ388-571}$ | Purification of $_{Sc}$Mcm10$^{N-term}$ | This study |
| vETC99 | pet28a/ALFA-Mcm10$^{107-387}$ | Purification of $_{Sc}$Mcm10$^{N-termΔN}$ | This study |
| vJY216 | pet28a/RECQL4$^{1-440}$-2xFLAG | Purification of $_{Hs}$RECQL4$^{Nterm}$ | This study |
| vJY233 | pet28a/Mcm10$^{Δ2-387}$-ALFA | Purification of $_{Sc}$Mcm10$^{C-term}$ | This study |
| vJY236 | pet28a/6xHis-Mcm10$^{M134A, R138A, V139A, F142A}$ | Purification of $_{Sc}$Mcm10$^{pin1}$ | This study |
| vJY239 | pet28a/6xHis-Mcm10$^{P202A, Y204A, N206D}$ | Purification of $_{Sc}$Mcm10$^{pin2}$ | This study |

**Supplementary Table 1: Plasmids used in this study.**

| Strain name | Genotype | Reference |
|---|---|---|
| yEF32<br>($_{Hs}$DONSON<br>purification) | *MATa ade2-1 ura3-1 his3-11,15 trp1-1 leu2-3,112*<br>*can1-100*<br>*bar1::Hyg*<br>*pep4::KanMX*<br>*his3::HIS3 pRS303/2xFLAG-TEV-hDONSON* | This study |
| yETC44 | *MATa ade2-1 ura3-1 his3-11,15 trp1-1 leu2-3,112*<br>*can1-100*<br>*MCM10 / mcm10-2A (URA3)* | This study |
| yETC48 | *MATα ade2-1 ura3-1 his3-11,15 trp1-1 leu2-3,112*<br>*can1-100*<br>*MCM10 / mcm10-3A (URA3)* | This study |
| yETC49 | *MATα ade2-1 ura3-1 his3-11,15 trp1-1 leu2-3,112*<br>*can1-100*<br>*MCM10 / mcm10-3A (URA3)*<br>*SML1 / sml1Δ::HIS3*<br>*MEC1 / mec1Δ::ADE2* | This study |
| yETC51 | *MATa ade2-1 ura3-1 his3-11,15 trp1-1 leu2-3,112*<br>*can1-100*<br>*MATα ade2-1 ura3-1 his3-11,15 trp1-1 leu2-3,112*<br>*can1-100*<br>*MCM10 / mcm10-3A (URA3)*<br>*SML1 / sml1Δ::HIS3*<br>*MEC1 / mec1Δ::ADE2* | This study |
| yETC55 | *MATa ade2-1 ura3-1 his3-11,15 trp1-1 leu2-3,112*<br>*can1-100*<br>*MATα ade2-1 ura3-1 his3-11,15 trp1-1 leu2-3,112*<br>*can1-100*<br>*MCM10 / mcm10-2A (URA3)* | This study |
| yETC64 | *MATa ade2-1 ura3-1 his3-11,15 trp1-1 leu2-3,112*<br>*can1-100*<br>*MCM10 / mcm10-DNA (URA3)* | This study |
| yETC66 | *MATa ade2-1 ura3-1 his3-11,15 trp1-1 leu2-3,112*<br>*can1-100*<br>*MATα ade2-1 ura3-1 his3-11,15 trp1-1 leu2-3,112*<br>*can1-100*<br>*MCM10 / mcm10-DNA (URA3)* | This study |
| yETC69 | *MATa ade2-1 ura3-1 his3-11,15 trp1-1 leu2-3,112*<br>*can1-100*<br>*MATα ade2-1 ura3-1 his3-11,15 trp1-1 leu2-3,112*<br>*can1-100*<br>*MCM10 / mcm10-DNA (URA3)*<br>*SML1 / sml1Δ::HIS3*<br>*MEC1 / mec1Δ::ADE2* | This study |
| yETC70 | *MATa ade2-1 ura3-1 his3-11,15 trp1-1 leu2-3,112*<br>*can1-100*<br>*MCM10 / mcm10-DNA (URA3)*<br>*SML1 / sml1Δ::HIS3*<br>*MEC1 / mec1Δ::ADE2* | This study |
| yETC71 | *MATa ade2-1 ura3-1 his3-11,15 trp1-1 leu2-3,112*<br>*can1-100*<br>*MATα ade2-1 ura3-1 his3-11,15 trp1-1 leu2-3,112*<br>*can1-100*<br>*MCM10 / mcm10-5A (URA3)* | This study |
| yETC73 | *MATa ade2-1 ura3-1 his3-11,15 trp1-1 leu2-3,112*<br>*can1-100* | This study |

| | | |
|---|---|---|
| | *MATα ade2-1 ura3-1 his3-11,15 trp1-1 leu2-3,112 can1-100*<br>*MCM10 / mcm10$^{Y245A, F247A, F230A}$ (URA3)* | |
| yETC74 | *MATa ade2-1 ura3-1 his3-11,15 trp1-1 leu2-3,112 can1-100*<br>*MCM10 / mcm10$^{Y245A, F247A, F230A}$ (URA3)* | This study |
| yETC75 | *MATa ade2-1 ura3-1 his3-11,15 trp1-1 leu2-3,112 can1-100*<br>*MATα ade2-1 ura3-1 his3-11,15 trp1-1 leu2-3,112 can1-100*<br>*MCM10 / mcm10$^{Y245A, F247A, F230A}$ (URA3)*<br>*SML1 / sml1Δ::HIS3*<br>*MEC1 / mec1Δ::ADE2* | This study |
| yETC76 | *MATα ade2-1 ura3-1 his3-11,15 trp1-1 leu2-3,112 can1-100*<br>*MCM10 / mcm10$^{Y245A, F247A, F230A}$ (URA3)*<br>*SML1 / sml1Δ::HIS3*<br>*MEC1 / mec1Δ::ADE2* | This study |
| yETC77 | *MATa ade2-1 ura3-1 his3-11,15 trp1-1 leu2-3,112 can1-100*<br>*MATα ade2-1 ura3-1 his3-11,15 trp1-1 leu2-3,112 can1-100*<br>*MCM10 / mcm10-3A (URA3)* | This study |

**Supplementary Table 2: *S. cerevisiae* strains used in this study.**

| Name | Usage | Sequence | Reference |
|---|---|---|---|
| DBo1 | Cryo-EM (lagging strand) | 5′-GGCAGGCAGGCAGGCACACAC<br><br>TCTCCAATTCTCTAATCACTTACCA<br><br>(Biotinylated-dT)CACTTCCTACTCTA-3′ | Baretic et al. 2020 |
| DBo2 | Cryo-EM (leading strand) | 5′-(Cy3)TAGAGTAGGAAGTGA (Biotinylated dT)GGTAAGTGAT TAGAGAATTGGAGAGTGTG$(T)_{34}$ T*T*T*T*T*T-3′ *phosphorothioate | Baretic et al. 2020 |
| ETC_oligo_154 | Primer used to amplify *MCM10-URA3* with 57 bp homology overhangs for integration into the *MCM10* locus | 5′-GTTTGTCAATTCAACCTCACATTTTCA ACGCACATTAAGCACTTGGTTCGTGGAG AAATGAATGATCCTCGTGAAATT-3' | This study |
| ETC_oligo_155 | Primer used to amplify *MCM10-URA3* with 57 bp homology overhangs for integration into the *MCM10* locus | 5′-CATTGTCCATCAAAAGTAATACCATTT TGGGGCCCTGAAAAGCACACCAATACTT ATTTAGTTTTGCTGGCCGCATCTTC-3' | This study |
| ETC_oligo_173 | Primer used to amplify Mcm10 allele linked to Ura3 marker | 5′-GTATGAACAAATTTTGGGTTTATTT GCC-3' | This study |
| ETC_oligo_174 | Primer used to amplify Mcm10 allele linked to Ura3 marker | 5′-CCAACAATAATAATGTCAGA TCCTGTAG-3' | This study |
| ETC_oligo_206 | ssDNA used for fluorescence anisotropy, sequence derived from ARS1 sequence | 5′-(ATTO488N)ATGCTAAATCATTTG GCTTTTTG-3′ | This study |
| JY_OLIGO_698 | Primer for generation of mini-circles | 5′-TTCAATATAAACTAGTACGACAGGTTT(6-FAM)CCCGACTGG-3' | This study |
| JY_OLIGO_699 | Primer for generation of mini-circles | 5′-TTCAATATAAGCTAGCGAGCGGGTCTG GTGAATAGTG-3' | This study |
| oJR021 | ssDNA used for gel shift assays | 5′-(Cy3)GTGGATCCATCGCCATTGTTC CGTG-3′ | This study |

**Supplementary Table 3: Oligonucleotides used in this study.**

| Protein | Affinity tag | Purified as in | Purification steps |
|---|---|---|---|
| Cdc45 | Internal 2xFLAG tag | Yeeles et al., 2015 | Anti-FLAG M2 Affinity Gel Bio-Gel HT Hydroxyapatite |
| Cdc6 | N-terminal cleavable GST tag | Coster et al., 2014 | Glutathione Sepharose 4B Bio-Gel HT Hydroxyapatite |
| Cdt1.Mcm2-7 | N-terminal cleavable CBP tag on Mcm3 | Coster et al., 2014 | Calmodulin Sepharose 4B Superdex 200 Increase 10/300 GL |
| CMG (endogenous Ctf4 depleted) | N-terminal cleavable CBP tag on Mcm3 Internal 2xFLAG tag on Cdc45 Twin strep tag on endogenous Ctf4 | Fletcher et al., 2025 | Anti-FLAG M2 Affinity Gel Strep-Tactin XT 4Flow Calmodulin Sepharose 4B MonoQ PC 1.6/5 |
| Ctf4 | N-terminal CBP tag | Yeeles et al., 2015 | Calmodulin Sepharose 4B MonoQ 5/50 GL Superdex 200 Increase 10/300 GL |
| DDK | CBP tag on Dbf4 | On et al, 2014 | Calmodulin Sepharose 4B Lambda phosphatase dephosphorylation Superdex 200 Increase 10/300 GL |
| Dpb11 | C-terminal 3xFLAG tag | Yeeles et al., 2015 | Anti-FLAG M2 Affinity Gel MonoS 5/50 GL |
| GINS | N-terminal His tag on Psf3 | Yeeles et al., 2015 | Ni-NTA Agarose MonoQ 5/50 GL Superdex 200 Increase 10/300 GL |
| Mcm10 (+mutants) | N-terminal 6xHis tag | Yeeles et al., 2015 | Ni-NTA Agarose MonoS 5/50 GL (twice) |
| Mrc1 | C-terminal 2xFLAG tag | Yeeles et al., 2017 | Anti-Flag M2 Affinity Gel Superose 6 Increase 10/300 GL |
| ORC | Cleavable CBP tag on Orc1 | Frigola et al., 2013 | Calmodulin Sepharose 4B Superdex 200 Increase 10/300 GL |
| PCNA | Untagged | Yeeles et al., 2017 | Nucleic acid precipitation with Polymin P Ammonium sulphate precipitation HiTrap DEAE Fast Flow MonoQ 5/50 GL (twice) HiTrap Q FF Superdex 200 Increase 10/300 GL |
| Pol α-primase | N-terminal CBP tag on Pri1 | Yeeles et al., 2017 | Calmodulin Sepharose 4B MonoQ 5/50 GL Superdex 200 Increase 10/300 GL |
| Pol δ | C-terminal CBP tag on Pol32 | Yeeles et al., 2017 | Calmodulin Sepharose 4B HiTrap Heparin HP Superdex 200 Increase 10/300 GL |

| | | | |
|---|---|---|---|
| Pol ε | C-terminal CBP tag on Dpb4 (3xFLAG tag on endogenous Pol ε) | Yeeles et al., 2015 | Calmodulin Sepharose 4B (Anti-FLAG M2 Affinity Gel) HiTrap Heparin HP Superdex 200 Increase 10/300 GL |
| RFC | N-terminal CBP tag on Rfc3 | Yeeles et al., 2017 | Calmodulin Sepharose 4B MonoS 5/50 GL Superdex 200 Increase 10/300 GL |
| RPA | Untagged | Baretic et al., 2020 | HiTrap Blue HP (twice) ssDNA Cellulose MonoQ 5/50 GL |
| S-CDK (Δ1-100 Clb5) | N-terminal cleavable CBP tag on Clb5 | Hill et al, 2020 | Calmodulin Sepharose 4B Elution by TEV cleavage Superdex 200 Increase 10/300 GL |
| Sld2 | C-terminal 2xFLAG tag | Yeeles et al., 2015 With modified 2xFLAG | Ammonium sulphate precipitation Anti-FLAG M2 Affinity Gel HiTrap SP HP |
| Sld3/7 | C-terminal cleavable TCP tag | Yeeles et al., 2015 | IgG Sepharose Fast Flow TEV removal with Ni-NTA Agarose Superdex 200 Increase 10/300 GL |
| Tof1-Csm3 | N-terminal cleavable CBP tag on Csm3 | Yeeles et al., 2017 | Calmodulin Sepharose 4B MonoQ 5/50 GL Superdex 200 Increase 10/300 GL |
| TopoI | N-terminal TEV-CBP tag | Yeeles et al., 2017 Westhorpe et al. 2024 | Calmodulin Sepharose 4B TALON column Superdex 200 Increase 10/300 GL |

**Supplementary Table 4: Purification strategies for *S. cerevisiae* proteins used in this study.**

| | $_{\mathrm{Hs}}$CMG-Pol ε-DONSON-MCM10-RECQL4 (EMDB-xxxx) (PDB xxxx) | $_{\mathrm{Sc}}$CMG-Mcm10-DNA (EMDB-xxxx) (PDB xxxx) | $_{\mathrm{Sc}}$CMG-Mcm10 Conformation I (EMDB-xxxx) (PDB xxxx) | $_{\mathrm{Sc}}$CMG-Mcm10 Conformation II (EMDB-xxxx) (PDB xxxx) | Apo-$_{\mathrm{Hs}}$CMG (EMDB-xxxx) (PDB xxxx) |
|---|---|---|---|---|---|
| **Data collection and processing** | | | | | |
| Magnification | 105,000x | 105,000x | 105,000x | | 105,000x |
| Voltage (kV) | 300 | 300 | 300 | | 300 |
| Electron exposure (e–/Å$^2$) | 40.05 | 61.74 | 40.14 | | 35 |
| Defocus range (μm) | -0.6 – 2.6 | -1.2 – 3.4 | -1 – 3.5 | | -1 – 3.0 |
| Pixel size (Å) | 0.725 | 0.73 | 0.73 | | 0.725 |
| Symmetry imposed | No | No | No | | No |
| Initial particle images (no.) | 6.5 million | 343,406 | 1,027,077 | | 248,366 |
| Final particle images (no.) | See Supplementary Table 7 | See Supplementary Table 6 | See Supplementary Table 6 | | See Supplementary Table 6 |
| Map resolution (Å)<br>0.143 FSC threshold | See Supplementary Table 7 | See Supplementary Table 6 | See Supplementary Table 6 | | See Supplementary Table 6 |
| Map resolution range (Å) | 2.6-10 | 2.6-10 | 2.6-10 | | 3.0-8.0 |
| Map sharpening *B* factor (Å$^2$) | See Supplementary Table 7 | See Supplementary Table 6 | See Supplementary Table 6 | | See Supplementary Table 6 |
| **Refinement** | | | | | |
| Initial model used (PDB code) | 7PFO, 8W0F, AlphaFold3 | 6SKL AlphaFold3 | 6SKL/6SKO AlphaFold3 | 6SKL/6SKO AlphaFold3 | 7PFO, 6XTX, 8W0F |
| Model resolution (Å)<br>0.143 FSC threshold | 4.0 | 3.7 | 4.2 | 4.3 | 3.3 |
| Model composition | | | | | |
| Non-hydrogen atoms | 108,540 | 43,085 | 43,662 | 43,220 | 42,187 |
| Protein residues | 13,709 | 5,332 | 5,405 | 5,405 | 5,370 |
| Ligands | ZN: 16 | ANP: 3, ZN: 5, MG: 3 | ZN: 5 | ZN: 5 | ZN: 5 |
| *B* factors (Å$^2$) | | | | | |
| Protein | 293.77 | 82.48 | 80.92 | 80.92 | 94.31 |
| Ligand | 440.26 | 43.08 | 48.12 | 48.12 | 217.63 |
| R.m.s. deviations | | | | | |
| Bond lengths (Å) | 0.003 | 0.005 | 0.006 | 0.005 | 0.003 |
| Bond angles (°) | 0.573 | 0.711 | 0.827 | 0.753 | 0.500 |
| Validation | | | | | |
| MolProbity score | 1.74 | 0.75 | 1.11 | 1.27 | 1.11 |
| Clashscore | 6.35 | 0.30 | 0.64 | 0.81 | 2.43 |
| Poor rotamers (%) | 2.89 | 0.84 | 2.21 | 2.50 | 1.15 |
| Ramachandran plot | | | | | |
| Favored (%) | 97.83 | 97.30 | 97.30 | 96.64 | 97.82 |
| Allowed (%) | 2.17 | 2.7 | 2.66 | 3.30 | 2.18 |
| Disallowed (%) | 0.00 | 0.00 | 0.04 | 0.06 | 0.00 |

**Supplementary Table 5: Statistics of cryo-EM data collection, refinement and validation.**

| **Data collection and processing** | Final particle images (no.) | Map resolution* (Å) | Map sharpening B factor (Å2) | Model resolution* (Å) | Sigma contour level |
|---|---|---|---|---|---|
| sc#1 | 59,400 | 3.57 | -60 | 3.7 | 0.0605 |
| sc#2 | 59,400 | 3.86 | -80 | 4.0 | 0.0662 |
| sc#3 | 59,400 | 3.52 | -60 | 3.7 | 0.0822 |
| sc#4 | 4,650 | 4.50 | -100 | 5.7 | 0.0433 |
| sc#5 | 172,368 | 3.20 | -50 | 3.4 | 0.246 |
| sc#6 | 78,220 | 5.76 | -140 | 8.0 | 0.599 |
| sc#7 | 23,013 | 5.37 | -120 | 6.0 | 0.32 |
| sc#8 | 23,013 | 4.17 | -70 | 4.2 | 0.357 |
| sc#9 | 22,079 | 2.98 | -20 | 7.6 | 0.367 |
| sc#10 | 22,079 | 4.41 | -95 | 4.3 | 0.361 |

*0.143 FSC threshold

**Supplementary Table 6: Statistics summary for *S. cerevisiae* cryo-EM maps.**

| **Data collection and processing** | Final particle images (no.) | Map resolution* (Å) | Map sharpening B factor (Å2) | Model resolution* (Å) | Sigma contour level |
|---|---|---|---|---|---|
| Hs1 | 37,000 | 4.56 | -25 | 4.0 | 0.006 |
| Hs2 | 37,994 | 3.93 | -25 | 3.8 | 0.175 |
| Hs3 | 33,303 | 3.58 | -63.6 | 3.5 | 0.3 |
| Hs4 | 563,957 | 2.61 | -25 | 2.7 | 0.13 |
| Hs5 | 162,673 | 3.29 | -50 | 3.2 | 0.1 |
| Hs6 | 44,390 | 3.82 | -150 | 5.9 | 0.11 |
| Hs7 | 56,131 | 3.23 | -15 | 3.7 | 0.2 |
| Hs8 | 59,962 | 3.38 | -50 | 3.6 | 0.18 |
| Hs9 | 458,469 | 3.20 | -25 | 3.1 | 0.135 |
| Hs10 | 127,700 | 3.90 | N/A | 13.8 | 0.33 |
| Hs11a | 34,693 | 3.40 | -71.1 | 3.3 | 0.22 |
| Hs12 | 53,976 | 3.20 | -86.3 | 3.2 | 0.33 |

*0.143 FSC threshold

**Supplementary Table 7: Statistics summary for *H. sapiens* cryo-EM maps.**
